# Liver microRNA transcriptome reveals miR-182 as link between type 2 diabetes and fatty liver disease in obesity

**DOI:** 10.1101/2023.10.02.560594

**Authors:** Christin Krause, Jan H. Britsemmer, Miriam Bernecker, Anna Molenaar, Natalie Taege, Nuria Lopez-Alcantara, Cathleen Geißler, Meike Kaehler, Katharina Iben, Anna Judycka, Jonas Wagner, Stefan Wolter, Oliver Mann, Paul T. Pfluger, Ingolf Cascorbi, Hendrik Lehnert, Kerstin Stemmer, Sonja C. Schriever, Henriette Kirchner

## Abstract

**Background:** The development of obesity-associated comorbidities such as type 2 diabetes (T2D) and hepatic steatosis has been linked to selected microRNAs in individual studies; however, an unbiased genome-wide approach to map T2D induced changes in the miRNAs landscape in human liver samples, and a subsequent robust identification and validation of target genes is still missing.

**Methods:** Liver biopsies from age- and gender-matched obese individuals with (n=20) or without (n=20) T2D were used for microRNA microarray analysis. The candidate microRNA and target genes were validated in 85 human liver samples, and subsequently mechanistically characterized in hepatic cells as well as by dietary interventions and hepatic overexpression in mice.

**Results:** Here we present the human hepatic microRNA transcriptome of type 2 diabetes in liver biopsies and use a novel seed prediction tool to robustly identify microRNA target genes, which were then validated in a unique cohort of 85 human livers. Subsequent mouse studies identified a distinct signature of T2D-associated miRNAs, partly conserved in both species. Of those, human-murine miR-182-5p was the most associated to whole-body glucose homeostasis and hepatic lipid metabolism. Its target gene *LRP6* was consistently lower expressed in livers of obese T2D humans and mice as well as under conditions of miR-182-5p overexpression. Weight loss in obese mice decreased hepatic miR-182-5p and restored *Lrp6* expression and other miR-182-5p target genes. Hepatic overexpression of miR-182-5p in mice rapidly decreased LRP6 protein levels and increased liver triglycerides and fasting insulin under obesogenic conditions after only seven days.

**Conclusion:** By mapping the hepatic miRNA-transcriptome of type 2 diabetic obese subjects, validating conserved miRNAs in diet-induced mice, and establishing a novel miRNA prediction tool, we provide a robust and unique resource that will pave the way for future studies in the field. As proof of concept, we revealed that the repression of *LRP6* by miR-182-5p, which promotes lipogenesis and impairs glucose homeostasis, provides a novel mechanistic link between T2D and non-alcoholic fatty liver disease, and demonstrate in vivo that miR-182-5p can serve as a future drug target for the treatment of obesity-driven hepatic steatosis.

**Graphical Abstract:** 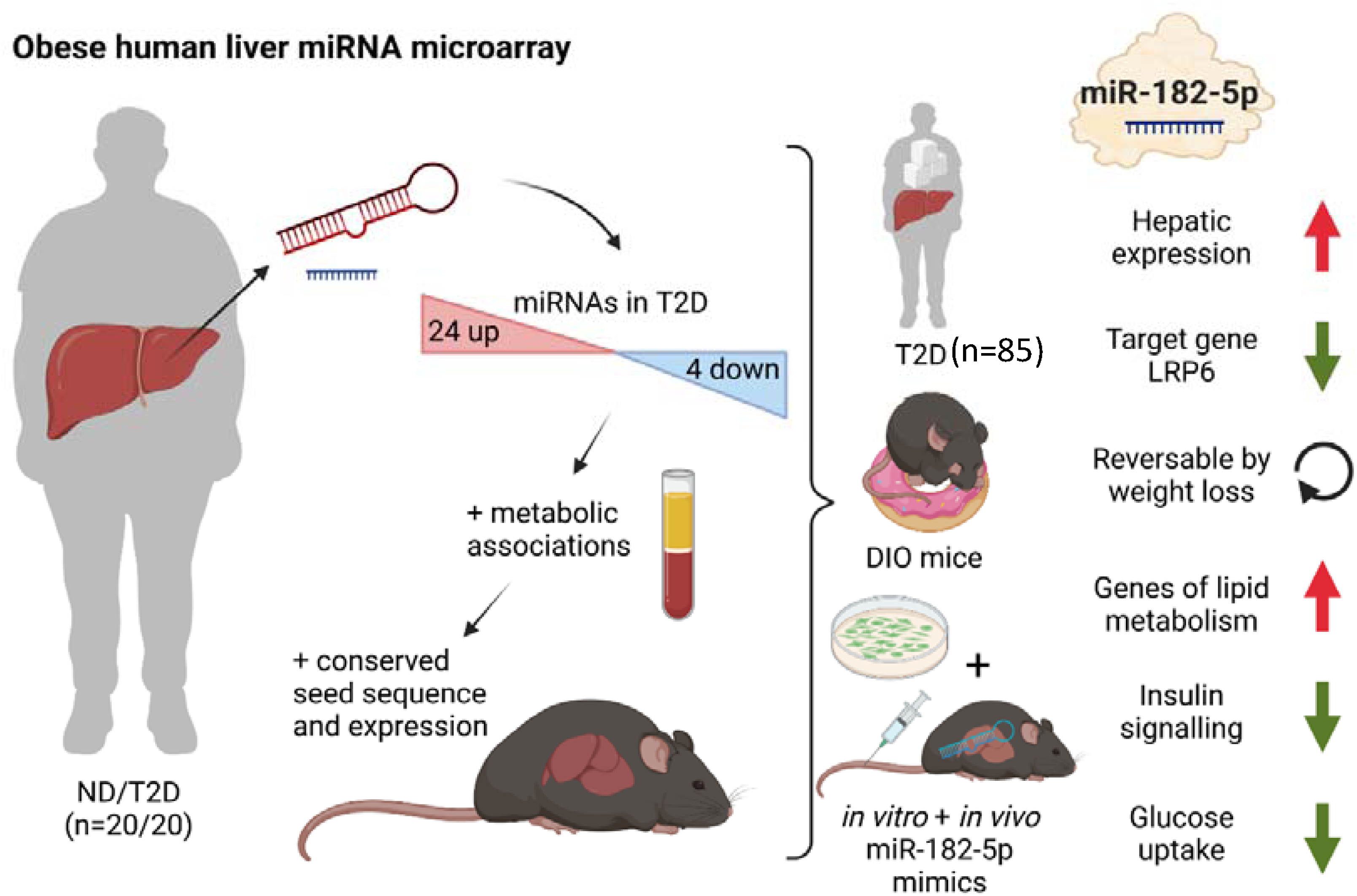

## 1. Introduction

The global prevalence of obesity has drastically increased over the past years and is associated to numerous metabolic and psychological diseases [1]. Accordingly, obesity is the leading risk factor for type 2 diabetes (T2D) and non-alcoholic fatty liver diseases (NAFLD). Yet, while almost 90% of individuals with T2D are overweight or obese, not all obese individuals develop T2D [2]. Multiple factors play a role in the development of T2D, including genetics, lifestyle, and the gut microbiome [3]. Non-genetic risk factors such as epigenetic changes induced by aging, weight gain, and intrauterine programming were shown to contribute to the increasing prevalence of T2D [4]. microRNAs (miRNAs), as one epigenetic mechanism, have the potential to simultaneously regulate the expression of multiple genes and entire pathways [5]. Additionally, miRNAs can mediate organ cross talk [6], and the dysregulation of a few miRNAs can have deleterious effects on a systemic level (reviewed in [7]). Aberrant expression of miRNAs in T2-diabetic subjects was already shown in human adipose tissue, pancreatic beta cells, skeletal muscle and blood (reviewed in [8]). However, although the liver is the central organ to maintain whole-body glucose homeostasis [9] little is known about alterations of hepatic miRNA expression in T2-diabetic subjects.

miR-103 and miR-107 directly regulate insulin sensitivity in liver and adipose tissue of obese mice by repressing caveolin-1 and insulin receptor signal transduction [10]. Likewise, miR-802 impairs hepatic insulin sensitivity [11] and the highly expressed miRNAs miR-34a [12] and miR-122 [13] perturb hepatic glucose and lipid metabolism in mice. Nevertheless, most studies on hepatic miRNAs in T2D pathogenesis were performed in laboratory rodents [10,14–16] or liver cell lines [17], human data is largely elusive. We have shown that miR-let-7e is increased in livers of T2-diabetic subjects and associated with down-regulation of the insulin receptor substrate 2 gene (*IRS2*) [18]. A comprehensive evaluation of all hepatic miRNAs transcribed and dysregulated in T2-diabetic subjects is nonetheless missing. Importantly, miRNAs are already used as drug-targets for cancer treatments and provide an excellent basis to develop novel therapies for metabolic diseases [19]. Accordingly, here we aimed to conduct a thorough profiling of hepatic miRNA signatures that associate with T2D in obese human subjects with or without type 2 diabetes, to identify miRNAs that control key genes involved in metabolic pathways as well as NAFLD and hepatic insulin resistance etiology.

## 2. Methods

A detailed description of all methods can be found in the supplemental material.

### 2.1 Study design human cohort

All participants of the human cohort signed informed consent. The study was approved by the local ethics committee (PV4889) and conformed to the ethical guidelines of the 1975 Declaration of Helsinki.

Liver biopsies were obtained during bariatric surgery at the University Hospital Eppendorf (UKE, Hamburg) as previously described [22]. Blood samples were drawn at the day of surgery after 6 h fasting. Subjects were categorized by (1) the use of anti-diabetic medication and (2) by definition of the American Diabetes Association (ADA) into subjects with T2D (receiving anti-diabetes medication or HbA1c ≥ 6.5%, n=41) and non-diabetic controls (not receiving anti-diabetes medication and HbA1c < 5.7%; n=44, Supplementary Table 1). The microarray sub-cohort (n=20 T2-diabetic and n=20 non-diabetic control subjects, Supplementary Table 1) was matched for age, sex and BMI and was part of the complete study cohort. Both disease groups contained individuals with different stages of NAFLD determined as NAFLD activity score (NAS) ranging from 0 (no NAFL) to 7 (high degree of fibrosis and steatosis), which was considered during data analysis. The sectional reporting guidelines were used according to STROBE.

### 2.2 Study design mouse cohorts

The murine studies were approved by the State of Bavaria, Germany. All experiments were performed in adult male C57BL/6J mice purchased from Janvier Labs (Saint-Berthevin, Cedex, France). Mice were maintained on a 12-h light-dark cycle with free access to water and standard chow diet (Altromin, #1314) or 58% high fat diet (HFD) (Research Diets, D12331). For the weight cycling study, mice were subjected to HFD for 24 weeks before intervention: The group termed HFD remained on HFD during the study. The remaining 2 groups of DIO mice were divided as follows: Diet switch (HC) animals continued to receive ad libitum access to HFD for 12 weeks before they were switched to chow diet for the remaining 12 weeks of the study. In contrast, weight cycling (YoYo) mice were switched to chow diet on day 0 of the study for 12 weeks and then fed again ad libitum with HFD for another 12 weeks. Age-matched mice fed chow were used as a control group. For organ collection, mice were sacrificed after a 3h fast.

Mice used for the DIO miRNA microarray analysis were fed HFD for 28 weeks. Hepatic triglyceride content was determined using the triglyceride assay kit (Wako Chemicals, Neuss, Germany). Plasma leptin and insulin levels were measured by murine ELISA kits (Merck Millipore, Darmstadt, Germany, Leptin: EZML-82K, Insulin: EZRMI-13K). Group sizes were based on power analysis.

### 2.3 Gene expression in human liver

Whole cell RNA was extracted (miRNeasy mini kit QIAGEN, Hilden, Germany) and reverse transcribed (SuperScript VILO cDNA synthesis kit Invitrogen, Carlsbad, US). Gene expression was measured by qPCR in duplicates and calculated with the ΔΔCt method. Data were normalized to CASC3 expression as reference gene [18]. Primers sequences and assay-IDs are given in tables A.2 and A.3

### 2.4 Array-based miRNA transcriptome analysis

The miRNA transcriptomes from human (n=20 type-2-diabetic, n=20 non-diabetic) and mouse (n=6; kept on HFD for 28 weeks) livers were measured with the GeneChip^TM^ miRNA 4.0 Array (Applied Biosystems, Foster City, US).

### 2.5 Gene expression analysis of murine liver samples

Whole cell RNA was extracted from snap frozen liver (miRNeasy mini kit QIAGEN, Hilden, Germany) and reverse transcribed (QuantiTect Reverse Transcription kit QIAGEN, Hilden, Germany). mRNA expression was calculated with the ΔΔCt method using Hprt1 as housekeeping gene. Primer sequences are listed in Supplementary Table 2.

### 2.6 microRNA expression analysis in human and murine liver

Human or mouse liver RNA was reverse transcribed with the qScript microRNA cDNA synthesis kit (Quanta Bioscience, Beverly, US). Self-designed forward and a pre-designed universal reverse primers were used to quantify miR-182-5p expression by SYBR green qPCR (PerfeCTa SYBR Green SuperMix, Quanta Bioscience, Beverly, US) in duplicates. Data were analyzed with the ΔΔCt method using hsa-miR-24-3p as housekeeping gene (Supplementary Table 2)

### 2.7 miRNA isolation and miRNA expression analysis in human serum

miRNAs were extracted from serum (miRNeasy Serum/Plasma Advanced kit QIAGEN, Hilden, Germany) with the spike-in control cel-miR-39-3p (QIAGEN, Hilden, Germany) and reverse transcribed (qScript microRNA cDNA synthesis kit Quanta Biosciences, Beverly, US) and quantified as described above using cel-miR-39-3p as reference.

### 2.8 Luciferase reporter gene assay

The 3’-UTR of *LRP6* containing a potential 7mer-m8 recognition site was generated by PCR amplification from human hepatic cDNA (Supplementary Table 2). The PCR fragment was directionally cloned downstream of the Renilla luciferase ORF in the psiCHECK-2 vector (Promega, Madison, US) using the Quick Ligase kit (M2200, NEB, Ipswich, US). For a site-directed mutagenesis the central guanine of the seed sequence was exchanged for an adenosine. HEK-293 cells were co-transfected in triplicates with miRNA precursor mimics (pri-miRNA-182-5p PM12369 or negative control #1 AM17110, Ambion, Applied Biosystems, Foster City, US) and the psiCHECK-2 vector containing the 3’UTR with either the consensus seed or the mutated seed sequence. Cells were harvested after 48 h. Renilla and Firefly luciferase signals were measured using the Dual-Glo® luciferase reporter gene assay (Promega, Madison, US). Each experiment was performed three times.

### 2.9 miRNA overexpression in HepG2 cells

HepG2 cells (ATCC, Manassas, US, Mycoplasma-free) were reverse transfected with miRNA precursor mimics or Negative control #1 as above. RNA was isolated (miRNeasy mini kit, QIAGEN, Hilden, Germany) and reverse transcribed (TaqMan^TM^ Advanced cDNA synthesis kit, Applied Biosystems, Foster City, US and High-Capacity cDNA Reverse Transcription Kit,Applied Biosystems, Foster City, US). miR-182-5p expression was quantified by qPCR using miR-24-3p as housekeeper. Target genes were measured as described above. LRP6 protein was quantified by Immunoblot after 72 h of miRNA mimic incubation. For western blot analysis for the measurement of phospho-Akt, transfected cells were treated with 20 nmol/l insulin (Lantus 100 E/ml, Sanofi, Paris, FR) for 10 min prior harvest. Each experiment was performed three times.

### 2.10 Western Blot analysis

HepG2 protein lysate was separated by SDS-Page (TGX Stain-Free^TM^ FastCast^TM^ Acrylamide Kit, Bio-Rad Laboratories, Inc., Hercules, US) and blotted on nitrocellulose membranes (Trans-Blot® Turbo^TM^, Bio-Rad Laboratories, Inc., Hercules, US). The blots were incubated with primary antibodies against LRP6 (1:1000, EPR2423(2) ab134146 rabbit mAb, lot GR3256666-1, Abcam, Cambridge, UK), phospho-Akt Ser473 (1:1000, D9E XP rabbit mAb #4060, lot 23, Cell Signaling Technology, Danvers, US), total Akt (1:1000, rabbit pAb #9272, lot 28, Cell Signaling Technology, Danvers, US) or HSP90 (1:1000, C45G5 rabbit mAb, lot 5, Cell Signalling Technology, Danvers, US). Each experiment was performed three times.

### 2.11. Glucose uptake of HepG2 cells

miR-182-5p precursor mimics, anti-miR miRNA inhibitors (AM10801, Ambion, Applied Biosystems, Foster City, US) and corresponding controls were reverse transfected in HepG2 cells in triplicates as described above and a glucose uptake assay (Glucose Uptake-Glo^TM^ Assay, Promega, Madison, US) was performed. Prior incubation with 1 mM 2-deoxyglucose, the cells were treated for 10 min with 20 nM insulin (Lantus 100 E/ml, Sanofi, Paris, FR). The luciferase signal was measured in duplicates and normalized to the cells treated with negative control. Each experiment was performed three times.

### 2.12 In vivo overexpression of miR-182-5p in mouse liver

Male C57BL/6J mice were purchased at 5-6 weeks of age from Janvier Labs (Saint-Berthevin, Cedex, France), maintained as described above and fed with HFD for four weeks prior and throughout the study. On day 0 and day 3.5, 1 mg/kg body weight miR-182-5p-mimic or negative control was injected via the tail vein using Invivofectamine® 3.0 (Invitrogen, Carlsbad, US). Mice were phenotypically characterized by NMR on days −1 and 7 and a glucose tolerance test after 6 h fasting on day 5. Plasma was collected from blood during the GTT to determine insulin levels by ELISA (EZRMI-13K, EMD Millipore Corporation, Burlington, US). Mice were sacrificed at day 7 after a 3h fast. Liver was snap frozen for the RNA and protein extraction as described above. Hepatic triglycerides were measured using the Triglyceride Quantification colorimetric-/fluorometric assay (MAK266, Sigma-Aldrich, St. Louis, US). A hematoxylin and eosin (HE) staining was performed from sections of the fresh frozen liver for visualization of lipid accumulation.

### 2.13 Microarray statistics

Regression analysis, statistical tests and visualization was performed by MATLAB R2020a (The MathWorks, Natick, US), R 3.5.1 (The R Foundation for Statistical Computing, Vienna, Austria) and GraphPad Prism 7.05 (GraphPad Software, Inc, San Diego, US). Data were analyzed with the Transcriptome Analysis Console (TAC) 4.0, MATLAB R2020a and R 3.5.1 and normalized by the Robust Multi-chip Analysis (RMA) algorithm. Probesets were considered expressed if more than 50 % of probes had a significantly (p<0.05) higher detection signal than the background (DABG, detection above background). Age, sex, BMI and the NAFLD activity score (NAS) were used as additional cofactors if not used as response variable in linear regression models. A p-value<0.05 was considered associated. microRNAs were filtered for confidently expressed by applying a log2 threshold of 2.3 and conserved between human and mouse was evaluated by 7mer seed match analysis.

### 2.14 General statistics

All data were adjusted for multiple testing (FDR) according to Benjamini-Hochberg when more than one test was performed. Changes in gene expression, luciferase signal, glucose uptake, protein abundance and metabolic parameter between two groups were tested by student’s t-test and between multiple groups by One-way ANOVA. All ΔCt-values which were not within a three standard deviations interval of all samples for the respective gene were defined as outliers and excluded for further analysis. Normal distribution was tested using the Lilliefors test implemented in MATLAB with a significance level of p<0.05. ΔCt values were correlated with metabolic parameters and other genes by Pearson correlation and corrected for age and gender by linear regression if applicable. Results prior adjustment are indicated with a p-value. A q-value<0.05 was considered as significant. Only applicable associations analyzed by Pearson’s correlation are plotted. Fold-change heatmaps and other graphs were generated by using GraphPad Prism 7.05 (GraphPad Software, Inc, San Diego, US).

### 2.15 Target gene and pathway analysis

Target genes of miRNAs were taken from miRTarBase and TarBase [20,21] and integrated into a SQLite database and queried by a function we termed ‘miRNA Nvis’ implemented in MATLAB R2020a (The MathWorks, Natick, US). Based on the mature miRNA and 3’-UTR sequence of the target genes miRNA Nvis performs a seed match for different pairing modes [22] and computes favorable conditions for seed binding [22,23]. Target genes were then manually selected according to literature research and after stratification for T2D-related gene ontology terms. For pathway enrichment analysis, we used only experimentally validated target genes from miRTarBase [20] as input for enrichKEGG from clusterProfiler (version 3.0.2 and R version 3.5) with the options organism = “hsa”, keyType = “kegg”, minGSize = 1 and otherwise default parameters.

### 2.16 Data and code availability

Data for the human (https://www.ncbi.nlm.nih.gov/geo/query/acc.cgi?acc=GSE176025) and murine (https://www.ncbi.nlm.nih.gov/geo/query/acc.cgi?acc=GSE211367) microarray analysis are uploaded at GEO. A database (SQLite) with all association parameters and any additional information required to reanalyze the data reported in this paper is available from the corresponding authors upon request. Source code for the target gene prediction by miRNA Nvis is available at GitHub (https://github.com/christinkrause55/microRNA_network_visualizer).

## 3. Results

### 3.1 Type 2 diabetes in obese individuals is associated with 28 miRNAs

To identify hepatic miRNAs that are associated to T2D independently of obesity we performed an array-based miRNA transcriptome analysis in liver biopsies of obese subjects with (n=20) and without T2D (n=20; Supplementary Table 1). The array contained a total of 4,603 human pre-miRNAs and mature miRNAs of which only 694 mature miRNAs (27% of the total array-based miRNAs) were expressed in obese livers, regardless of the presence of T2D (Fig. 1A). To increase the confidence of detection, the minimum signal threshold was set to a log2 value > 2.3 resulting in the detection of 594 mature miRNAs (23%) with high confidence in the obese human liver samples of both groups (Fig. 1A). Of the 30 pre-miRNAs that passed the minimal expression threshold 22 pre-miRNAs (73%) were also detected as mature miRNA transcripts indicating that they undergo a full maturation process in obese human livers (Fig. 1A).

**Figure 1:**
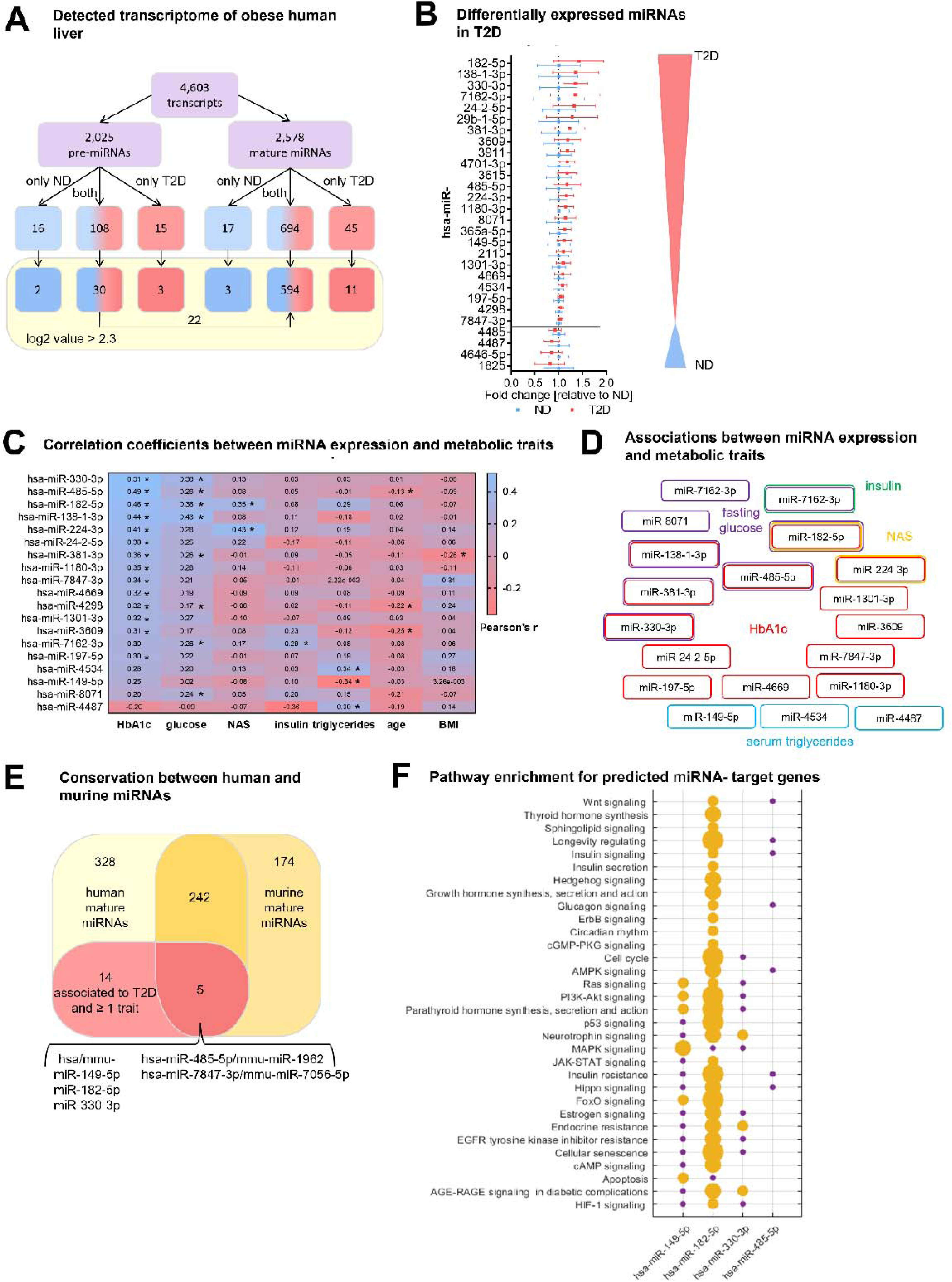
Type 2 diabetes in obese individuals is associated with a distinct hepatic miRNA signature. (**A**) Overview of all analyzed human transcripts within the GeneChip^TM^ miR4.0 assay. (**B**) Expression of the 28 regulated miRNAs in T2D compared to ND after adjustment for age, sex, BMI and NAS ranked by fold-change. Data is represented as relative mean to ND ± SD. (**C**) Pearson’s correlation coefficients (r) for the correlation between microarray miRNA log2 values and metabolic parameters. miRNAs are ranked by their association strength to first HbA1c, glucose, NAS, insulin, triglycerides, age and lastly BMI. (**D**) Mapping of significant correlations from (C) to metabolic parameters. (**E**) Venn diagram of all mature miRNA detected by microarray measurement in human and diet-induced obese (DIO) murine liver samples. (**F**) Pathway enrichment analysis of validated target genes of conserved miRNAs from (E) available in miRTarBase (19). Yellow dots indicate a significant enrichment of target genes in the respective pathway which correlates with the dot size and violet dots a potential implication meaning a non-significant enrichment of target genes in this pathway. Corrected for multiple testing: *q<0.05.

To identify hepatic miRNAs that potentially drive T2D pathogenesis in obesity we applied a logistic regression analysis for the 594 mature miRNAs using the NAFLD activity score (NAS) as cofactor to exclude any bias by hepatic fat content, lobular inflammation and fibrosis. 28 mature miRNAs were differentially expressed in T2D subjects compared to obese non-diabetic (ND) controls of which 24 miRNAs were up and four miRNAs were down regulated (Fig. 1B and Supplementary Table 4). The expression levels of 19 of 28 differentially expressed miRNAs correlated significantly with at least one metabolic trait such as HbA1c, fasting blood glucose and NAS (Fig. 1C, Supplementary Table 4). miRNAs that were significantly associated to age or BMI were excluded (Supplementary Table 5) since we aimed to uncover miRNAs that contribute to T2D in obese subjects irrespective of age and BMI. Six miRNAs were associated with two metabolic traits and miR-182-5p was the only miRNA associated with three metabolic traits: HbA1c, fasting glucose and NAS (Fig. 1D).

Next, we analyzed livers of DIO, insulin-resistant male C57BL/6J mice and compared the murine array-based hepatic miRNA expression with the human obese hepatic miRNA transcriptome to identify miRNAs that are conserved between species. Of the 594 mature human miRNAs detected with high confidence, 247 miRNAs (42%) were also expressed in livers of DIO mice based on a seed sequence match between human and murine mature miRNAs (Fig. 1E). The remaining 342 miRNAs were exclusively expressed in humans. Similarly, about half of the murine hepatic miRNAs (n=174) were either not conserved or not expressed in human liver. Only five out of 19 miRNAs that are dysregulated in liver of obese T2D subjects and are associated with at least one metabolic trait (HbA1c, fasting glucose, insulin, NAS, serum triglycerides) after adjustment for confounding factors (age, sex, BMI and NAS) are conserved in mice (Fig. 1E, Supplementary Table 6). Among these conserved miRNAs is miR-182-5p.

Many of the T2D-associated miRNAs identified in our human liver (Fig. 1C) were annotated only recently, and their target genes remain to be validated and integrated to public databases. However, for 4 out of the 5 conserved T2D-associated miRNAs information about potential target genes is available in miRTarBase [23]. These miRNAs and their target genes are significantly involved in metabolic pathways including insulin signaling, insulin resistance and PI3K-Akt signaling (p_adj_<0.05, Fig. 1F and Supplementary Table 7). Taken together, our data indicate that the identified hepatic miRNAs associate with T2D pathogenesis in human and murine obesity.

### 3.2 hsa-miRNA-182-5p is a gate keeper for *LRP6*-dependent regulation of hepatic glucose and lipid metabolism

Prompted by the strong association of hepatic miR-182-5p to metabolic traits and its likely regulatory impact on genes involved in glucose homeostasis and lipid metabolism, we next extended our, now qPCR-based, expression analysis of miR-182-5p to liver biopsies of our full study cohort of n=85 obese subjects of which n=44 were ND and n=41 were T2D according to ADA stratification and use of anti-diabetic medication. The full cohort included the original 40 discovery individuals (Supplementary Table 1). Confirming our microarray results, the expression of miR-182-5p was 2.3-fold upregulated in the diabetic livers (Fig. 2A). In this full cohort hepatic miR-182-5p expression was significantly correlated with age after adjustment for multiple testing (Fig. 2B, blue box), therefore we adjusted all consecutive regression analysis for age. The correlations of hepatic miR-182-5p expression with HbA1c and NAS were confirmed in the full cohort as well as a significant association with fasting glucose levels (Fig. 2 B red box). Furthermore, miR-182-5p correlated significantly with serum triglyceride levels and hepatic lipid content in the complete cohort, both important indicators of lipid homeostasis (Fig. 2B, red box). These data confirm that, miR-182-5p is potentially involved in the regulation of glucose tolerance and lipid metabolism in obese subjects. To test whether cell-free miR-182-5p might be a potential biomarker for T2D in obesity, we measured serum concentrations in a subset of n=30 obese ND and n=29 T2D subjects where non-hemolytic serum was available. Serum miR-182-5p was neither altered in T2D (Supplementary Fig. 1A) nor correlated with hepatic miR-182-5p expression (Supplementary Fig. 1B).

**Figure 2:**
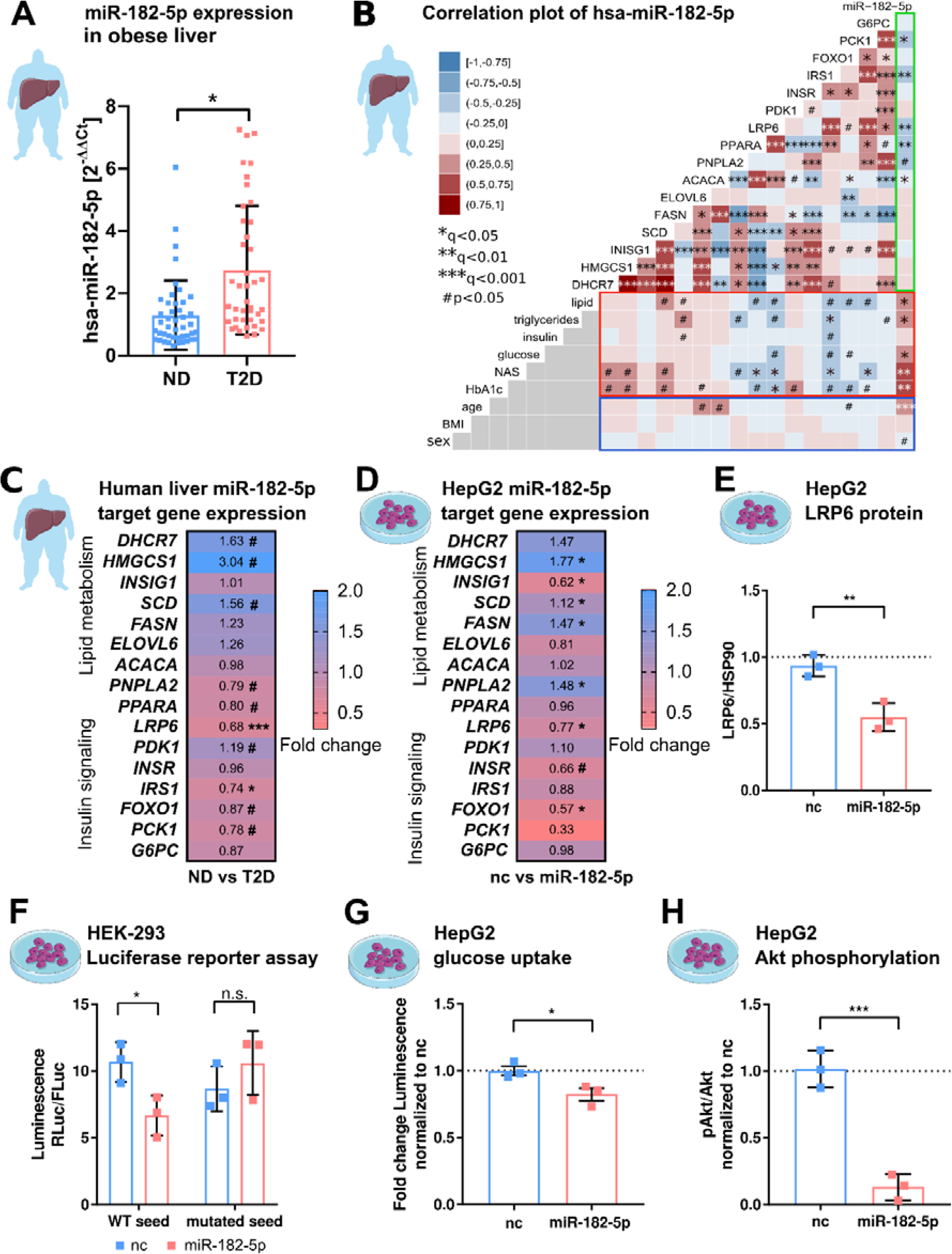
hsa-miRNA-182-5p is a gate keeper for LRP6-dependent regulation of glucose homeostasis and hepatic lipid metabolism. (**A**) Expression of miR-182-5p is 2.3-fold upregulated in liver tissue of obese subjects with type 2 diabetes (T2D, HbA1c ≥ 6.5 % or anti-diabetic medication) compared to non-diabetic (ND, HbA1c < 5.7 %) obese subjects in the extended human liver cohort (n=85). (**B**) Correlation plot of hepatic miR-182-5p expression and its target genes in human liver (green box) and of gene expression with metabolic parameters from blood (red box) as well as with confounders (blue box). Non-tested correlations are indicated by gray squares. (**C**) Expression of miR- 182-5p target genes in human diabetic liver (See also Figure A.1 C). (**D**) Expression of target genes after overexpressing miR-182-5p for 48h in HepG2 cells in comparison to a negative control (nc, n=3). (**E**) Protein abundance of the novel target gene *LRP6* is reduced after overexpression of miR- 182-5p for 72h in HepG2 cells (n=3). (**F**) Overexpression of miR-182-5p in HEK-293 cells decreases luciferase activity after 48h in a luciferase reporter assay for the LRP6 wild type (WT) sequence but not in the mutated seed (n=3). (**G**) Glucose uptake is significantly reduced (0.82-fold) in HepG2 cells after 48h of miR-182-5p overexpression and acute insulin stimulation for 20 min (n=3). (**H**) Measurement of pAkt/Akt via Western Blot indicate insulin resistance in HepG2 cells 48h after overexpression of miR-182-5p (0.13-fold). Data are shown as scatter dot plots with mean ± SD (A, E-H) or correlation matrices (B-D). Without multiple testing correction: ***p<0.001, **p<0.01, *p<0.05 (A, E-H); corrected for multiple testing: ***q<0.001, *q<0.05, significant prior to adjustment: #p<0.05 (B,C,D); Students t-test (A,C,D, E-H) or Pearson’s correlation (B).

Among the predicted target genes of miR-182-5p available in miRTarBase and TarBase are genes involved in lipid metabolism and localization (*DHCR7*, *HMGCS1*, *INSIG1*, *SCD, FASN, ELOVL6, ACACA, PNPLA2, PPARA, LRP6*) and glucose homeostasis (*PDK1, INSR, IRS1, FOXO1, G6PC*) [24,25]. According to gene ontology these genes are involved in T2D related pathways (Supplementary Table 8). Additionally, to identify previously unknown target genes of miR-182-5p we developed a prediction tool that we called *miRNA Nvis* which is based on the identification of various potential mRNA-binding sites and favorable binding conditions such as a high AU-content in the surrounding sequence, a non-central location within the 3’-UTR and additional base pairing on the 3’ end of the seed [5,28]. Accordingly, we identified *PCK1* as a novel potential target of miR-182-5p (Supplementary Table 7). Using *miRNA Nvis* we could confirm the 3’UTR seed match for all database targets except for *ACACA*. Interestingly, only eight of the 16 predicted target genes of miR-182-5p had previously been validated using cell lines (Supplementary Table 9). *ELOVL6* and *LRP6* were previously non-significantly correlated with the expression of hsa-miR-182-5p in liver metastasis [29]. To corroborate and extend these findings to humans, we measured the hepatic expression of T2D-related miR-182-5p target genes in our full cohort. The expression of *LRP6, IRS1, PNPLA2*, *PPARA, PCK1* and *FOXO1* was significantly (^#^p<0.05) reduced in T2D compared to ND controls, whereby *LRP6* and *IRS1* were the only targets passing adjustment for multiple testing (*q<0.05; ***q<0.001) (Fig. 2C, Supplementary Fig. 1C). The hepatic expression of *LRP6*, *IRS1, PPARA* and *PCK1* correlated negatively with hepatic miR-182-5p levels (Fig. 2B, green box and Supplementary Fig. 1D-G) and with at least one parameter of lipid and glucose homeostasis (Fig. 2B, red box). *ACACA* correlated with hepatic miR-182-5p but was not associated to any other parameter. We also observed an intercorrelation between target genes. Reduced *LRP6* was associated to elevated *SCD* and *HMGCS1* expression and to decreased IRS1 expression (Fig. 2B).

To validate the regulation of target genes by miR-182-5p, we transfected HepG2 cells with a miR-182-5p mimic to overexpress miR-182-5p (Supplementary Fig. 2). miR-182-5p overexpression significantly increased the expression of *HMGCS1*, *SCD*, *FASN* and *PNPLA2* (*q<0.05) and reduced the expression of *INSIG1*, *LRP6, FOXO1* (*q<0.05) and *INSR* (^#^p<0.05; Fig. 2D). This pattern of miR-182-5p target gene regulation in HepG2 cells partly mirrored the expression pattern observed in livers of T2D subjects. In both conditions, expression of *SCD* and *HMGCS1* was upregulated. Nevertheless, *ELOVL6* and *PNPLA2* showed an opposite trend and *INSR* and *FASN* showed a less pronounced effect in human liver (Fig. 2C). *LRP6* was the only target gene that was consistently repressed in human liver and in HepG2 cells after miR-182-5p mimic transfection. Consequently, LRP6 protein levels were significantly decreased after miR-182-5p overexpression in HepG2 cells (Fig. 2E and Supplementary Fig. 3A) and renilla luciferase intensity reporting *LRP6* expression was significantly repressed upon miR-182-5p overexpression in comparison to negative control (nc) transfected cells (Fig. 2F, left panel). However, when we mutated the central cytosine of the seed sequence into an adenosine (5’-TTG**C**CAA to 5’-TTG**A**CAA) *LRP6* expression was unaltered thus confirming sequence specificity of miR-182-5p/LRP6 pairing (Fig. 2F, right panel). Lastly, miR-182-5p overexpression significantly decreased glucose uptake (Fig. 2G) and suppressed insulin signaling in HepG2 cells (Fig. 2H and Supplementary Fig. 3B). Reducing basal miR-182-5p in HepG2 cells by an antagomir did not change glucose uptake (Supplementary Fig. 4A and B).

Overall, these results suggest that miR-182-5p might enhance hepatic *de novo* lipogenesis via repression of LRP6 and disinhibition of *HMGCS1*, *SCD* and *FASN.* Furthermore, it decreases hepatic insulin sensitivity by reducing *IRS1* and *INSR* mRNA in obese T2D subjects.

### 3.3 Hepatic miR-182-5p expression in obese mice can be reversed by weight-loss

To corroborate that hepatic miR-182-5p plays a role in the regulation of glucose homeostasis and hepatic lipid metabolism, we performed a weight cycling experiment in DIO, glucose intolerant male C57BL/6J mice (Fig. 3A). Hepatic miR-182-5p expression was significantly increased in obese mice exposed to HFD for 24 weeks compared to lean, chow-fed controls (1.9-fold, Fig. 3B), while 12 weeks of HFD feeding were not sufficient to induce miR-182-5p expression (data not shown), indicating that miR-182-5p induction requires chronic metabolic stress and is not a marker of mere obesity. Upon weight-loss by nutritional intervention, i.e. DIO mice were switched from HFD to chow (HC) which equates to mild calorie restriction, hepatic miR-182-5p levels decreased by 30% demonstrating that the hepatic overexpression of miR-182-5p in obese mice is reversible. However, hepatic miR-182-5p expression increased to DIO levels again when the group of mice undergoing weight loss (HC) were re-exposed to HFD to regain all the previously lost weight (YoYo) (Fig. 3B, Supplementary Fig. 5A). In case of the other human-murine conserved metabolic miRNAs, only mmu-miR-149-3p followed this pattern. Other miRNAs reduced expression after HFD feeding and were only partially reversible (Supplementary Fig. 5B).

**Figure 3:**
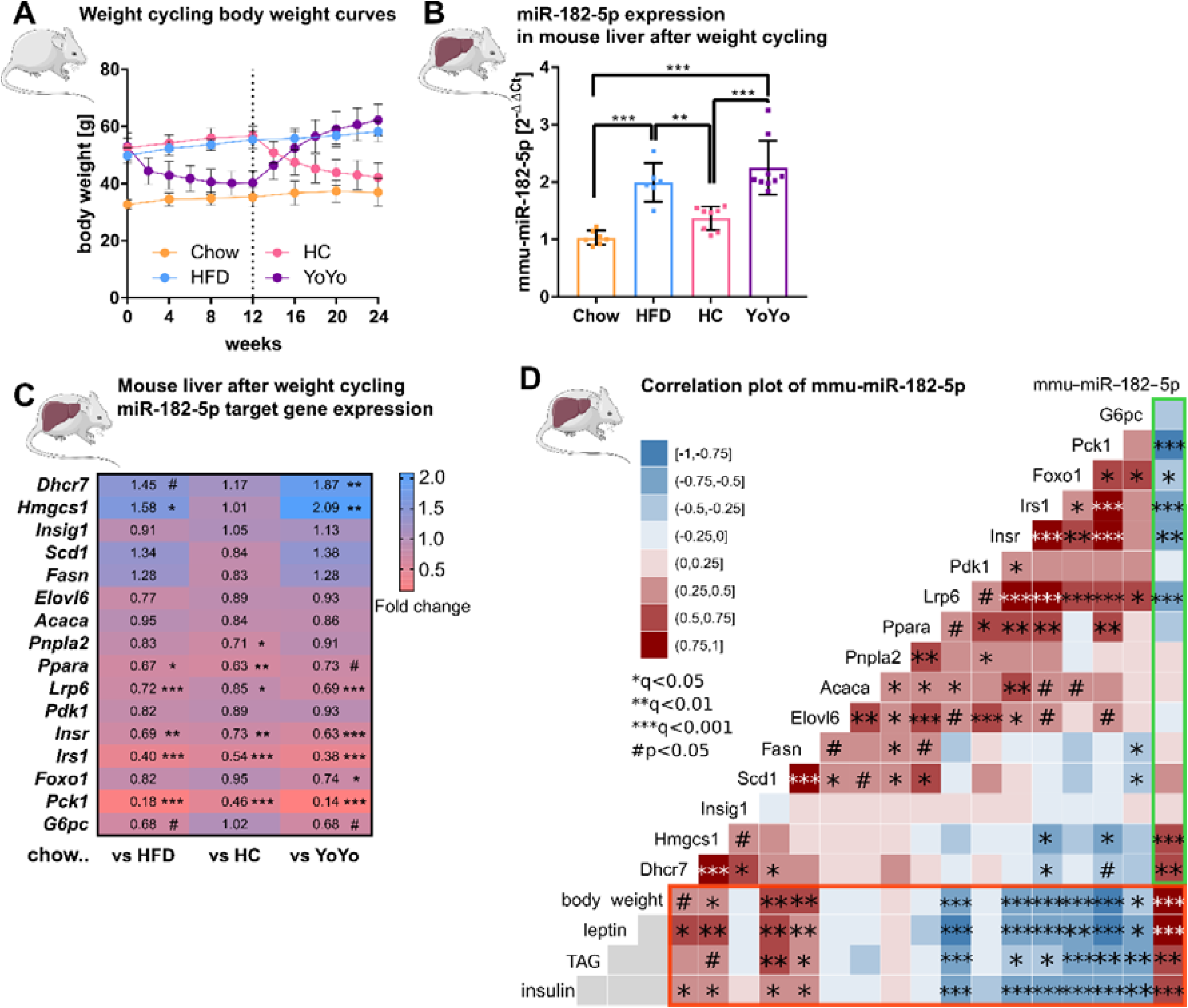
Hepatic miR-182-5p expression in obese mice can be reversed by weight-loss. (**A**) Body weight in gram of mice in the weight cycling cohort (n=7 Chow, n=6 HFD, n=8 HC, n=9 YoYo). Groups of obese HFD-fed mice were switched to chow for 12 weeks (HC) to induce body weight loss and re-fed with HFD for another 12 weeks (YoYo). (**B**) Expression of miR-182-5p in liver of mice undergoing weight cycling. (**C**) Expression of target genes of miR-182-5p in murine liver compared to the Chow control group (see also Figure S2). (**D**) Correlation plot of hepatic miR-182-5p expression and its target genes in murine liver (green box) and of gene expression with relevant metabolic parameters (red box). Non-tested correlations are indicated by gray squares. TAG: triacylglycerol. Data are shown as mean ± SD (A,B) or correlation matrices (C,D). Corrected for multiple testing: ***q<0.001, ***q<0.01, *q<0.05, significant prior to adjustment: #p<0.05 (C, D); One-Way ANOVA (B) or Pearson’s correlation (C,D).

Next, we measured the 16 miR-182-5p target genes in our murine weight cycling model. The hepatic expression of *Lrp6*, *Insr*, *Irs1, Ppara* and *Pck1* was significantly reduced in HFD-fed mice compared to chow controls (Fig. 3C) which is consistent with the increased expression of miR-182-5p and the regulation of these target genes -except for *INSR* - in obese T2D human livers. However, whereas *INSR* was not changed in human liver it was significantly decreased in HepG2 cells after miR-182-5p mimic transfection. Upon weight loss, target gene expression was partially equalized or improved in comparison to chow mice and in comparison to the HFD group the expression of some downregulated miR-182-5p target genes e.g. *Pck1* was at least partially reversed. Upon weight re-gain miR-182-5p expression increased in the YoYo mice (Fig. 3B) and expression of the targets *Lrp6*, *Irs1* and *Pck1* was significantly repressed (Fig. 3C and Supplementary Fig. 5C).

The correlation analysis confirmed the strong associations of miR-182-5p with *Pck1*, *Irs1* and *Lrp6 in* murine liver (Fig. 3D, green box) as observed for the human cohort. Furthermore, we could validate the miR-182-5p associated decrease of *Insr* and *Foxo1* expression, which was previously only observed in HepG2 cells. Metabolic traits such as body weight, plasma leptin and insulin as well as hepatic triacylglycerol (TAG) levels had the strongest correlations with hepatic miR-182-5p expression in obese mice also in regard to other conserved metabolic miRNAs (Fig. 3D, red box and Supplementary Fig. 5D). Taken together, the dynamic regulation of miR-182-5p and its conserved impact on key genes of insulin signaling and hepatic lipid metabolism in mice and humans (Supplementary Fig. 5E) highlight the important role of hepatic miR-182-5p in T2D.

### 3.4 miR-182-5p overexpression increases hepatic fat and insulin content in metabolically challenged mice

To validate the metabolic effects of miR-182-5p in vivo, we overexpressed miR-182-5p in male C57BL/6J mice by intravenous injections (twice in seven days) of miR-182-5p mimic or control while receiving HFD (Fig. 4A). Hepatic miR-182-5p expression increased 584-fold in mimic treated mice compared to the control group (Fig. 4B) and was highest expressed in liver compared to spleen and heart (Supplementary Fig. 6A). Fat mass tended to be increased (p=0.17, Fig. 4C) whereas body weight remained unchanged. (Fig. 4D, Supplementary Fig. 6B). Glucose tolerance (Fig. 4E, Supplementary Fig. 6C) and fasting glucose levels (Fig. 3F) were not altered after 5 days of miR-182-5p overexpression, but fasting insulin levels were significantly increased in miR-182-5p treated mice (Fig. 3G) indicating the development of impaired glucose homeostasis. Moreover, miR-182-5p overexpression significantly increased hepatic triglyceride levels (Fig. 4H) and lipid droplets (Supplementary Fig. 6D) which is in line with the trend towards increased fat mass (Fig. 4C) and our hypothesis based on the findings in obese humans and mice that hepatic miR-182-5p overexpression would worsen glucose homeostasis and liver fat content. The hepatic expression of *Lrp6, Irs1 FASN, Foxo1* and *G6pc* tended to be decreased in miR-182-5p mimic treated mice (Fig. 4I, Supplementary Fig. 6E). Importantly, hepatic LRP6 protein content decreased significantly in mice overexpressing miR-182-5p (Fig. 4J) which confirms our data from mimic-treated HepG2 cells and the predicted interaction between miR-182-5p and Lrp6. Overall, the results from this acute overexpression study indicate that miR-182-5p has a strong potential to worsen hepatic steatosis and possibly promotes insulin resistance and hyperinsulinemia via downregulation of LRP6.

**Figure 4:**
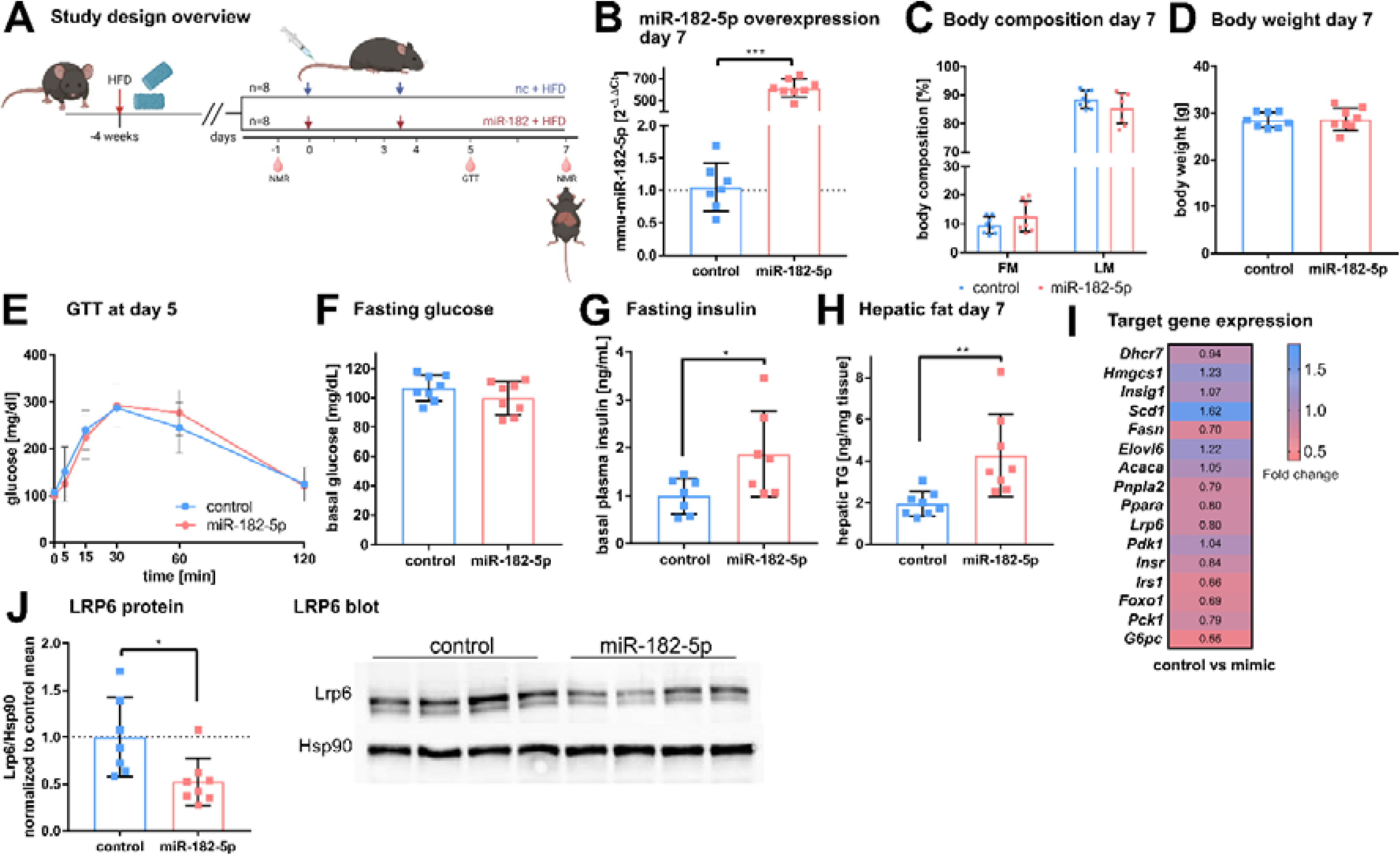
*In vivo* overexpression of miR-182-5p elevates fasting insulin levels and hepatic fat content and reduces LRP6 protein levels. (**A**) Mice were challenged with HFD for four weeks prior to the first injection of 1 mg miRNA mimic per kg body weight and sacrificed after seven days (n=8 per group). A second injection was performed after 3.5 days. Body composition was examined at days −1 and 7 by NMR. Glucose and insulin levels were determined from blood at days −1, 5 and 7. Glucose tolerance was evaluated at day 5. (**B**) Hepatic miR-182-5p expression was 584-fold upregulated at day 7. (**C**) Fat mass tended to be increased in the miR-182-5p injected group but (**D**) body weight was not different between control and miR-182-5p treated mice. (**E**) Glucose tolerance and (F) fasting glucose were not different between both groups. (**G**) Fasting insulin levels and (**H**) hepatic triglyceride content were increased 2.25-fold and 2.19-fold in miR-182-5p treated mice, respectively. (**I**) Comparable mRNA, but (**J**) 0.52-fold diminished LRP6 protein levels in liver samples of miR-182-5p treated mice. Data are shown as mean ± SD (B-H). *p<0.05, **p<0.01; Student’s t-test (B-H).

## 4. Discussion

In the present study, we describe a functional role for the miRNA miR-182-5p and its target gene *LRP6* in the regulation of glucose homeostasis and hepatic lipid metabolism in obese T2D humans. In a medium-sized but well characterized human cohort, we were able to assess for the first time the hepatic miRNA transcriptome of obese individuals with T2D compared to obese control subjects. We identified a distinct signature of 28 hepatic miRNAs that were associated with T2D of which 19 miRNAs were associated with further metabolic traits. Of those, the human/murine conserved miRNA miR-182-5p showed the strongest upregulation in T2D subjects and moreover correlated to multiple metabolic traits and pathways of glucose and lipid metabolism including fasting glucose, HbA1c and liver fat content. The newly validated miR-182-5p target gene *LRP6* was consistently downregulated in livers of obese T2-diabetics and in obese insulin-resistant mice as well as by *in vivo* and *in vitro* miR-182-5p overexpression. While hepatic miR-182-5p overexpression increased liver fat content and fasting insulin levels in mice, weight loss in DIO mice reversed miR-182-5p induction and partially restored its target genes in liver (Fig. 5) indicating a therapeutic potential for future miR-182-5p-based drugs.

**Figure 5:**
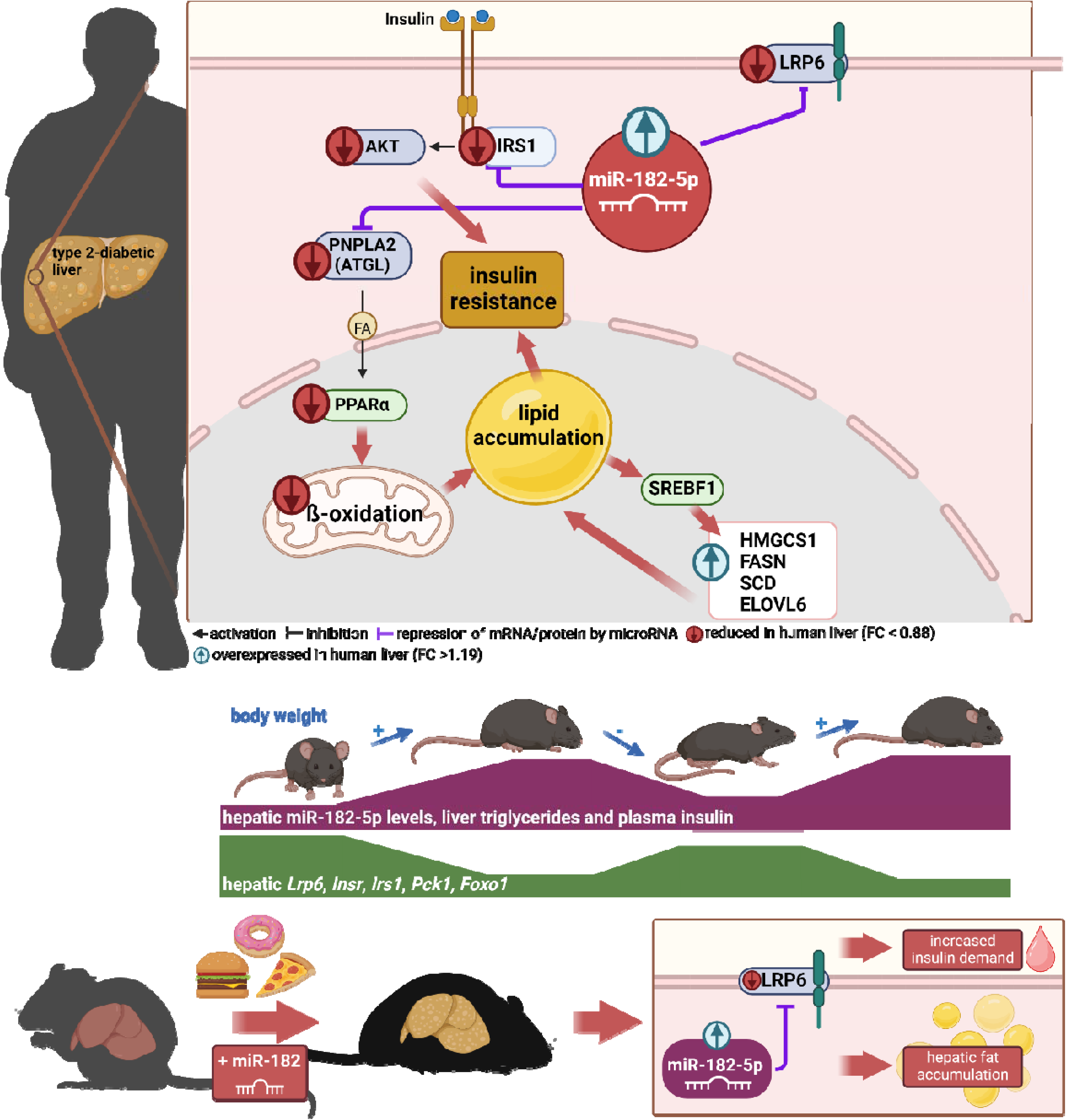
Dynamic regulation of insulin resistance and hepatic lipogenesis by miR-182-5p. Upregulation of hsa-miR-182-5p in human obese diabetic liver simultaneously decreases the metabolic pathways of beta oxidation and stimulates lipogenesis. LRP6 (highlighted) is the main target gene and consistently altered in humans, cell culture and mouse. miR-182-5p is induced by long-term feeding of high-fat diet in mice and reversed by weight loss. Identified target genes are consequently altered inversely to miRNA-182-5p expression. To prove direct effects, liver-specific upregulation of miR-182-5p in metabolically challenged mice caused a significant reduction of LRP6 which is accompanied by increased hepatic fat accumulation and increased fasting insulin levels.

Surprisingly, the here presented human hepatic miRNA transcriptome revealed that hepatic T2D-driving miRNAs previously identified in laboratory rodents [10,14–16] or liver cell lines [17], such as miR-107 [10] or miR-802 [11] are likely not dysregulated in obese T2D humans. Only five miRNAs that were dysregulated in livers of obese T2D humans were conserved in obese insulin-resistant mice. Of note, the discrepancy between the here identified miRNAs and known miRNAs regulating glucose metabolism in murine studies [10,11,13,16,26,27] may be based on the design of our human study. Both groups of our human study cohort were severely obese, and our analysis did not compare the obese miRNA transcriptome to a lean control group. Consequently, obesity-induced miRNAs likely escaped our analysis. Furthermore, in our logistic regression analysis we used the NAFLD activity score (NAS) as cofactor to exclude hepatic lipid content and fibrosis as driving factor for altered miRNA expression. Accordingly, our previously identified T2D-induced hsa-let-7e-5p [18] shows under this stratification only a strong trend (p=0.0518) to be elevated in T2D. If we exclude NAS as cofactor, we can reproduce the significant association (p=0.0265, Supplementary Table 2) which is accompanied by an aging effect (p=0.0286, Supplementary Table 3). Nevertheless, we here comprehensively evaluated all hepatic miRNAs transcribed in humans and identified miRNAs specifically dysregulated in obese T2-diabetic, compared to obese non-diabetic subjects. hsa-miR-182-5p is transcribed as part of a highly conserved cluster including miR-183/96/182 whose expression is upregulated in a variety of non-sensory diseases, including cancer, neurological and auto-immune disorders [28]. Interestingly, in our cohort, only the expression of miR-182-5p was increased and associated to T2D and other metabolic traits whereas the other cluster members were not detected in the here investigated biopsies. Our findings are in agreement with a previous study where increased hepatic miR-182-5p expression was associated to the severity of NAFLD-related fibrosis in a smaller cohort of 30 subjects matched for age, sex, BMI and T2D [29]. Furthermore, increased hepatic miR-182-5p expression levels were also reported in mouse [30] and rat [31] models of NAFLD. Here, we reproduced and mechanistically extended these findings in liver biopsies of 85 obese subjects and in DIO mice undergoing weight-cycling. In addition to the known associations of miR-182-5p with hepatic lipid metabolism, we revealed strong correlations to genes and clinical parameters associated with glucose tolerance. This new link of miR-182-5p to glucose homeostasis is further support by our *in vitro* findings of reduced glucose uptake and impaired insulin signaling after miR-182-5p overexpression. Additionally, *in vivo* miR-182-5p overexpression not only increased hepatic fat content but also increased fasting insulin levels which itself is a predictor of T2D in humans [32].

Since miRNAs can be released into circulation from various tissues, serum miRNAs are potential, minimally invasive biomarkers for tissue health [33]. However, since most of the previously reported circulating miRNAs which are associated with T2D [34,35] are not among the 28 hepatic miRNAs discovered in our analysis (Fig. 1), our human hepatic miRNA transcriptome now reveals that serum miRNAs might not be suitable to assess dysregulations of hepatic liver glucose and lipid metabolism. We could also not reproduce the previously described association between T2D and serum miR-182-5p (Supplementary Fig. 1A and B) [36]. The validity of serum miR-182-5p as biomarker for T2D manifestation [36] in a disease-duration and treatment-specific manner in T2D [37] could not be tested in our cohort as disease duration and precise duration and dosages of the diabetes treatment was not available for all study participants. Furthermore, serum miR-182-5p concentrations might be biased by miR-182-5p released from dead cells of other tissues than liver [38] and our analysis did not distinguish between cell-free miRNA and extracellular vesicular miRNAs, potentially impacting the diagnostic potential [39]. Importantly, recent motif analyses identified that miRNAs might contain liver-specific cell-retention signals, such as “AGAAC” that prevent their release into circulation [40]. This retention signal is present in hsa-miR-182-5p (UUUGGCAAUGGU**<u>AGAAC</u>**UCACACU) [40] and could explain why we did not find differences in serum miR-182-5p between our study groups. Furthermore, it strengthens our assumption that miR-182-5p is particularly relevant for regulating liver metabolism in an autocrine/paracrine manner. Lastly, our results highlight the risk to miss important disease mechanisms when solely focusing on blood-based epigenetic markers [41] and the urgent need for tissue-specific resource databases as presented in this study.

According to the miRNA-gene interaction database [21], the lipogenic target genes of miR-182-5p *SCD* and *FASN* should be downregulated upon miR-182-5p induction. However, we hypothesize that the increased expression of key genes of cellular *de novo* lipogenesis is rather an effect of the miR-182-5p mediated repression of *LRP6* than a direct miRNA-mRNA interaction*. LRP6* is a Wnt co-receptor [42,43] and is linked to metabolic diseases in humans since several loss-of-function mutations within *LRP6* were shown to cause hypertriglyceridemia [44,45], hypercholesterinemia [46,47], NAFLD [48] and atherosclerosis [45,49]. Loss of *LRP6* mediated activation of the Wnt/beta-catenin signaling pathway (Fig. 5) enhances hepatic lipid accumulation by increasing *de novo* lipogenesis and triglyceride synthesis [42,46] which is in line with the observed increased expression of *SCD* in T2D obese humans and *SCD* and *FASN* in HepG2 cells after miR-182-5p mimic transfection. Additionally, mice carrying the loss-of-function mutation LRP6 p.R611C develop fatty liver disease including insulin resistance, liver inflammation and steatosis due to increased hepatic *de novo* lipogenesis [50]. Besides its crucial role as regulator of *de novo* lipogenesis *LRP6* regulates glucose metabolism by promoting TCF7L2-dependent insulin receptor expression in humans [51] and by regulating *IRS1* expression in cells [52]. Our findings of reduced *IRS1* expression in diabetic obese humans and reduced *INSR* and *IRS1* expression in DIO mice as well as downregulated glucose uptake and insulin signaling in miR-182-5p overexpressing cells are consistent with this glucoregulatory role of *LRP6*, acting under tight control by miR-182-5p. Our acute hepatic miR-182-5p overexpression study in mice strongly corroborates this miR-182-5p-Lrp6 link, and warrants additional, long-term studies on whole body glucose tolerance and hepatic metabolism in miR-182-5p overexpressing mice.

In naïve mice, hepatic miR-182-5p expression was only upregulated after 20 weeks of HFD feeding, i.e. long after the mice develop obesity compared to standard diet fed mice. Accordingly, obesity *per se* does not seem to be the sole driver of hepatic miR-182-5p. This is consistent with the human situation, were only diabetic obese but not “healthy” obese humans show the increase of hepatic miR-182-5p expression. Whether miR-182-5p dictates if an obese individual becomes type 2 diabetic or remains glucose tolerant, and whether miR-182-5p mechanistically drives hepatic dysregulation of glucose and lipid metabolism, are thus intriguing questions for future studies. Such follow-up studies should be conducted in humans, given that mice are rarely developing diabetic pathologies surpassing glucose intolerance, and should compare lean with overweight or obese non-diabetic and T2D subjects to specifically address whether exacerbated hepatic insulin resistance and lipid deposition affect miR-182-5p expression levels. In our cohort, T2D subjects received anti-diabetic treatment, which is a limitation of our study. HbA1c levels in the T2-diabetic subjects were nonetheless still significantly higher compared to the ND obese individuals (Supplementary Table 1). To ultimately dissect if hepatic miR-182-5p upregulation is causally involved in the development of insulin resistance or if it is a marker of manifested T2D would mandate longitudinal studies. First mechanistic evidence is revealed by our *in vitro* and *in vivo* studies, which show that miR-182-5p overexpression acutely decreases glucose uptake and insulin signal transduction in hepatic cells and increases fasting insulin levels and hepatic fat content in mice. Future studies should investigate the mechanisms inducing miR-182-5p expression in the liver of T2-diabetic subjects, and whether miR-182-5p antagonism can rescue the metabolic phenotype in mice.

## 5. Conclusions

We provide a network of novel microRNAs that are dysregulated in livers of T2-diabetic subjects and identify miR-182-5p and its target genes as potential drivers of dysregulated glucose tolerance and fatty acid metabolism in obese T2-diabetics. Mechanistic studies with miRNA mimics in cells and mice revealed that hepatic miR-182 expression elicits repressive effects on its target gene *LRP6* and subsequently on glucose and lipid metabolism. The discoveries that miR-182-5p is associated to T2D and NAFLD via *LRP6* downregulation in liver of obese subjects and the reversibility by weight loss in DIO mice offer unique mechanistic insight into diabetes pathology and point towards promising future anti-diabetic strategies, built either on antagonizing miR-182-5p activity or reducing hepatic miR-182-5p expression by pharmacological or dietary means.

## Abbreviations

ADA: American Diabetes Association
BMI: body mass index
DIO: diet-induced obese
miRNA: microRNA
HbA1c: hemoglobin A1c
NAFLD: non-alcoholic fatty liver disease
NAS: NAFLD activity score
ND: non-diabetic
qPCR: quantitative real-time polymerase chain reaction
STROBE: STrengthening the Reporting of OBservational studies in Epidemiology
T2D: type 2 diabetes
UTR: untranslated region

## Acknowledgements

We thank M. Grohs (Institute for Human Genetics, University of Lübeck), P. Schroeder and A. Heinicke (both Department of General, Visceral and Thoracic Surgery, University Medical Center Hamburg-Eppendorf) as well as Katrin Huber, Noémi Mallet and Miriam Krekel (Research Unit NeuroBiology of Diabetes, Helmholtz Zentrum Munich) for their excellent technical assistance.

## Funding

This work was supported by research funding from the Deutsche Forschungsgemeinschaft (KI 1887/2-1, KI 1887/2-2, KI 1887/3-1 and CRC-TR296), the European Research Council (ERC, CoG Yoyo-LepReSens no. 101002247; PTP), the Helmholtz Association (Initiative and Networking Fund International Helmholtz Research School for Diabetes; MB) and the German Center for Diabetes Research (DZD Next Grant 82DZD09D1G). Graphics were created with Bio Render (https://biorender.com/).

## Declaration of interests

All authors declare no conflicts of interest.

## Author contributions

**Christin Krause:** Conceptualization, Methodology, Formal analysis, Investigation, Writing - Original Draft; **Jan H. Britsemmer:** Methodology, Investigation; **Miriam Bernecker:** Methodology, Investigation; **Anna Molenaar:** Methodology, Investigation; **Natalie Taege:** Methodology, Investigation; Nuria Lopez-Alcantara: Methodology, Investigation; **Cathleen Geißler:** Methodology, Investigation**; Meike Kaehler:** Methodology, Formal analysis; **Katharina Iben:** Methodology, Investigation; **Anna Judycka:** Methodology, Investigation; **Jonas Wagner:** Investigation, Project administration**; Stefan Wolter:** Investigation, Project administration; **Oliver Mann**: Resources, Project administration; **Paul T. Pfluger**: Resources, Funding acquisition, Writing - Review & Editing; **Ingolf Cascorbi:** Resources; **Hendrik Lehnert:** Resources, Funding acquisition, Writing - Review & Editing**; Kerstin Stemmer:** Conceptualization, Methodology; **Sonja C. Schriever:** Conceptualization, Methodology, Investigation, Writing - Review & Editing, Funding acquisition; **Henriette Kirchner:** Conceptualization, Methodology, Investigation, Writing - Review & Editing, Supervision, Project administration, Funding acquisition

## Appendices

### 6.1. Supplemental Figures

**Supplementary Fig. 1:**
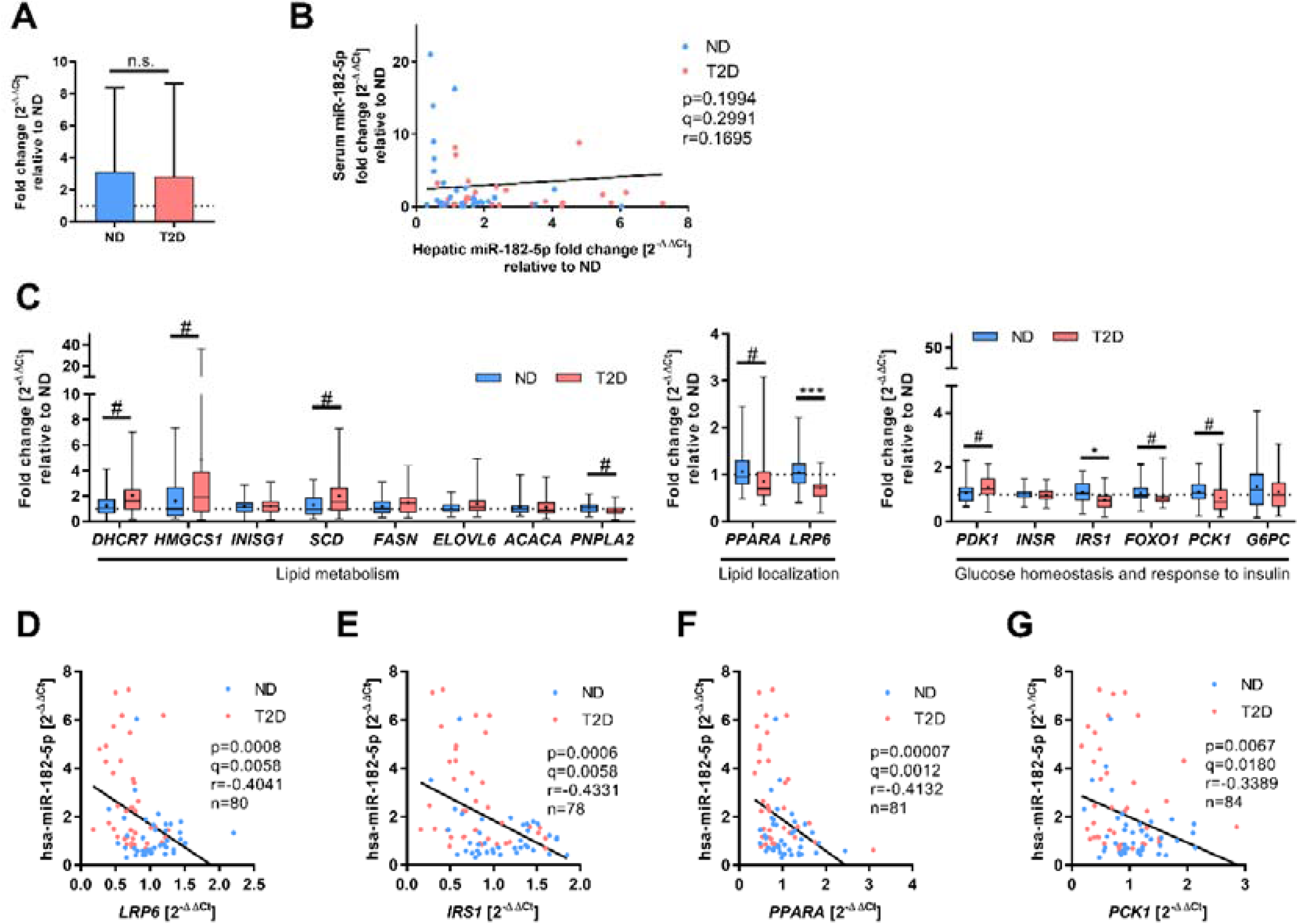
Hepatic gene expression of miR-182-5p target genes and correlation to miRNA expression as described by correlation plot (main Figure 2B) and heat map (main Figure 2C). (**A**) Serum miR-182-5p levels are not altered between obese human subjects without (ND) or with T2D. (**B**) Serum miR-182-5p levels are not associated to hepatic miR-182-5p expression in obese subjects. (**C**) Expression of metabolic miR-182-5p target genes. Correlation analysis between miR-182-5p expression and its target genes *LRP6* (**D**), *IRS1* (**E**), *PPARA* (**F**) and *PCK1* (**G**) in obese human liver. Data are shown as mean ± SD (A), scatter plot (B, D-G) or box-whisker plots with min and max values, expression mean is indicated as cross within the box plots (C). Corrected for multiple testing: ***q<0.001, *q<0.05, significant prior to adjustment for multiple testing: #p<0.05 (B,C,D); Students t-test (A,C) or Pearson’s correlation (B, D-G).

**Supplementary Fig. 2:**
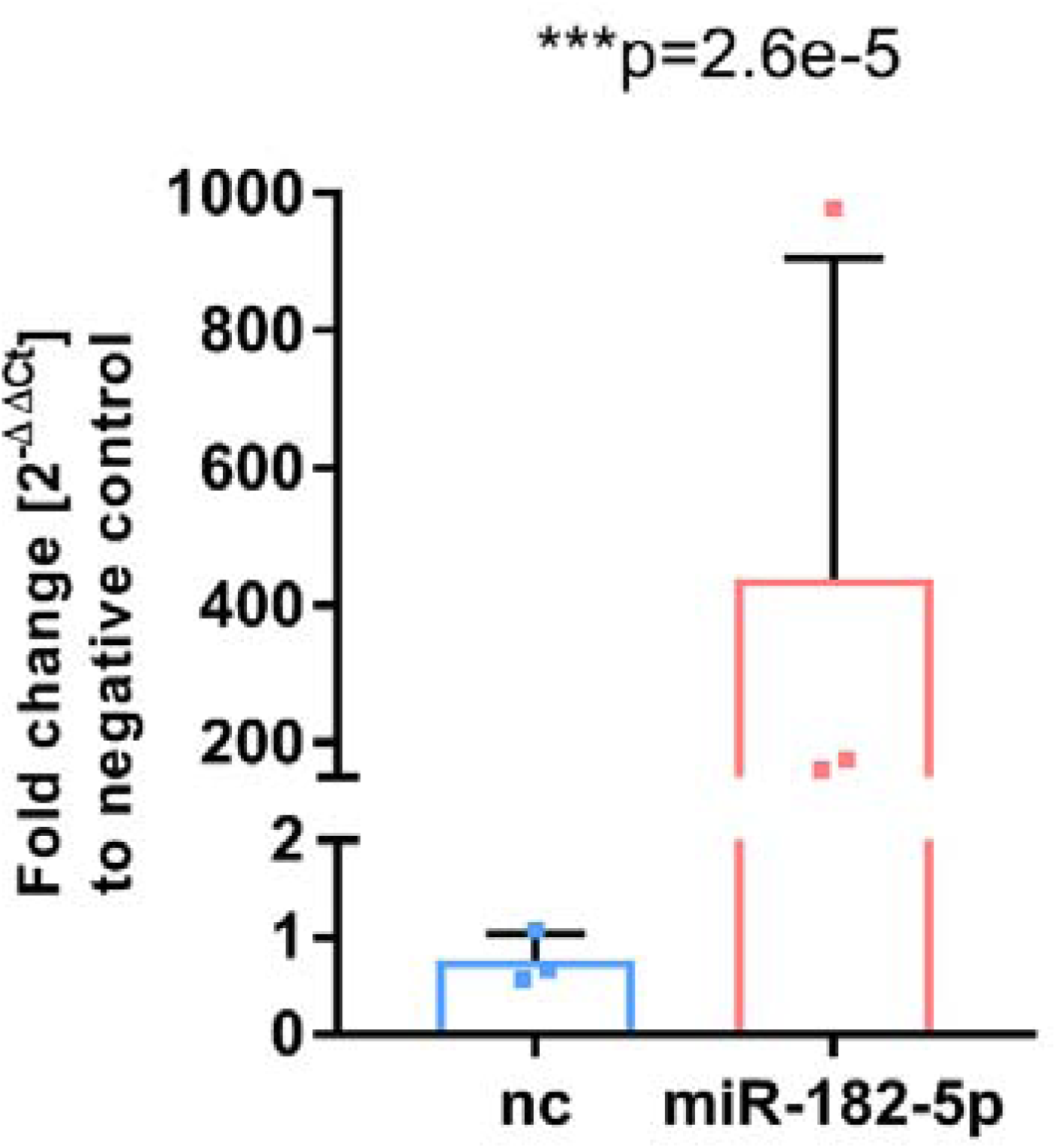
Mimic transfection in HepG2 cells led to a significant 400-fold upregulation of miR-182-5p.

**Supplementary Fig. 3:**
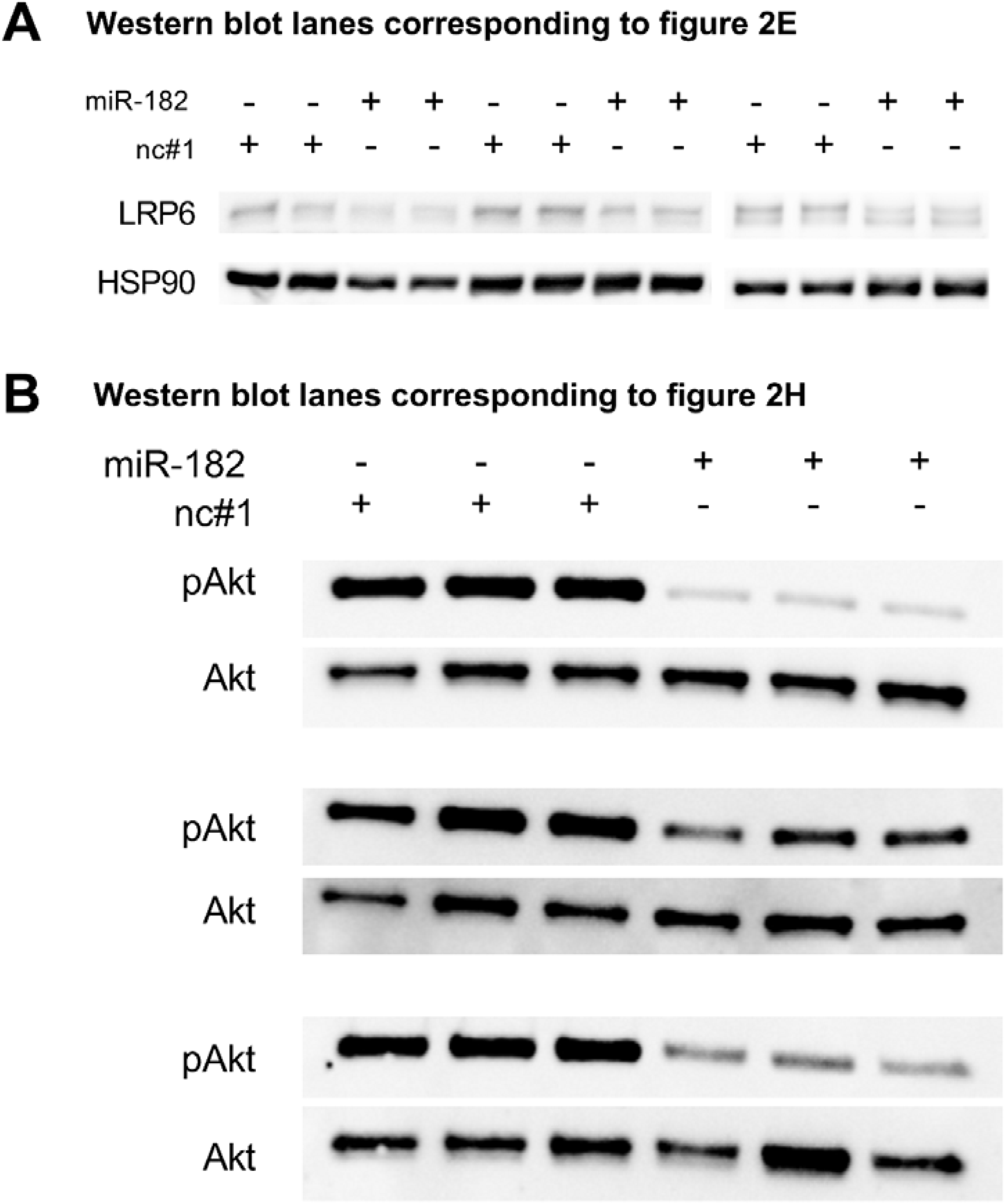
Western blot lanes used for quantification and which are corresponding to figure 2E (**A**) or figure 2H (B).

**Supplementary Fig. 4:**
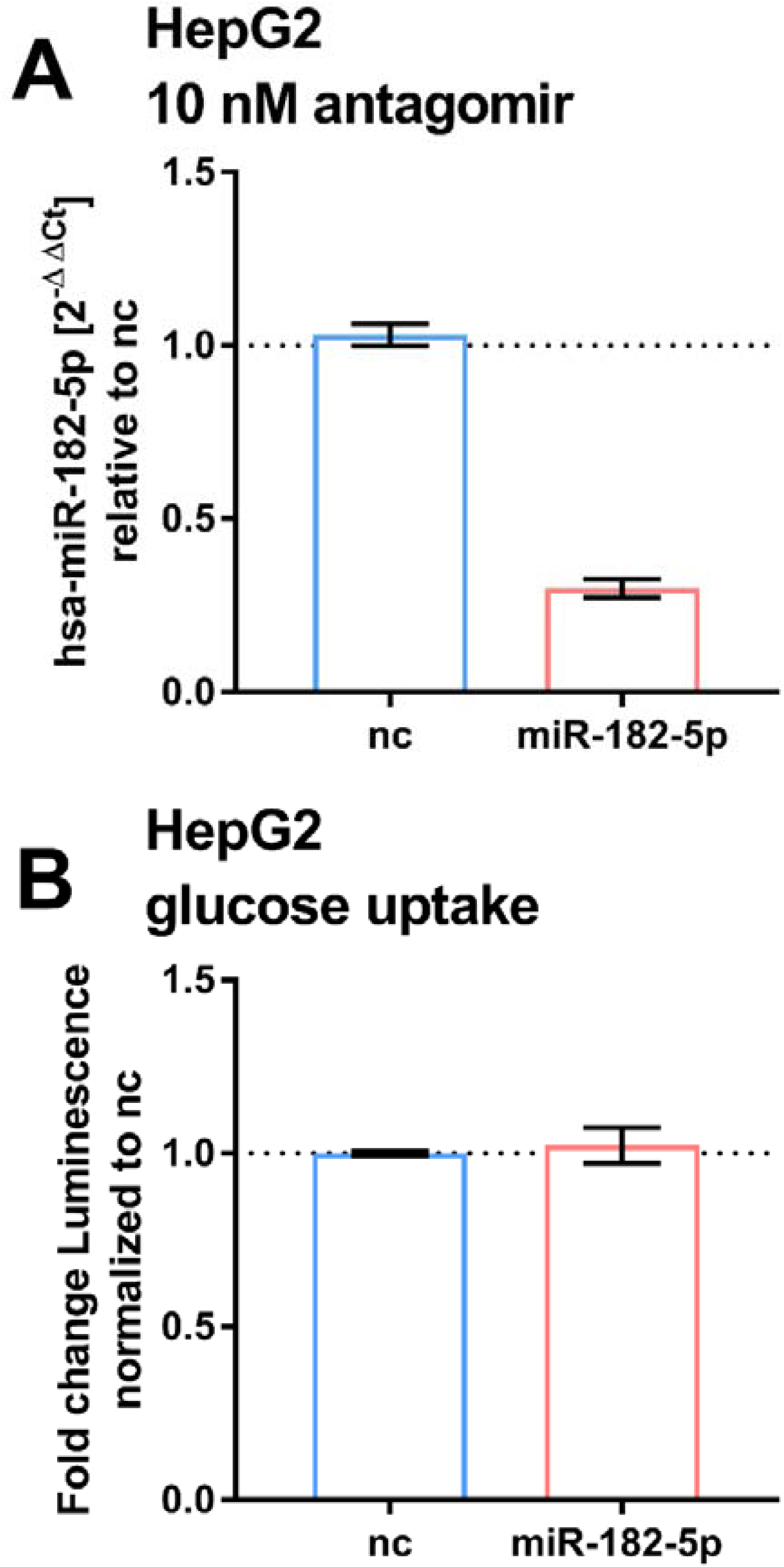
Effects of suppressing basal miR-182-5p in HepG2. **(A)** Treatment of HepG2 cells with 10 nM miR-182-5p antagomir lead to a 0.4-fold reduction of miR-182-5p expression. (B) The reduction did not lead to a significant change in glucose uptake.

**Supplementary Fig. 5:**
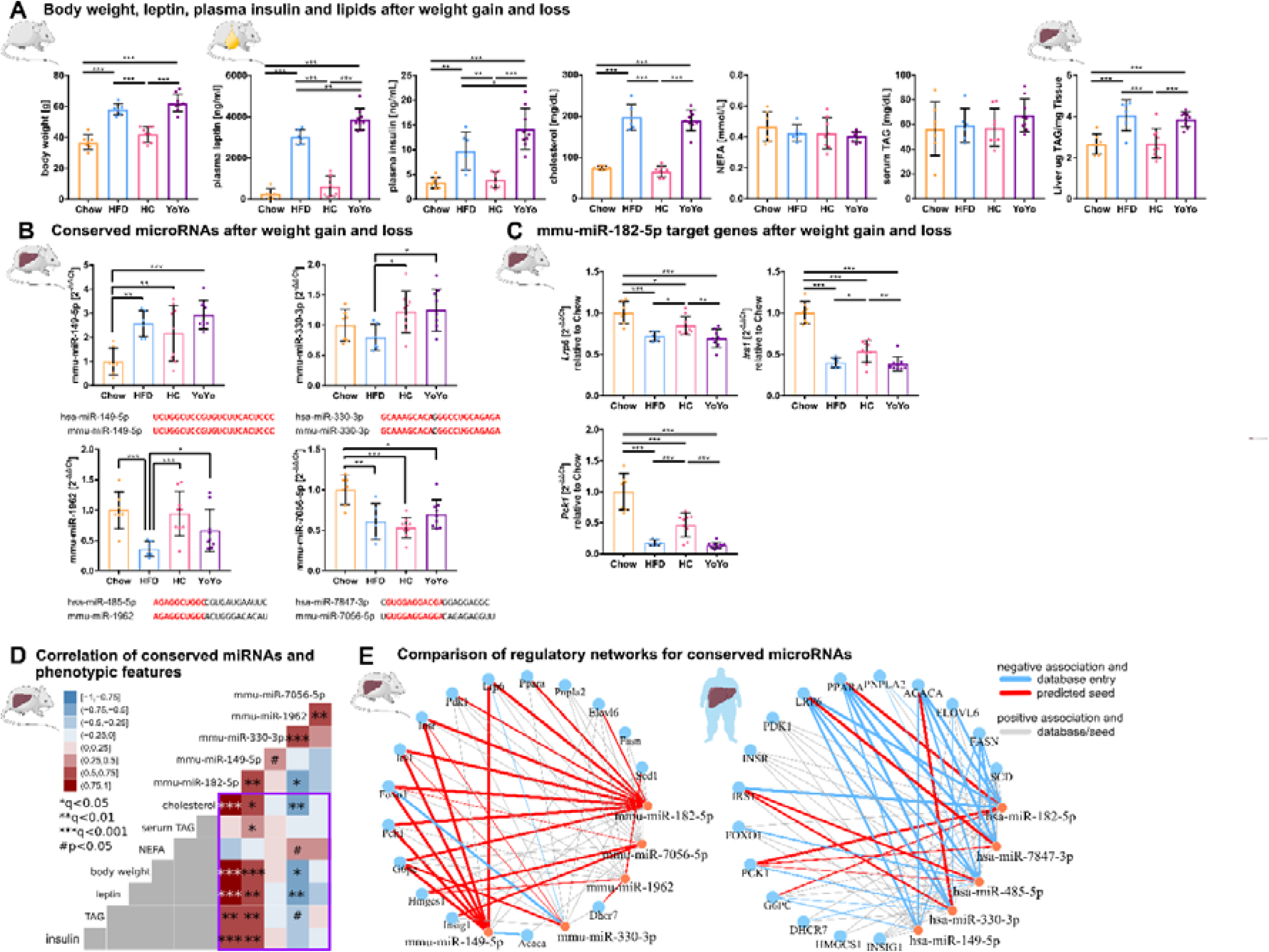
Expression and phenotypic association of conserved metabolic microRNAs and target genes in murine liver during weight cycling with a comparison between predicted murine and human regulatory gene networks. (**A**) Weight cycling in mice over 24 weeks induces changes in body weight that are associated with alterations in plasma markers and hepatic fat content (n=7 Chow, n=6 HFD, n=8 HC, n=9 YoYo). (**B**) Only mmu-miR-149-5p follows the expected expression pattern of overexpression in HFD and reversal during weight cycling as observed in human T2D, whereby the other three conserved miRNAs are reduced after HFD feeding. The miRNAs miR-149-5p and miR-330-3p are expressed from conserved genes, whereby mmu-miR-1962 and mmu-miR-7056-5p constitute orthologues to human hsa-miR-485-5p and hsa-miR-7847-3p respectively based on seed matches. Shared sequences are indicated in red. (**C**) Hepatic expression of *Lrp6*, *Irs1* and *Pck1* is significantly altered by weight cycling in obese mice. (**D**) Mmu-miR-182-5p shows the strongest positive associations to hepatic triacylglycerol (TAG), plasma leptin and insulin, body weight and cholesterol in the weight cycling cohort. mmu-miR-149-5p is further associated to serum TAG, which is also observed for human hepatic hsa-miR-149-5p expression. (**E**) Direct comparison between murine and human miRNA-target gene networks reveals a highly conserved and connected network with every metabolic miRNA potentially targeting several genes. The expression values of the murine network are based on qPCR measurements from the weight cycling cohort and the expression values of the human network are based on miRNA microarray and target gene qPCR measurements of n=40 individuals. Data are shown as scatter dot plots with mean ± SD (A-C), correlation matrices (D) or interaction network (E). Corrected for multiple testing: ***q<0.001, **q<0.01, *q<0.05, significant prior to adjustment: #p<0.05 (A-D); One-Way ANOVA (A-C) or Pearson’s correlation (D).

**Supplementary Fig. 6:**
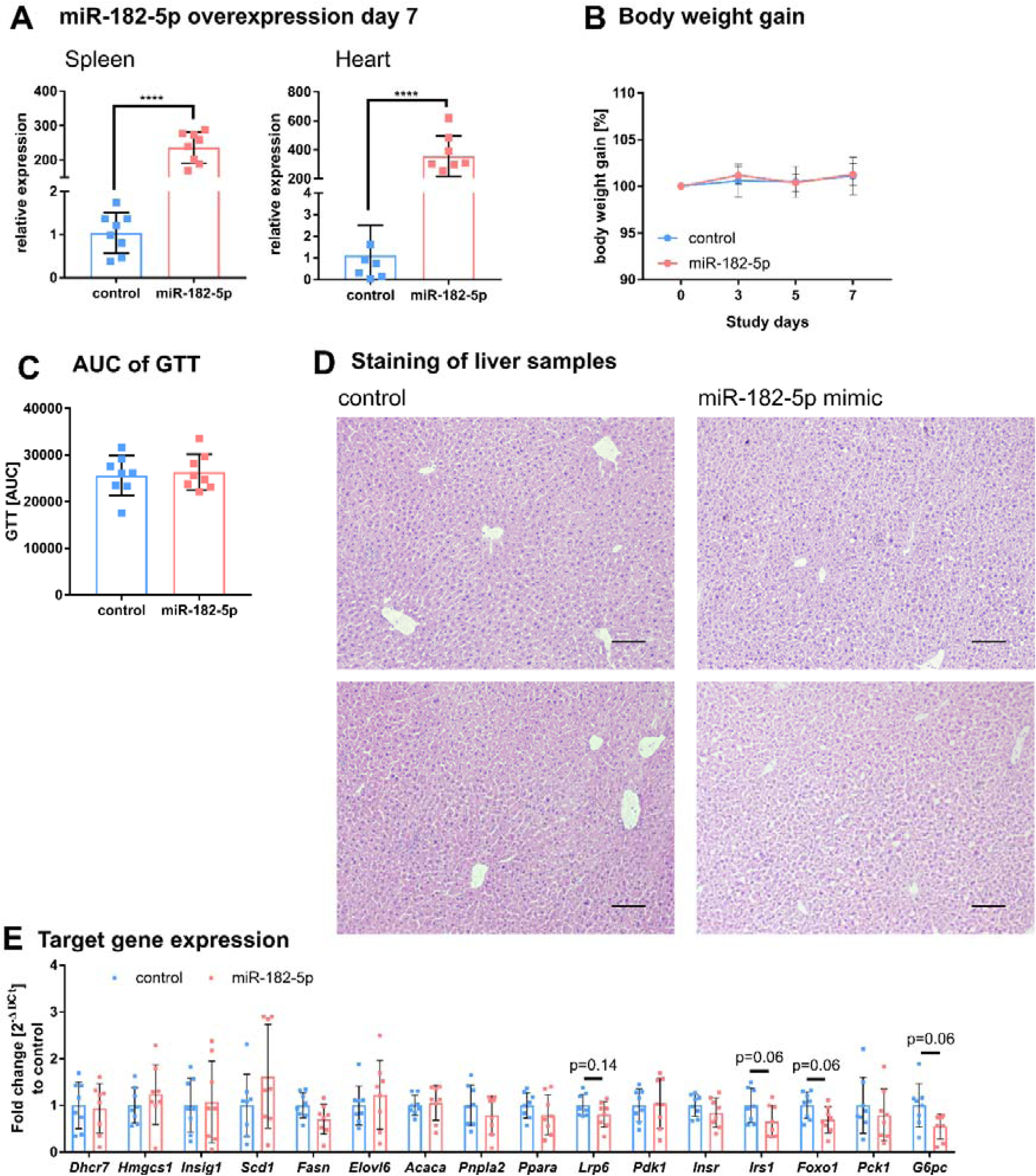
Additional phenotypic characteristics of mice overexpressing of miR-182-5p in liver. (**A**) Detection of miR-182-5p after tail vain injection in spleen and heart is significantly increased. (**B**) Body weight gain and (**C**) area under the curve (AUC) of the glucose tolerance test (GTT) did not differ between control and miR-182-5p treated mice (n=8 per group). (**D**) HE liver staining of two representative control and mimic treated animals. Mice overexpressing miR-182-5p show more lipid accumulations. Scale bar represents 100 µm. (**E**) miR-182-5p target gene expression was not changed after one week of acute miRNA overexpression, but *Lrp6*, *Irs1*, *Foxo1* and *G6pc* showed a strong tendency to reduced expression. Data are shown as mean ± SD (B) or scatter dot plots with mean ± SD (A,C,E).

### 6.2. Supplemental Table legends

**Supplementary Table 1: Clinical Characteristics for the complete (n=85) and the microarray (n=40) human liver cohort.** A rank sum test was performed to find significant differences between both subgroups.

**Supplementary Table 2: Primer sequences for SYBR-green analysis of cDNA by qPCR and cloning**

**Supplementary Table 3: Assay-IDs for TaqMan^TM^ Gene-expression Assays and TaqMan™ Advanced miRNA Assays**

**Supplementary Table 4: Associations between miRNAs and T2D.** Logistic regression models for the incidence of T2D were generated using self-written scripts in MATLAB. Age, sex, BMI and the NAFLD activity score (NAS) were used as additional cofactors. The coefficient estimate and p-value is indicated for the respective miRNA expression. ID is the respective identifier for the human microRNA on the GeneChip^TM^ miRNA 4.0 Array.

**Supplementary Table 5: Associations between miRNAs and metabolic traits.** Effect sizes describe the change of the trait if the miRNA expression changes by 1 log2 value. Age, sex, BMI and the NAFLD activity score (NAS) were used as additional cofactors whenever they did not serve as response value. ID is the respective identifier for the human microRNA on the GeneChip^TM^ miRNA 4.0 Array.

**Supplementary Table 6: Associations between miRNAs and excluding factors age and BMI.** To control for associations with both confounding factors, also the HbA1c level was considered as cofactor for linear regression models. Effect sizes describe the change of the trait if the miRNA expression changes by 1 log2 value. ID is the respective identifier for the human microRNA on the GeneChip^TM^ miRNA 4.0 Array.

**Supplementary Table 7: Predicted target genes from miRTarBase used for pathway analysis.**

**Supplementary Table 8: Candidate genes for hsa-mR-182-5p from database entries stratified by metabolic pathways from Gene Ontology.** Additional column qPCR contains target genes which were considered for analysis by qPCR in the complete cohort.

**Supplementary Table 9: Result of all consulted target gene databases and a comprehensive 3’UTR screening for each potential target gene of hsa-miR-182-5p.** Besides the relative position of the seed sequences within the 3’-UTR of the target gene, also the relative AU content of the surrounding 60 nt, whether there is an additional base pairing between the mRNA and the miRNA 3’ of the seed and the relative position of the seed within the target 3’UTR is listed.

## Notes

https://www.ncbi.nlm.nih.gov/geo/query/acc.cgi?acc=GSE176025

https://www.ncbi.nlm.nih.gov/geo/query/acc.cgi?acc=GSE211367

### Competing Interest Statement

The authors have declared no competing interest.

### Summary of Updates

Performed H&E Stainings; performed qPCRs to show off-target effects of miR-182-5p mimics in vivo; Simplified fig. 5; include original blots

## References

[1] Blüher M. Obesity: global epidemiology and pathogenesis. Nat Rev Endocrinol 2019;15:288–98. 10.1038/s41574-019-0176-8.

[2] Narayan KMV, Boyle JP, Thompson TJ, Gregg EW, Williamson DF. Effect of BMI on lifetime risk for diabetes in the U.S. Diabetes Care 2007;30:1562–6. 10.2337/dc06-2544.

[3] Chobot A, Górowska-Kowolik K, Sokołowska M, Jarosz-Chobot P. Obesity and diabetes-Not only a simple link between two epidemics. Diabetes Metab Res Rev 2018;34:e3042. 10.1002/dmrr.3042.

[4] Ling C, Groop L. Epigenetics: A Molecular Link Between Environmental Factors and Type 2 Diabetes. Diabetes 2009;58:2718–25. 10.2337/db09-1003.

[5] Friedman RC, Farh KK-H, Burge CB, Bartel DP. Most mammalian mRNAs are conserved targets of microRNAs. Genome Res 2009;19:92–105. 10.1101/gr.082701.108.

[6] Gurwitz D. Exosomal MicroRNAs in Tissue Crosstalk. Drug Development Research 2015;76:259–62. 10.1002/ddr.21264.

[7] Ambros V. microRNAs: Tiny Regulators with Great Potential. Cell 2001;107:823–6. 10.1016/S0092-8674(01)00616-X.

[8] Agbu P, Carthew RW. MicroRNA-mediated regulation of glucose and lipid metabolism. Nature Reviews Molecular Cell Biology 2021:1–14. 10.1038/s41580-021-00354-w.

[9] Rines AK, Sharabi K, Tavares CDJ, Puigserver P. Targeting hepatic glucose metabolism in the treatment of type 2 diabetes. Nat Rev Drug Discov 2016;15:786–804. 10.1038/nrd.2016.151.

[14] Trajkovski M, Hausser J, Soutschek J, Bhat B, Akin A, Zavolan M, et al. MicroRNAs 103 and 107 regulate insulin sensitivity. Nature 2011;474:649–53. 10.1038/nature10112.

[15] Kornfeld J-W, Baitzel C, Könner AC, Nicholls HT, Vogt MC, Herrmanns K, et al. Obesity-induced overexpression of miR-802 impairs glucose metabolism through silencing of Hnf1b. Nature 2013;494:111–5. 10.1038/nature11793.

[16] Ding J, Li M, Wan X, Jin X, Chen S, Yu C, et al. Effect of miR-34a in regulating steatosis by targeting PPARα expression in nonalcoholic fatty liver disease. Sci Rep 2015;5:13729. 10.1038/srep13729.

[17] Yang YM, Seo SY, Kim TH, Kim SG. Decrease of microRNA-122 causes hepatic insulin resistance by inducing protein tyrosine phosphatase 1B, which is reversed by licorice flavonoid. Hepatology 2012;56:2209–20. 10.1002/hep.25912.

[18] Li S, Chen X, Zhang H, Liang X, Xiang Y, Yu C, et al. Differential expression of microRNAs in mouse liver under aberrant energy metabolic status. J Lipid Res 2009;50:1756–65. 10.1194/jlr.M800509-JLR200.

[19] Yang W-M, Jeong H-J, Park S-W, Lee W. Obesity-induced miR-15b is linked causally to the development of insulin resistance through the repression of the insulin receptor in hepatocytes. Mol Nutr Food Res 2015;59:2303–14. 10.1002/mnfr.201500107.

[20] Yang W-M, Min K-H, Lee W. MicroRNA expression analysis in the liver of high fat diet-induced obese mice. Data in Brief 2016;9:1155–9. 10.1016/j.dib.2016.11.081.

[21] Gerin I, Clerbaux L-A, Haumont O, Lanthier N, Das AK, Burant CF, et al. Expression of miR-33 from an SREBP2 intron inhibits cholesterol export and fatty acid oxidation. J Biol Chem 2010;285:33652–61. 10.1074/jbc.M110.152090.

[22] Krause C, Geißler C, Tackenberg H, El Gammal AT, Wolter S, Spranger J, et al. Multi-layered epigenetic regulation of IRS2 expression in the liver of obese individuals with type 2 diabetes. Diabetologia 2020;63:2182–93. 10.1007/s00125-020-05212-6.

[23] Krützfeldt J, Rajewsky N, Braich R, Rajeev KG, Tuschl T, Manoharan M, et al. Silencing of microRNAs in vivo with ‘antagomirs.’ Nature 2005;438:685–9. 10.1038/nature04303.

[24] Chou C-H, Shrestha S, Yang C-D, Chang N-W, Lin Y-L, Liao K-W, et al. miRTarBase update 2018: a resource for experimentally validated microRNA-target interactions. Nucleic Acids Res 2018;46:D296–302. 10.1093/nar/gkx1067.

[25] Karagkouni D, Paraskevopoulou MD, Chatzopoulos S, Vlachos IS, Tastsoglou S, Kanellos I, et al. DIANA-TarBase v8: a decade-long collection of experimentally supported miRNA–gene interactions. Nucleic Acids Research 2018;46:D239–45. 10.1093/nar/gkx1141.

[26] Bartel DP. MicroRNAs: Target Recognition and Regulatory Functions. Cell 2009;136:215–33. 10.1016/j.cell.2009.01.002.

[27] Peterson SM, Thompson JA, Ufkin ML, Sathyanarayana P, Liaw L, Congdon CB. Common features of microRNA target prediction tools. Front Genet 2014;5. 10.3389/fgene.2014.00023.

[28] Grimson A, Farh KK-H, Johnston WK, Garrett-Engele P, Lim LP, Bartel DP. MicroRNA Targeting Specificity in Mammals: Determinants beyond Seed Pairing. Molecular Cell 2007;27:91–105. 10.1016/j.molcel.2007.06.017.

[29] Huynh C, Segura MF, Gaziel-Sovran A, Menendez S, Darvishian F, Chiriboga L, et al. Efficient in vivo microRNA targeting of liver metastasis. Oncogene 2011;30:1481–8. 10.1038/onc.2010.523.

[30] Davalos A, Goedeke L, Smibert P, Ramirez CM, Warrier NP, Andreo U, et al. miR-33a/b contribute to the regulation of fatty acid metabolism and insulin signaling. Proceedings of the National Academy of Sciences 2011;108:9232–7. 10.1073/pnas.1102281108.

[31] Hanin G, Yayon N, Tzur Y, Haviv R, Bennett ER, Udi S, et al. miRNA-132 induces hepatic steatosis and hyperlipidaemia by synergistic multitarget suppression. Gut 2018;67:1124–34. 10.1136/gutjnl-2016-312869.

[32] Dambal S, Shah M, Mihelich B, Nonn L. The microRNA-183 cluster: the family that plays together stays together. Nucleic Acids Res 2015;43:7173–88. 10.1093/nar/gkv703.

[33] Leti F, Malenica I, Doshi M, Courtright A, Van Keuren-Jensen K, Legendre C, et al. High-throughput sequencing reveals altered expression of hepatic microRNAs in nonalcoholic fatty liver disease-related fibrosis. Transl Res 2015;166:304–14. 10.1016/j.trsl.2015.04.014.

[34] Dolganiuc A, Petrasek J, Kodys K, Catalano D, Mandrekar P, Velayudham A, et al. MicroRNA expression profile in Lieber-DeCarli diet-induced alcoholic and methionine choline deficient diet-induced nonalcoholic steatohepatitis models in mice. Alcohol Clin Exp Res 2009;33:1704–10. 10.1111/j.1530-0277.2009.01007.x.

[35] Nie J, Li C, Li J, Chen X, Zhong X. Analysis of nonLJalcoholic fatty liver disease microRNA expression spectra in rat liver tissues. Molecular Medicine Reports 2018. 10.3892/mmr.2018.9268.

[36] Weyer C, Hanson RL, Tataranni PA, Bogardus C, Pratley RE. A high fasting plasma insulin concentration predicts type 2 diabetes independent of insulin resistance: evidence for a pathogenic role of relative hyperinsulinemia. Diabetes 2000;49:2094–101. 10.2337/diabetes.49.12.2094.

[37] Hayes CN, Chayama K. MicroRNAs as Biomarkers for Liver Disease and Hepatocellular Carcinoma. Int J Mol Sci 2016;17. 10.3390/ijms17030280.

[38] Jones A, Danielson KM, Benton MC, Ziegler O, Shah R, Stubbs RS, et al. miRNA signatures of insulin resistance in obesity. Obesity (Silver Spring) 2017;25:1734–44. 10.1002/oby.21950.

[39] Jiménez-Lucena R, Camargo A, Alcalá-Diaz JF, Romero-Baldonado C, Luque RM, Ommen B van, et al. A plasma circulating miRNAs profile predicts type 2 diabetes mellitus and prediabetes: from the CORDIOPREV study. Exp Mol Med 2018;50:1–12. 10.1038/s12276-018-0194-y.

[40] Weale CJ, Matshazi DM, Davids SFG, Raghubeer S, Erasmus RT, Kengne AP, et al. Circulating miR-30a-5p and miR-182-5p in Prediabetes and Screen-Detected Diabetes Mellitus. Diabetes Metab Syndr Obes 2020;13:5037–47. 10.2147/DMSO.S286081.

[41] Weale CJ, Matshazi DM, Davids SFG, Raghubeer S, Erasmus RT, Kengne AP, et al. Expression Profiles of Circulating microRNAs in South African Type 2 Diabetic Individuals on Treatment. Frontiers in Genetics 2021;12.

[42] Szilágyi M, Pös O, Márton É, Buglyó G, Soltész B, Keserű J, et al. Circulating Cell-Free Nucleic Acids: Main Characteristics and Clinical Application. Int J Mol Sci 2020;21:6827. 10.3390/ijms21186827.

[43] Endzeliņš E, Berger A, Melne V, Bajo-Santos C, Soboļevska K, Ābols A, et al. Detection of circulating miRNAs: comparative analysis of extracellular vesicle-incorporated miRNAs and cell-free miRNAs in whole plasma of prostate cancer patients. BMC Cancer 2017;17:730. 10.1186/s12885-017-3737-z.

[44] Garcia-Martin R, Wang G, Brandão BB, Zanotto TM, Shah S, Kumar Patel S, et al. MicroRNA sequence codes for small extracellular vesicle release and cellular retention. Nature 2022;601:446–51. 10.1038/s41586-021-04234-3.

[45] Krause C, Sievert H, Geißler C, Grohs M, El Gammal AT, Wolter S, et al. Critical evaluation of the DNA-methylation markers ABCG1 and SREBF1 for Type 2 diabetes stratification. Epigenomics 2019;11:885–97. 10.2217/epi-2018-0159.

[46] Thompson MD, Monga SPS. WNT/beta-catenin signaling in liver health and disease. Hepatology 2007;45:1298–305. 10.1002/hep.21651.

[47] Tamai K, Semenov M, Kato Y, Spokony R, Liu C, Katsuyama Y, et al. LDL-receptor-related proteins in Wnt signal transduction. Nature 2000;407:530–5. 10.1038/35035117.

[48] Go G. Low-Density Lipoprotein Receptor-Related Protein 6 (LRP6) Is a Novel Nutritional Therapeutic Target for Hyperlipidemia, Non-Alcoholic Fatty Liver Disease, and Atherosclerosis. Nutrients 2015;7:4453–64. 10.3390/nu7064453.

[49] Mani A, Radhakrishnan J, Wang H, Mani A, Mani M-A, Nelson-Williams C, et al. LRP6 mutation in a family with early coronary disease and metabolic risk factors. Science 2007;315:1278–82. 10.1126/science.1136370.

[50] Liu H, Fergusson MM, Wu JJ, Rovira II, Liu J, Gavrilova O, et al. Wnt Signaling Regulates Hepatic Metabolism. Science Signaling 2011;4:ra6–ra6. 10.1126/scisignal.2001249.

[51] Ye Z, Go G-W, Singh R, Liu W, Keramati AR, Mani A. LRP6 protein regulates low density lipoprotein (LDL) receptor-mediated LDL uptake. J Biol Chem 2012;287:1335–44. 10.1074/jbc.M111.295287.

[52] Go G-W, Srivastava R, Hernandez-Ono A, Gang G, Smith SB, Booth CJ, et al. The combined hyperlipidemia caused by impaired Wnt-LRP6 signaling is reversed by Wnt3a rescue. Cell Metab 2014;19:209–20. 10.1016/j.cmet.2013.11.023.

[53] Keramati AR, Singh R, Lin A, Faramarzi S, Ye Z, Mane S, et al. Wild-type LRP6 inhibits, whereas atherosclerosis-linked LRP6R611C increases PDGF-dependent vascular smooth muscle cell proliferation. Proc Natl Acad Sci USA 2011;108:1914–8. 10.1073/pnas.1019443108.

[54] Wang S, Song K, Srivastava R, Dong C, Go G-W, Li N, et al. Nonalcoholic fatty liver disease induced by noncanonical Wnt and its rescue by Wnt3a. FASEB J 2015;29:3436–45. 10.1096/fj.15-271171.

[55] Singh R, De Aguiar RB, Naik S, Mani S, Ostadsharif K, Wencker D, et al. LRP6 Enhances Glucose Metabolism by Promoting TCF7L2-dependent Insulin Receptor Expression and IGF Receptor stabilization in Humans. Cell Metab 2013;17:197–209. 10.1016/j.cmet.2013.01.009.

[56] Bommer GT, Feng Y, Iura A, Giordano TJ, Kuick R, Kadikoy H, et al. IRS1 Regulation by Wnt/β-Catenin Signaling and Varied Contribution of IRS1 to the Neoplastic Phenotype. J Biol Chem 2010;285:1928–38. 10.1074/jbc.M109.060319.

