## Supplementary figures and images for "Liver microRNA transcriptome reveals miR-182 as link between type 2 diabetes and fatty liver disease in obesity"

### Supplementary Figure 1

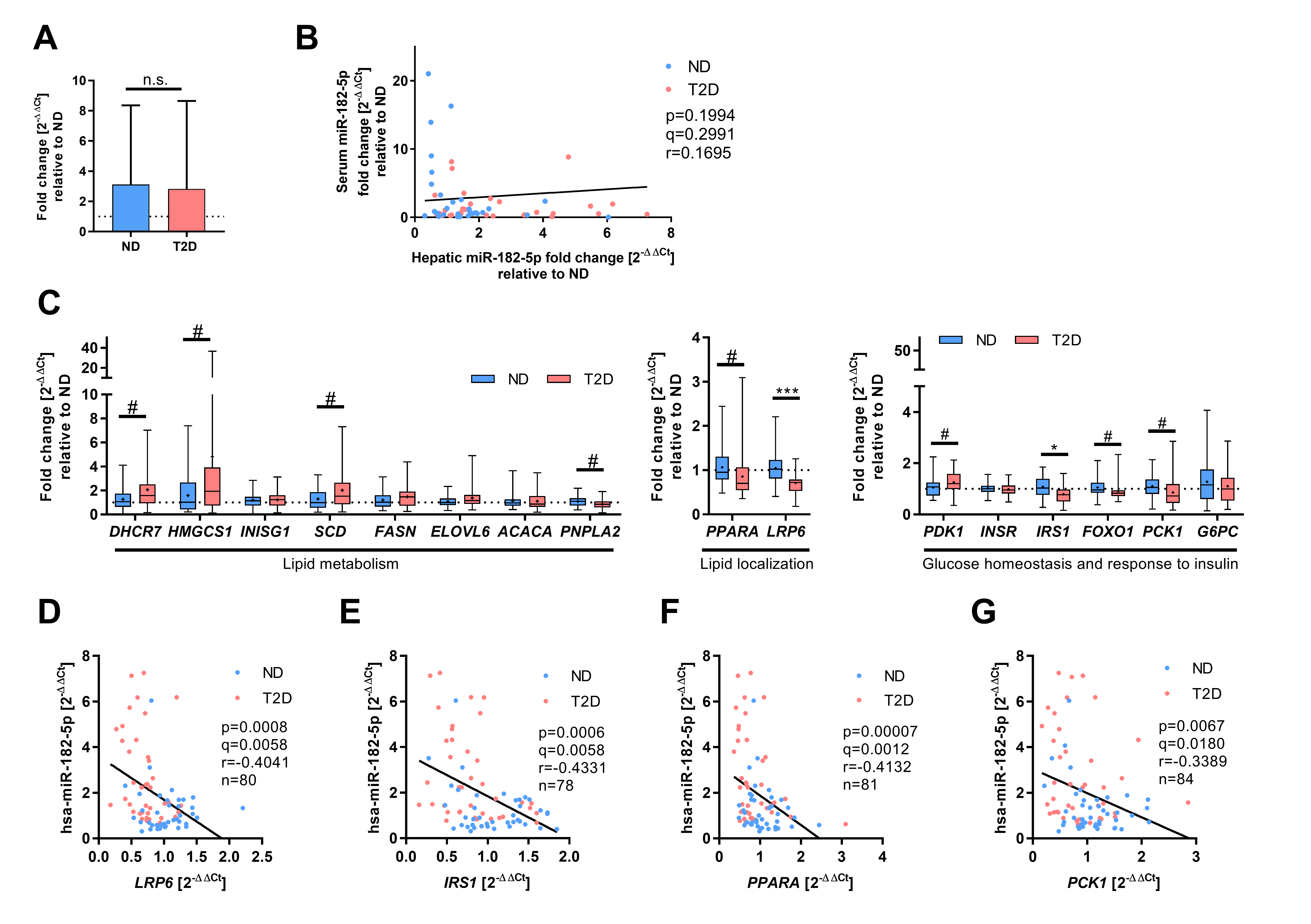

### Supplementary Figure 2

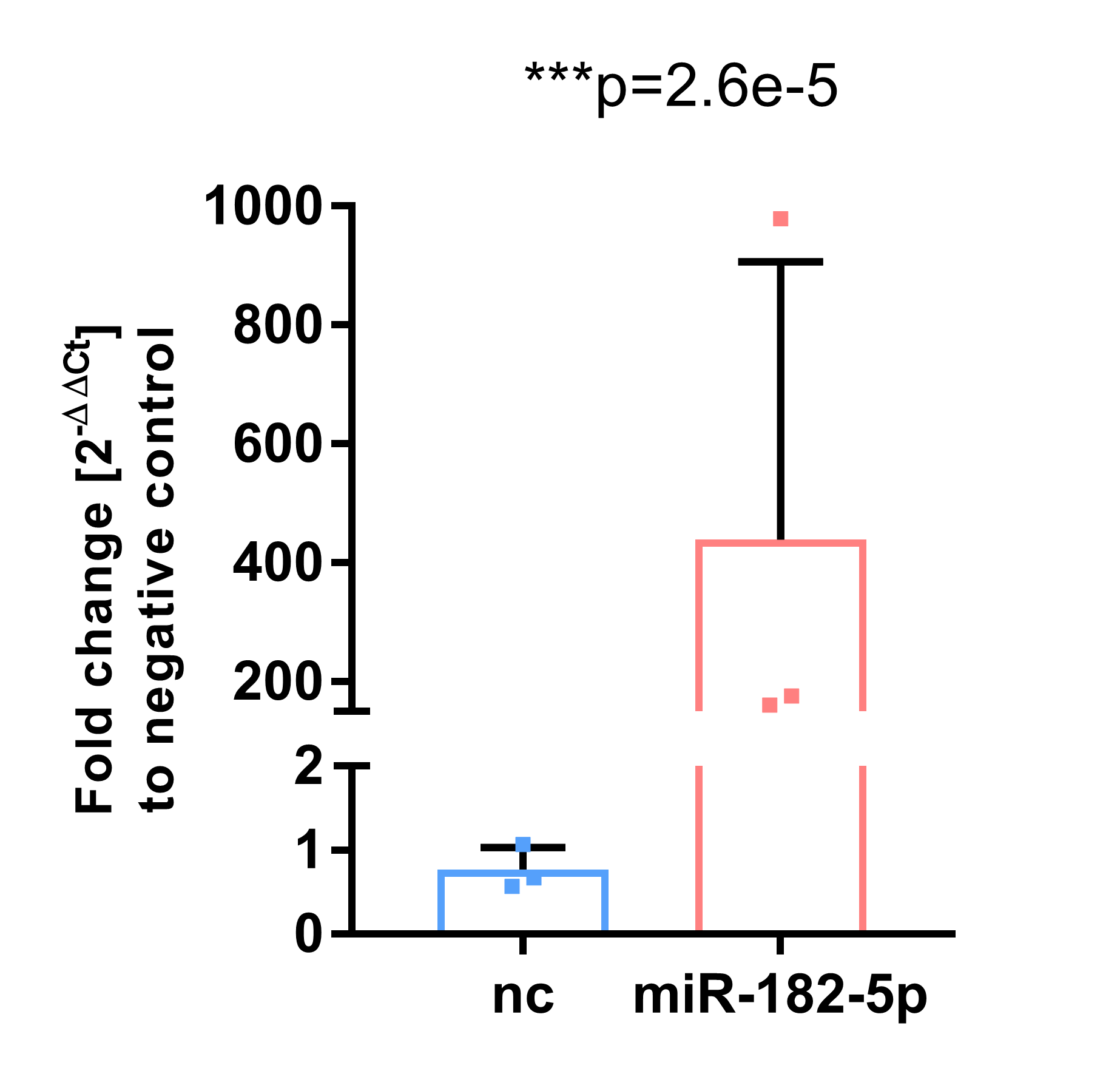

### Supplementary Figure 3

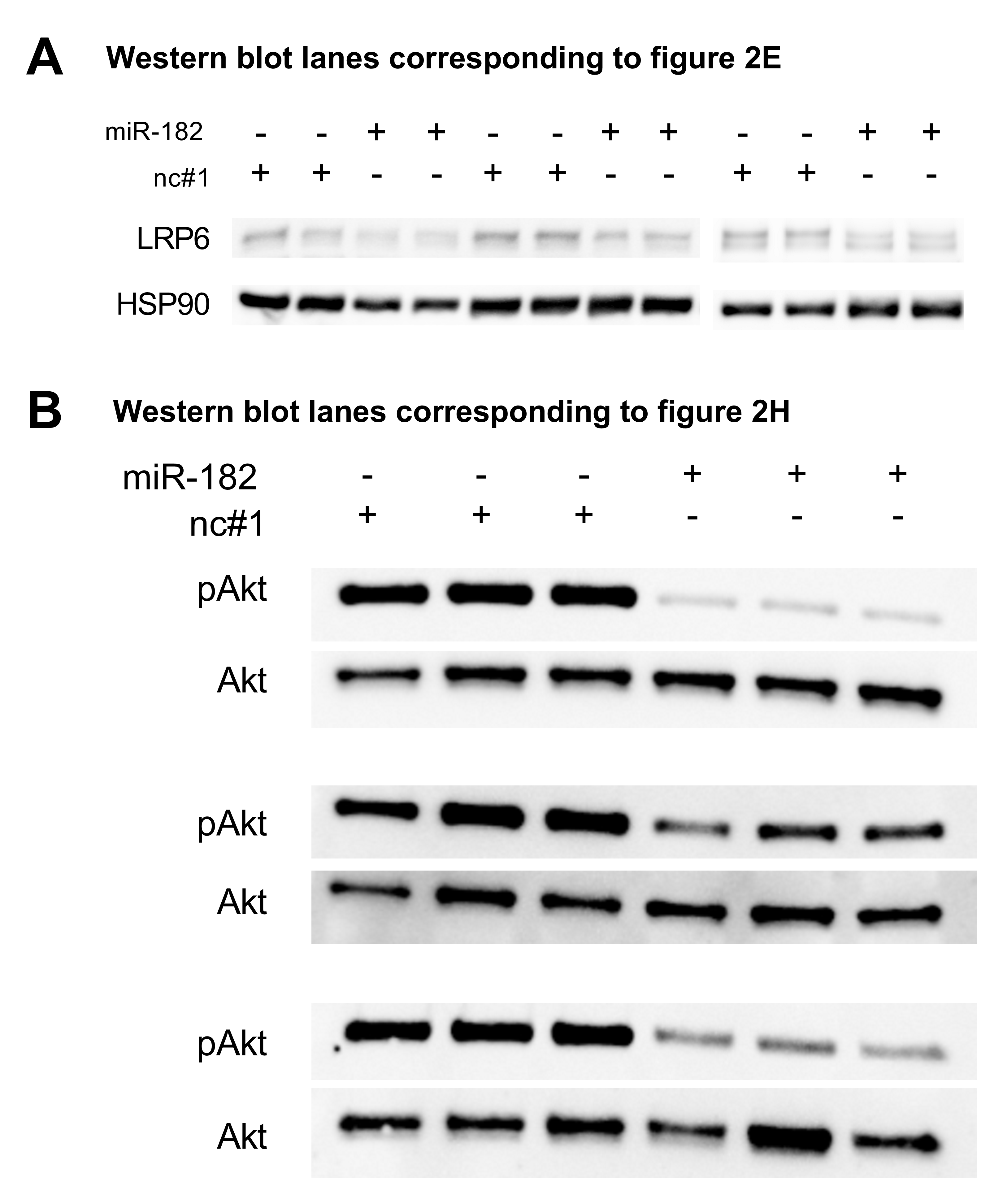

### Supplementary Figure 4

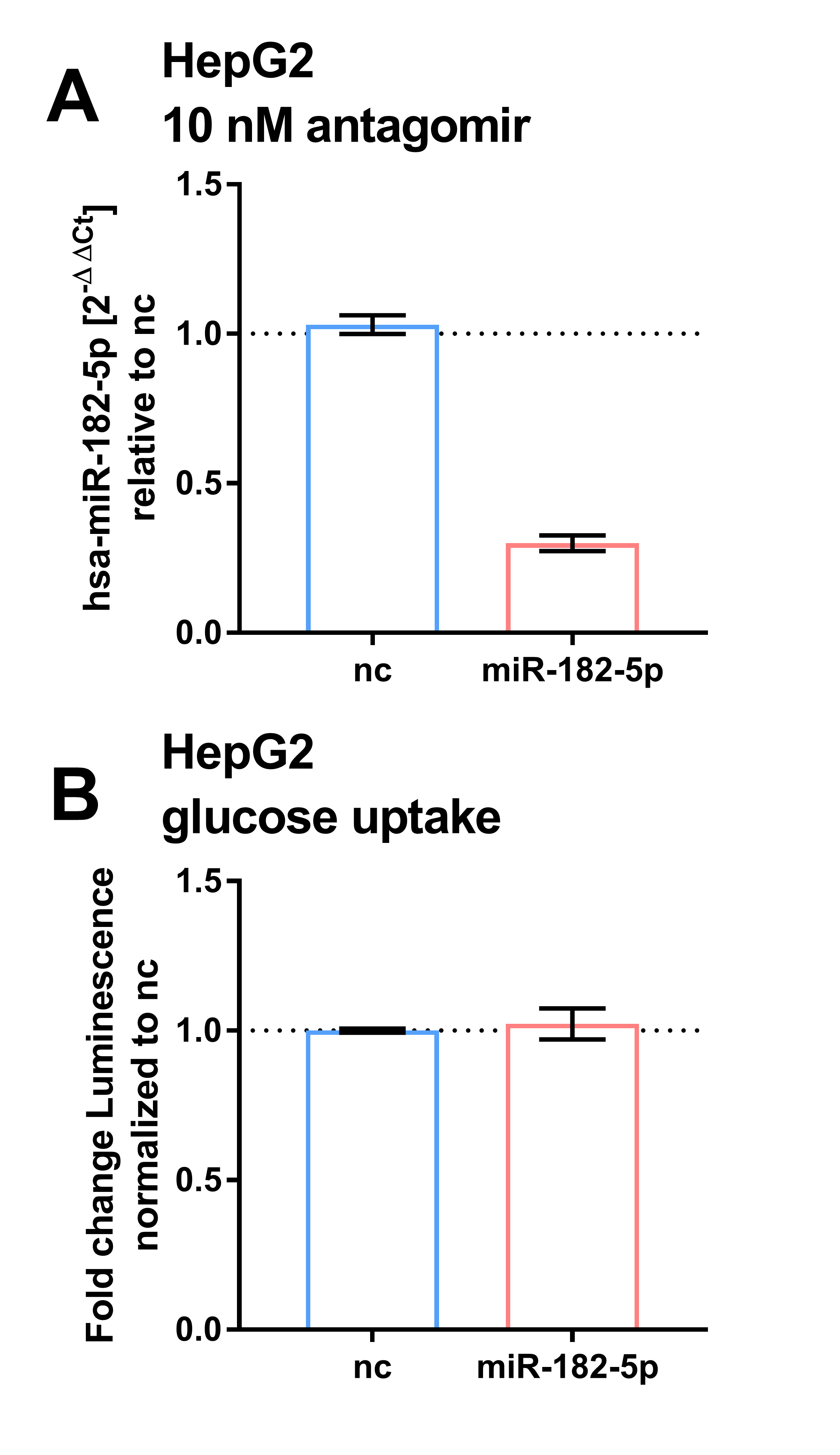

### Supplementary Figure 5

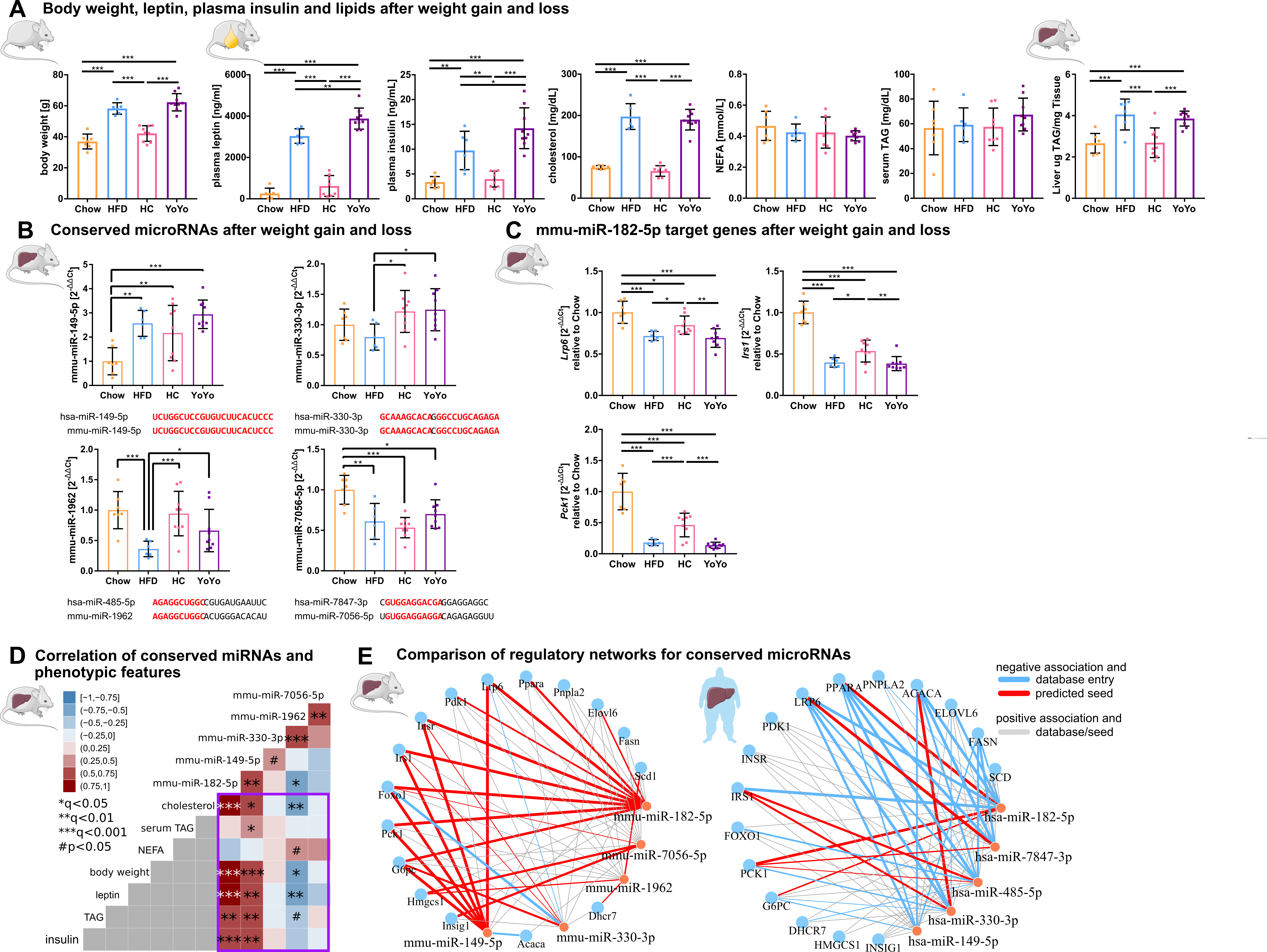

### Supplementary Figure 6

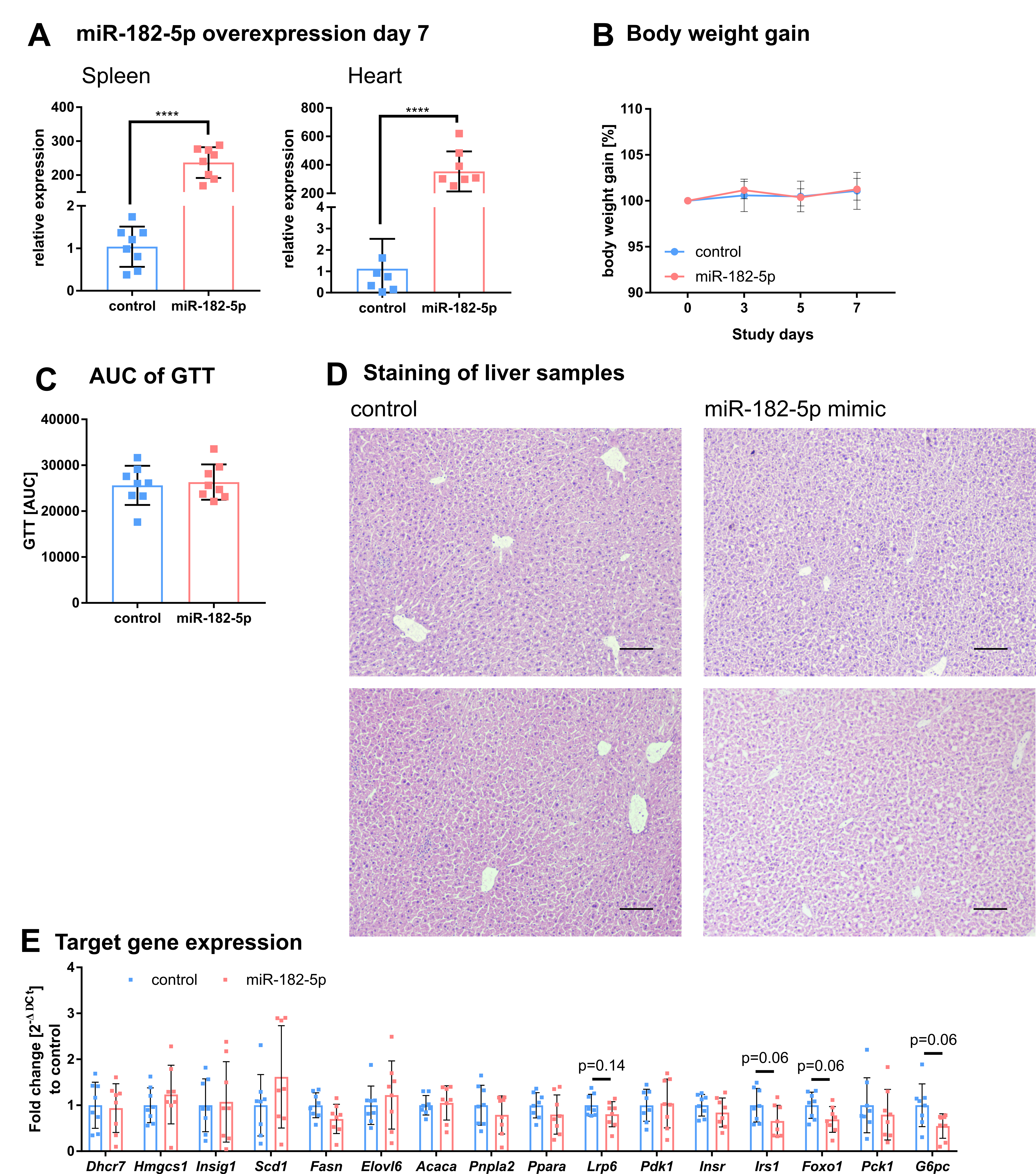
