## Supplemental Methods for "Liver microRNA transcriptome reveals miR-182 as link between type 2 diabetes and fatty liver disease in obesity"

### Supplemental Material

#### Supplemental Methods

##### Human cohort:

All clinical parameters were measured at the Institut für Klinische Chemie und Laboratoriumsmedizin, Zentrum für Diagnostik, Universitätsklinikum Eppendorf, Hamburg, Germany, according to the DIN EN ISO 15189:2014 certification. Serum glucose, cholesterol, HDL and triacylglycerols were determined using photometric assays (kinetic bichromatic analysis; in-house assay). HbA1c was quantified by capillary electrophoresis or by turbidimetric inhibition assays (in-house assay). Insulin levels were quantified from frozen serum samples using the Human/Canine/Porcine Insulin Quantikine ELISA Kit (R&D Systems, Inc, Minneapolis, US). The NAFLD activity score (NAS) and degree of hepatic steatosis was determined during surgery according to the current recommendations by two expert pathologists. Anti-diabetic treatment was not uniform in the T2D group and included oral glucose-lowering medications or insulin. The mean BMI was 52.37 kg/m$^2$ ranging from 32.19 to 84.87 kg/m$^2$. The complete cohort was imbalanced in age, which was treated as potential confounding factor and adjusted for if applicable.

##### Mouse Models

At the end of the diet-intervention study, all mice were fasted for 4 hours at the start of the light phase before being sacrificed by cervical dislocation for organ withdrawal. To obtain plasma blood was collected in EDTA-containing tubes and centrifuged at 4°C and 2000 x g for 10 min.

##### RNA isolation and gene expression analysis in human liver

Whole cell RNA was extracted from 25 mg of snap frozen liver using the miRNeasy mini kit (QIAGEN, Hilden, Germany) and quantified spectrometrically. 2 µg of RNA was reverse transcribed into cDNA using the SuperScript VILO cDNA synthesis kit (Invitrogen, Carlsbad, US).

##### Array-based miRNA transcriptome analysis

500 ng RNA from 40 subjects (n=20 type-2-diabetic, n=20 non-diabetic) and 1000 ng RNA from n=6 mice kept on HFD for 28 weeks were used for probe preparation by the FlashTag™ Biotin HSR RNA Labelling Kit (Applied Biosystems, Foster City, US). Prepared samples and control oligo B2 from the

GeneChip™ Hybridization Control Kit (Applied Biosystems, Foster City, US) were hybridized to the GeneChip™ miRNA 4.0 Array (Applied Biosystems, Foster City, US). After incubation for 18-h and washing, each array was scanned by a GeneChip™ Scanner 3000 (Applied Biosystems, Foster City, US).

**RNA isolation and gene expression analysis of murine liver samples**

Whole cell RNA was extracted from 20 mg of snap frozen liver using the miRNeasy mini kit (QIAGEN, Hilden, Germany) and quantified spectrometrically. 1 µg of RNA was reverse transcribed using the QuantiTect Reverse Transcription kit (QIAGEN, Hilden, Germany). Expression of Acaca, Dhcr7, Elovl6, Fasn, Foxo1, G6pc, Hmgcs1, Insig1, Insr, Irs1, Scd1, Lrp6, Pck1, Pdk1, Pnpla2 and Ppara was measured in duplicates by SYBR green qPCR (SYBR Green Master Mix, Thermo Fisher Scientific, Inc., Rockford, US) and calculated with the ΔΔCt method using Hprt1 as housekeeping gene. Primer sequences can be found in the supplementary material.

**microRNA expression analysis in human and murine liver**

1 µg total human or mouse liver RNA was reversely transcribed with the qScript microRNA cDNA synthesis kit (Quanta Bioscience, Beverly, US). Self-designed forward and a pre-designed universal reverse primers were used for the expression measurement of miR-182-5p by SYBR green qPCR (PerfeCTa SYBR Green SuperMix, Quanta Bioscience, Beverly, US) as duplicates and expression values were calculated with the ΔΔCt method using hsa-miR-24-3p as housekeeping gene. Primer sequences can be found in the supplementary material.

**miRNA isolation and miRNA expression analysis in human serum**

Serum miRNA was extracted from 200 µl frozen serum by the miRNeasy Serum/Plasma Advanced kit (QIAGEN, Hilden, Germany) using the spike-in control cel-miR-39-3p (QIAGEN, Hilden, Germany). MiRNAs from 5 µl eluate (corresponding to 40 µl serum input) were reverse transcribed by qScript microRNA cDNA synthesis kit (Quanta Biosciences, Beverly, US). Expression was quantified as mentioned before. Each qPCR reaction contained 266 nl input serum. Target expression was normalized to the spike-in control cel-miR-39-3p (cel-miR-39-3p 5'-CACCGGGTGTAAATCAGCTTG) expression using the ΔΔCt method.

**Plasmid cloning, site-directed mutagenesis and luciferase reporter gene assay**

For plasmid cloning, the 3'-UTR of *LRP6* containing a potential 7mer-m8 recognition site was generated by PCR amplification using human hepatic cDNA with primer containing the recognition motifs for *NotI* and *XhoI* restriction.

After restriction digestion with *NotI* and *XhoI* (NEB, Ipswich, US) the PCR fragment was directionally cloned downstream of the Renilla luciferase ORF of the psiCHECK-2 vector (Promega, Madison, US) using the Quick Ligase protocol (M2200, NEB, Ipswich, US). A constitutively expressed Firefly luciferase gene on the same vector served for normalization of the Renilla luciferase signal. 10 ng of this plasmid served as PCR template for a site-directed mutagenesis whereby the central nucleotide of the seed sequence (5'-TTGGCAA) was exchanged by an adenosine to (5'-TTGACAA). The site-directed mutagenesis reaction was performed by using the Phusion Hot Start II High-Fidelity PCR Master Mix (Thermo Scientific, Waltham, US) and a PCR protocol for 12 cycles. The parental strand was digested by *DpnI* for 15 min (NEB, Ipswich, US). All primer sequences can be found in the supplementary material.

Plasmids were transformed in DH5α *E. coli* bacteria (NEB, Ipswich, US) and extracted with the QIAprep Spin Midiprep kit (QIAGEN, Hilden, Germany). HEK-293 cells (obtained from ATCC, Manassas, US, Mycoplasma-free) were cultivated at 37 °C and 5 % $CO_2$ in high glucose (4.5 g/l) DMEM medium supplemented with 10 % (vol./vol.) FBS and 1 % penicillin/streptomycin.

HEK-293 cells were co-transfected in triplicates with miRNA precursor mimics (pri-miRNA-182-5p PM12369 and negative control #1 AM17110, Ambion, Applied Biosystems, Foster City, US) and the luciferase plasmid containing the 3'UTR with either the consensus seed or the mutated seed sequence generated by site-directed mutagenesis using a final concentration of 10 nmol/l for mimic or negative control and 50 ng plasmid per well of a 96-well plate. Transfection was performed by reverse transfection using Lipofectamine™ 3000 (Invitrogen, Carlsbad, US) and a cell concentration of 100.000 cells per 1 ml high glucose (4.5 g/l) DMEM medium supplemented with 10 % (vol./vol.) FBS. Cells were harvested after 48 h of incubation and lysed in 80 µl 1x passive lysis buffer (Dual-Glo® luciferase reporter gene assay, Promega, Madison, US). Renilla and Firefly luciferase signals were measured in 20 µl cell lysates by using the Dual-Glo® luciferase reporter gene assay (Promega, Madison, US). The Renilla signal was normalized to the Firefly signal which are both encoded on the same plasmid. The miR182-5p mimic signal was compared to the respective negative control in each experiment. Each experiment was performed three times.

**miRNA overexpression in HepG2 cells**

HepG2 (obtained from ATCC, Manassas, US, Mycoplasma-free) were cultivated at 37 °C and 5 % $CO_2$ in low glucose (1.5 g/l) DMEM medium supplemented with 10 % (vol./vol.) FBS, 1 mM sodium pyruvate and 1 % penicillin/streptomycin. miRNA precursor mimics as mentioned before were reverse transfected into HepG2 cells in triplicates with a final concentration of 10 nmol/l per well of a 6-well plate using Lipofectamine RNAiMAX Transfection Reagent. Transfected cells were washed with PBS and lysed in 700 µl TRIzol after 48 h incubation. Whole cell RNA was isolated by miRNeasy mini kit (QIAGEN, Hilden, Germany). The quantification of HepG2 miR-182-5p (477935_mir) expression was performed by reverse transcription of 10 ng of HepG2 whole cell RNA with the TaqMan™ Advanced cDNA synthesis kit (Applied Biosystems, Foster City, US) using miR-24-3p (477992_mir) as housekeeper assay. For the quantification of potential target genes, 500 ng RNA was reversely transcribed into cDNA using the High-Capacity cDNA Reverse Transcription Kit (Applied Biosystems, Foster City, US). Gene expression measurement was performed as previously stated for hepatic genes. Negative control #1 transfected cells served as reference for comparison. For western blot analysis of the LRP6 protein, cells were harvested in ice-cold PBS after 72 h incubation. Total protein was isolated by incubation of the cell pellet for 30 min on ice in modified RIPA buffer (50 mmol/l Tris-HCl, 1% (vol./vol.) NP-40, 0.25 % (wt/vol.) Na-deoxycholat, 150 mmol/l NaCl, 1 mmol/l EDTA, 1 mmol/l PMSF, Na3VO4, 1 mmol/l NaF, supplemented with protease inhibitor cocktail cOmplete™ (Roche, Basel, CH)), subsequent centrifugation for 10 min at 14.000 g and 4 °C and the supernatant was collected. For western blot analysis for the measurement of phospho-Akt, transfected cells were treated with 20 nmol/l insulin (Lantus 100 E/ml, Sanofi, Paris, FR) for 10 min prior harvest. Each experiment was performed three times.

**Western Blot analysis**

25 µg (Akt/pAkt) to 35 µg (LRP6) of HepG2 protein lysate was separated for 1.5 h at 100 V by SDS-Page (7.5 % (wt/vol.) acrylamide, TGX Stain-Free™ FastCast™ Acrylamide Kit, Bio-Rad Laboratories, Inc., Hercules, US) and blotted on nitrocellulose membranes by fast blotting (mixed MW, Trans-Blot® Turbo™, Bio-Rad Laboratories, Inc., Hercules, US). For the analysis of LRP6 in murine liver, protein was extracted by homogenization in modified RIPA buffer and subsequent centrifugation for 20 min at 18,000 g and 4 °C. 20 µg protein lysate was separated by SDS-Page (7.5 % (wt/vol.) acrylamide Criterion TGX Precast Midi Protein gel, Bio-Rad Laboratories Inc, US) and blotted on 0.2 µm PVDF membranes (Bio-Rad Laboratories Inc, US). Total protein was imaged from the membrane as a control after UV-activation of the gel for 60 s. Unspecific binding sites were blocked by incubation for 1 h at room temperature in 5 % (wt/vol.) milk in TBS + 0.1 % (vol./vol.) Tween (TBST). The blots were incubated with primary antibodies against LRP6 (1:1000, EPR2423(2) ab134146 rabbit mAb, lot GR3256666-1, Abcam, Cambridge, UK) or HSP90 (1:1000, C45G5 rabbit mAb, lot 5, Cell Signalling Technology, Danvers, US) over night at 4 °C. After washing in TBST, the membrane was incubated for 1 h at room temperature with a secondary antibody (1:5000) conjugated with HRP (polyclonal goat anti-rabbit HRP, lot 20061231, Dako, Agilent, Santa Clara, US). For the analysis of phospho-Akt/Akt protein, the protein blots were first incubated with a primary antibody against phospho-Akt Ser473 (1:1000, D9E XP rabbit mAb #4060, lot 23, Cell Signaling Technology, Danvers, US) and after stripping of the membrane with a primary antibody against total Akt (1:1000, rabbit pAb #9272, lot 28, Cell Signaling Technology, Danvers, US). Each experiment was performed three times.

**Glucose uptake of HepG2 cells**

miR-182-5p precursor mimics and controls were reverse transfected in HepG2 cells as triplicates as described above. A glucose uptake assay (Glucose Uptake-Glo™ Assay, Promega, Madison, US) was performed in 6-well plates using a final concentration of 1 mM 2-deoxyglucose (2DG) in PBS for the uptake reaction. Prior incubation with 2DG, the cells were treated for 10 min with 20 nM insulin (Lantus 100 E/ml, Sanofi, Paris, FR). The luciferase signal was measured in duplicates for each sample in a 96-well format. The mimic signal was normalized to the respective cells treated with negative control. Each experiment was performed three times.

**In vivo overexpression of miR-182-5p in mouse liver**

Male C57BL/6J mice were purchased at 5-6 weeks of age from Janvier Labs (Saint-Berthevin, Cedex, France), maintained as described before and fed with HFD for four weeks prior and throughout the study. Based on their body weight, the mice were assigned at day -1 to either the control or the miR-182-5p mimic group so that each group contained eight weight-matched mice. On day 0, the first injection of 1 mg miR-182-5p-mimic or negative control per kg body weight was performed via tail vein using Invivofectamine® 3.0 (Invitrogen, Carlsbad, US) as described by the manufacturer. A second injection was performed at day 3.5 to maintain the microRNA level. Mice were phenotypically characterized by NMR on days -1 and 7 and a glucose tolerance test after 6 h fasting on day 5. Plasma was collected from blood samples during the GTT to determine insulin levels by ELISA (EZRMI-13K, EMD Millipore Corporation, Burlington, US). All mice were sacrificed at day 7 after a 3h fast. Liver, spleen and kidney were collected and snap frozen for the extraction of RNA and protein as described before. Hepatic triglycerides were measured using the Triglyceride Quantification colorimetric-/fluorometric Kit (MAK266, Sigma-Aldrich, St. Louis, US) from 50 mg of liquid-nitrogen pulverized tissue homogenized in 5 % (wt/vol.) IGEPAL® CA-630 (Sigma-Aldrich, St. Louis, US).

**Microarray statistics**

Regression analysis, statistical tests and visualization was performed by MATLAB R2020a (The MathWorks, Natick, US), R 3.5.1 (The R Foundation for Statistical Computing, Vienna, Austria) and GraphPad Prism 7.05 (GraphPad Software, Inc, San Diego, US).

Generated CEL data was imported into Transcriptome Analysis Console (TAC) 4.0 and pre-processed for further analysis in MATLAB R2020a and R 3.5.1. Array data was normalized by Robust Multi-chip Analysis (RMA) algorithm as indicated by the manufacturer, which contains background adjustment, log2 transformation and quantile normalization to increase the fold change ratios. Probesets were considered expressed if more than 50 % of probes have a significantly ($p<0.05$) higher detection signal than the background (DABG, detection above background).

Linear regression models for continuous responses (metabolic traits) or logistic regression models for the incidence of T2D were generated in MATLAB. Consequently, age, sex, BMI and the NAFLD activity score (NAS) as marker for hepatic steatosis and fibrosis were used as additional cofactors if not used as response variable:

Trait or T2D = $\beta_0 + \beta_1$*NAS + $\beta_2$*age + $\beta_3$*sex + $\beta_4$*BMI + $\beta_5$*log2$_{\mathrm{miRNA}}$

By using NAS as cofactor we ensured that NAFLD is not driving the altered miRNA expression in T2D.

To control for associations with confounding factors (age, BMI) also the HbA1c level was considered as cofactor:

Age or BMI = $\beta_0 + \beta_1$*HbA1c + $\beta_3$*sex + $\beta_4$*[BMI or age] + $\beta_5$*$\log2_{miRNA}$

Resulting effect sizes β describe the change of the trait if the miRNA expression changes by 1 log2 value.

A p-value<0.05 was considered associated. Since FDR adjustment of thereby generated q-values resulted in no significantly associated miRNA (q<0.05), another approach for candidate identification was used. First, we filtered for confidently expressed microRNAs by applying a log2 threshold of 2.3. This is reasoned by the minimal group mean log2 value which was declared as "is expressed" by the TAC software and which could be quantified by qPCR. Moreover, multiple associations to the incidence of type 2 diabetes or other metabolic traits and missing associations to BMI and age were mandatory to be declared as candidate gene for qPCR validation. Secondly, we evaluated which microRNAs are conserved between human and mouse by 7mer seed match analysis. By use of these exclusion criteria, we were able to identify candidate miRNAs which are differentially expressed in T2D independently of obesity and not driven by NAS.

**General statistics**

Changes in gene expression between two groups were tested by student's t-test and between multiple groups by One-way ANOVA with adjustment for multiple testing by controlling the FDR with the Benjamini-Hochberg procedure. Fold-change gene expression values were calculated by $2^{-\Delta\Delta Ct}$ relative to the control group (ND in humans and chow or control in mouse).

Luciferase signal, glucose uptake, changes in HepG2 cell gene expression or protein abundance were tested by student's t-test and adjusted for multiple testing by controlling the FDR with the Benjamini-Hochberg procedure if more than one test was performed. Metabolic parameters of the *in vivo* mouse study were tested by student's t-test.

All ΔCt-values which are not within a three standard deviations interval of all samples for the respective gene were defined as outliers and excluded for further analysis. Normal distribution was tested using the Lilliefors test implemented in MATLAB with a significance level of p<0.05. ΔCt values were correlated with metabolic parameters and other genes by Pearson correlation and corrected for age and gender by linear regression if applicable. Results prior adjustment are indicated with a p-value. All correlation results were corrected for multiple testing by controlling the FDR with the Benjamini-Hochberg procedure. A q-value<0.05 was considered as significant. Correlation matrices were plotted using the ggcorr-function of ggplot2-extension GGally (https://CRAN.R-project.org/package=GGally) and R 3.5.1 (The R Foundation for Statistical Computing, Vienna, Austria). Only applicable associations analyzed by Pearson's correlation are plotted. Non-tested associations are indicated by gray squares. Fold-change heatmaps and other graphs were generated by using GraphPad Prism 7.05 (GraphPad Software, Inc, San Diego, US).

**Target gene and pathway analysis**

Target genes of miRNAs were identified by database research from miRTarBase and TarBase [23,24]. Data tables from repositories of these databases were collected into a self-generated SQLite database and queried by a function we termed ‘miRNA Nvis’ implemented in MATLAB R2020a (The MathWorks, Natick, US). We implemented a target gene prediction tool as part of the microarray analysis framework to identify and list potential target genes of miRNAs. Thereby miRNA Nvis uses the mature miRNA and the respective 3’-UTR sequence of the target genes as a string and performs a seed match (2-7 nt of the 5’ miRNA sequence string) for different pairing modes (8mer, 7mer-m8, 7mer-A1, 6mer and offset 6mer)[25]. Moreover, this function computes favorable conditions for seed binding, such as the AU content within 60 nt surrounding the seed, additional binding 3’ to the seed sequence of the miRNA, the relative position of the seed within the 3’-UTR of the potential target, proximity to the stop codon and additional or cooperative binding of the same or other miRNA species [25,26]. The output can be exported as a table with all information on binding conditions and position for all pairing options within the sequence input string. Target genes were then manually selected by subsequent literature research and after stratification for T2D-related gene ontology terms.

For pathway enrichment analysis, we used only experimentally validated target genes from miRTarBase [23] as input for enrichKEGG from clusterProfiler (version 3.0.2 and R version 3.5) with the options organism = “hsa”, keyType = “kegg”, minGSize = 1 and otherwise default parameters.
