## Supplemental Tables 1-9 for "Liver microRNA transcriptome reveals miR-182 as link between type 2 diabetes and fatty liver disease in obesity"

**Supplementary Table 1: Clinical Characteristics for the complete (n=85) and microarray (n=40) obese human liver cohort. A ranksum test was performed to find significant differences between both subgroups.**

| | ND (n=44) all | | | ND (n=20) microarray | | | T2D (n=41) all | | | T2D (n=20) microarray | | | ranksum test complete cohort | ranksum test microarray |
|---|---|---|---|---|---|---|---|---|---|---|---|---|---|---|
| | Mean | SD | Count | | | | Mean | SD | Count | | | | | |
| **Diagnosed (yes/no)** | 0/44 | | 44 | 0/20 | | 20 | 36/5 | | | 20/0 | | 20 | NA | NA |
| **T2D by ADA-HbA$_{1c}$ (yes/no)** | 0/44 | | 44 | 0/20 | | 20 | 35/6 | | | 20/0 | | 20 | NA | NA |
| **HbA$_{1c}$ (%)** | 5.15 | 0.37 | 44 | 5.20 | 0.32 | 20 | 8.00 | 1.69 | 41 | 8.64 | 1.49 | 20 | *** | *** |
| **NAS (score)** | 1.90 | 1.95 | 41 | 2.50 | 2.25 | 20 | 3.72 | 1.99 | 32 | 4.15 | 1.53 | 20 | *** | * |
| **Sex** | m=7/w=37 | | 44 | m=7/w=13 | | 20 | m=14/w=27 | | 41 | m=8/w=12 | | 20 | n.s. | n.s. |
| **BMI (kg/m$^2$)** | 54.15 | 11.54 | 44 | 55.05 | 12.42 | 20 | 50.43 | 9.52 | 41 | 54.01 | 8.09 | 20 | n.s. | n.s. |
| **Age (years)** | 37.73 | 10.79 | 44 | 45.15 | 9.07 | 20 | 50.15 | 11.82 | 41 | 50.75 | 9.56 | 20 | *** | n.s. |
| **fasting glucose (mg/dL)** | 95.20 | 16.83 | 44 | 98.50 | 21.46 | 20 | 172.05 | 74.18 | 39 | 198.75 | 71.34 | 20 | *** | *** |
| **serum insulin (pmol/L)** | 74.29 | 51.60 | 42 | 90.07 | 55.90 | 19 | 169.09 | 195.79 | 36 | 140.61 | 125.27 | 16 | * | n.s. |
| **cholesterol (mg/dL)** | 185.77 | 32.55 | 43 | 191.15 | 29.80 | 20 | 191.16 | 49.68 | 38 | 186.89 | 56.65 | 19 | n.s. | n.s. |
| **serum triglycerides (mg/dL)** | 166.81 | 76.14 | 43 | 224.60 | 120.07 | 20 | 258.50 | 135.63 | 38 | 266.26 | 145.21 | 19 | *** | n.s. |
| **HDL (mg/dL)** | 46.63 | 11.93 | 43 | 42.60 | 10.69 | 20 | 42.68 | 13.51 | 38 | 40.79 | 13.66 | 19 | n.s. | n.s. |
| **LDL (mg/dL)** | 105.76 | 31.58 | 42 | 104.95 | 29.04 | 19 | 95.12 | 38.29 | 33 | 86.33 | 29.43 | 15 | n.s. | n.s. |
| **hepatic lipid content (%)** | 22.97 | 24.90 | 36 | 32.60 | 24.39 | 15 | 40.32 | 21.12 | 34 | 43.94 | 20.40 | 18 | *** | n.s. |
| **AST (U/L)** | 21.25 | 11.38 | 44 | 27.50 | 17.29 | 20 | 36.44 | 34.55 | 39 | 32.40 | 18.58 | 20 | ** | n.s. |
| **ALT (U/L)** | 27.75 | 18.97 | 44 | 36.35 | 27.58 | 20 | 37.36 | 18.97 | 39 | 34.95 | 12.85 | 20 | ** | n.s. |
| **CRP (mg/L)** | 14.39 | 11.13 | 44 | 13.75 | 10.03 | 20 | 16.68 | 14.81 | 38 | 19.25 | 16.96 | 20 | n.s. | n.s. |
| **TSH (mU/L)** | 2.21 | 1.22 | 40 | 1.93 | 0.80 | 19 | 2.79 | 7.31 | 37 | 3.96 | 9.63 | 20 | n.s. | n.s. |
| **Cortisol (nmol/L)** | 114.55 | 52.71 | 40 | 93.89 | 37.81 | 19 | 145.78 | 57.85 | 36 | 150.45 | 52.03 | 20 | * | *** |
| **Vitamin B12 (pmol/L)** | 509.83 | 320.50 | 40 | 538.11 | 368.75 | 19 | 593.92 | 319.40 | 36 | 533.26 | 195.12 | 19 | n.s. | n.s. |
| **folic acid (ng/mL)** | 5.98 | 3.88 | 40 | 6.95 | 5.10 | 19 | 8.74 | 5.39 | 36 | 8.43 | 5.10 | 19 | ** | n.s.+ |

Abbrevations: ADA, American Diabetes Association; ALT, Alanine transaminase; AST, Aspartate transaminase; BMI, Body mass index; CRP, C-reactive protein; HDL, high density lipoprotein; LDL, low density lipoprotein; m, men; NAS, NAFLD acitivity score; ND, non-diabetic; T2D, type 2 diabetic; TSH, thyroid stimulating hormone; w, women;

Group differences were tested by comparing ranks using the nonparametric Mann-Whitney test. Significane is indicated as $^{n.s}$p > 0.05; *p<0.05; **p<0.01; ***p<0.001. The microarray sub-cohort is part of the complete cohort.

**Supplementary Table 2: Primer sequences for SYBR-green analysis of cDNA by qPCR and cloning**

| Application | Gene | Species | Orientation | Sequence |
|---|---|---|---|---|
| mRNA quantification by qPCR | CASC3 | Human | forward | ACCTCGGAAAGGGCTCTTCTT |
| mRNA quantification by qPCR | CASC3 | Human | reverse | CGACCCTCATCCTTCCATAGC |
| mRNA quantification by qPCR | INSR | Human | forward | AAAACGAGGCCCGAAGATTTC |
| mRNA quantification by qPCR | INSR | Human | reverse | GAGCCCATAGACCCGGAAG |
| mRNA quantification by qPCR | Acaca | Mouse | forward | ATGGGCGGAATGGTCTCTTTC |
| mRNA quantification by qPCR | Acaca | Mouse | reverse | TGGGGACCTTGTCTTCATCAT |
| mRNA quantification by qPCR | Dhcr7 | Mouse | forward | AGGCTGGATCTCAAGGACAAT |
| mRNA quantification by qPCR | Dhcr7 | Mouse | reverse | GCCAGACTAGCATGGCCTG |
| mRNA quantification by qPCR | Hmgcs1 | Mouse | forward | AACTGGTGCAGAAATCTCTAGC |
| mRNA quantification by qPCR | Hmgcs1 | Mouse | reverse | GGTTGAATAGCTCAGAACTAGCC |
| mRNA quantification by qPCR | Insig1 | Mouse | forward | CACGACCACGTCTGGAACTAT |
| mRNA quantification by qPCR | Insig1 | Mouse | reverse | TGAGAAGAGCACTAGGCTCCG |
| mRNA quantification by qPCR | Elovl6 | Mouse | forward | GAAAAGCAGTTCAACGAGAACG |
| mRNA quantification by qPCR | Elovl6 | Mouse | reverse | AGATGCCGACCACCAAAGATA |
| mRNA quantification by qPCR | Fasn | Mouse | forward | GCCCCTTCAGTTGTGAGTCAT |
| mRNA quantification by qPCR | Fasn | Mouse | reverse | AGAGCGTTGCCGTATTCGTC |
| mRNA quantification by qPCR | Foxo1 | Mouse | forward | GAGGTGGTCGAGTTGGACTG |
| mRNA quantification by qPCR | Foxo1 | Mouse | reverse | CGCTGCAGATCCCGTAAGAC |
| mRNA quantification by qPCR | G6pc | Mouse | forward | TCTGTCCCGGATCTACCTTG |
| mRNA quantification by qPCR | G6pc | Mouse | reverse | GAAAGTTTCAGCCACAGCAA |
| mRNA quantification by qPCR | Insr | Mouse | forward | CGAGTGCCCGTCTGGCTATA |
| mRNA quantification by qPCR | Insr | Mouse | reverse | GGCAGGGTCCCAGACATG |
| mRNA quantification by qPCR | Irs1 | Mouse | forward | CGATGGCTTCTCAGACGTG |
| mRNA quantification by qPCR | Irs1 | Mouse | reverse | CAGCCCGCTTGTTGATGTTG |
| mRNA quantification by qPCR | Scd1 | Mouse | forward | TTCTTGCGATACACTCTGGTGC |
| mRNA quantification by qPCR | Scd1 | Mouse | reverse | CGGGATTGAATGTTCTTGTCGT |
| mRNA quantification by qPCR | Lrp6 | Mouse | forward | TTGTTGCTTTATGCAAACAGACG |
| mRNA quantification by qPCR | Lrp6 | Mouse | reverse | GTTCGTTTAATGGCTTCTTCGC |
| mRNA quantification by qPCR | Pck1 | Mouse | forward | CTGCATAACGGTCTGGACTTC |
| mRNA quantification by qPCR | Pck1 | Mouse | reverse | CAGCAACTGCCCGTACTCC |
| mRNA quantification by qPCR | Pdk1 | Mouse | forward | GGACTTCGGGTCAGTGAATGC |
| mRNA quantification by qPCR | Pdk1 | Mouse | reverse | TCCTGAGAAGATTGTCGGGGA |
| mRNA quantification by qPCR | Pnpla2 | Mouse | forward | GGATGGCGGCATTTCAGACA |
| mRNA quantification by qPCR | Pnpla2 | Mouse | reverse | CAAAGGGTTGGGTTGGTTCAG |
| mRNA quantification by qPCR | Ppara | Mouse | forward | TACTGCCGTTTTCACAAGTGC |
| mRNA quantification by qPCR | Ppara | Mouse | reverse | AGGTCGTGTTCACAGGTAAGA |
| mRNA quantification by qPCR | Hprt1 | Mouse | forward | AAGCTTGCTGGTGAAAAGGA |
| mRNA quantification by qPCR | Hprt1 | Mouse | reverse | TTGCGCTCATCTTAGGCTTT |
| miRNA quantification by qPCR | miR-182-5p | Human/mou | forward | GCTTTGGCAATGGTAGAACTCA |
| miRNA quantification by qPCR | miR-24-3p | Human/mou | forward | TGGCTCAGTTCAGCAGGAACA |
| miRNA quantification by qPCR | miR-149-5p | Mouse | forward | TCTGGCTCCGTGTCTTCACT |
| miRNA quantification by qPCR | miR-330-3p | Mouse | forward | CAAAGCACACGGCCT |
| miRNA quantification by qPCR | miR-1962 | Mouse | forward | GCTGGCACTGGGACA |
| miRNA quantification by qPCR | miR-7056-5p | Mouse | forward | TGTGGAGGAGGACAGAGA |
| miRNA quantification by qPCR | cel-miR-39-3p | | forward | CACCGGGTGTAAATCAGCTTG |
| miRNA quantification by qPCR | universal | | reverse | GCATAGACCTGAATGGCGGTA |
| cloning | LRP6 | human | forward | ATCTCTCGAGCACATGTGTAGCACTTCTCTGCT |
| cloning | LRP6 | human | reverse | TGCGGCCGCGACACTGTGGGCACTTAGTT |
| site-directed mutagenesis | LRP6 | human | sense | CTTTTTTAATTCTGTCTTGACAAGGCCTCTCTGGGTTTC |
| site-directed mutagenesis | LRP6 | human | antisense | GAAACCCAGAGAGGCCTTGTCAAGACAGAATTAAAAAAG |

**Supplementary Table 3: Assay-IDs for TaqManTM Gene-expression Assays**

| Application | Gene | Species | Assay-ID |
|---|---|---|---|
| mRNA quantification by qPCR | CASC3 | human | Hs00201226_m1 |
| mRNA quantification by qPCR | ACACA | human | Hs01046047_m1 |
| mRNA quantification by qPCR | G6PC | human | Hs00609178_m1 |
| mRNA quantification by qPCR | IRS1 | human | Hs00178563_m1 |
| mRNA quantification by qPCR | SCD | human | Hs01682761_m1 |
| mRNA quantification by qPCR | LRP6 | human | Hs00233945_m1 |
| mRNA quantification by qPCR | PCK1 | human | Hs00159918_m1 |
| mRNA quantification by qPCR | PDK1 | human | Hs01561847_m1 |
| mRNA quantification by qPCR | PNPLA2 | human | Hs00386101_m1 |
| mRNA quantification by qPCR | PPARA | human | Hs00947536_m1 |
| miRNA quantification by qPCR | miR-182-5p | human | 477935_mir |
| miRNA quantification by qPCR | miR-24-3p | | 477992_mir |

**Supplementary Table 4: Associations between miRNA and T2D. Logistic regression models for the incidence of T2D were generated using self-written scripts in MATLAB. Age, sex, BMI and the NAFLD activity score (NAS) were used as additional cofactors. The coefficient estimate and p value is indicated for the respective miRNA expression. ID is the respective identifier for the human microRNA on the GeneChipTM miRNA 4.0 Array.**

| NAS as cofactor | | | |
|---|---|---|---|
| **ID** | **TranscriptID** | **Coefficient_estimates** | **pvalue** |
| 20500149 | hsa-miR-24-2-5p | -1.126062525 | 0.022811482 |
| 20500187 | hsa-miR-29b-1-5p | -1.06213816 | 0.034091937 |
| 20500394 | hsa-miR-197-5p | -1.693000353 | 0.048694075 |
| 20500450 | hsa-miR-182-5p | -0.579180569 | 0.044961512 |
| 20500490 | hsa-miR-224-3p | -1.28840005 | 0.023203954 |
| 20500777 | hsa-miR-138-1-3p | -0.953986557 | 0.024387533 |
| 20500780 | hsa-miR-149-5p | -1.118781253 | 0.048797213 |
| 20501209 | hsa-miR-365a-5p | -1.513715732 | 0.042248794 |
| 20501249 | hsa-miR-381-3p | -1.889421194 | 0.014527909 |
| 20501276 | hsa-miR-330-3p | -1.586007318 | 0.00946818 |
| 20503103 | hsa-miR-485-5p | -1.2506043 | 0.035235638 |
| 20504572 | hsa-miR-1301-3p | -1.482193752 | 0.026019798 |
| 20506712 | hsa-miR-1180-3p | -1.156376275 | 0.046583398 |
| 20509071 | hsa-miR-1825 | 1.103671869 | 0.032006065 |
| 20511549 | hsa-miR-2110 | -1.051442863 | 0.039024194 |
| 20517680 | hsa-miR-4298 | -3.167436002 | 0.008247224 |
| 20517816 | hsa-miR-3609 | -0.878849249 | 0.015764196 |
| 20517824 | hsa-miR-3615 | -1.265248944 | 0.017494041 |
| 20518432 | hsa-miR-3911 | -1.447383511 | 0.043124403 |
| 20518879 | hsa-miR-4485 | 0.975733475 | 0.03773906 |
| 20518881 | hsa-miR-4487 | 1.13958907 | 0.047536744 |
| 20518935 | hsa-miR-4534 | -3.13143753 | 0.014752795 |
| 20519425 | hsa-miR-4646-5p | 1.184512509 | 0.010567489 |
| 20519467 | hsa-miR-4669 | -1.506362813 | 0.034425526 |
| 20519518 | hsa-miR-4701-3p | -1.958342956 | 0.024100997 |
| 20526885 | hsa-miR-7162-3p | -0.650144721 | 0.049172928 |
| 20529139 | hsa-miR-7847-3p | -2.918318612 | 0.015079767 |
| 20529781 | hsa-miR-8071 | -1.098997635 | 0.032240159 |

| NAS is not a cofactor | | | |
|---|---|---|---|
| **ID** | **TranscriptID** | **Coefficient_estimates** | **pvalue** |
| 20500121 | hsa-let-7e-5p | -1.240798343 | 0.026529746 |
| 20500149 | hsa-miR-24-2-5p | -1.087035336 | 0.011553292 |
| 20500187 | hsa-miR-29b-1-5p | -0.912339599 | 0.028875512 |
| 20500394 | hsa-miR-197-5p | -1.662124299 | 0.042097029 |
| 20500442 | hsa-miR-34a-5p | -1.141127971 | 0.040984788 |
| 20500450 | hsa-miR-182-5p | -0.634095195 | 0.017503866 |
| 20500489 | hsa-miR-224-5p | -0.792661281 | 0.016341158 |
| 20500490 | hsa-miR-224-3p | -1.436928039 | 0.005685515 |
| 20500777 | hsa-miR-138-1-3p | -0.923860686 | 0.024208758 |
| 20501083 | hsa-miR-155-5p | -1.148285797 | 0.029343942 |
| 20501177 | hsa-miR-99b-3p | -0.88765557 | 0.03919395 |
| 20501209 | hsa-miR-365a-5p | -1.380721568 | 0.036359842 |
| 20501249 | hsa-miR-381-3p | -1.3899115 | 0.018622208 |
| 20501276 | hsa-miR-330-3p | -1.405379753 | 0.007457949 |
| 20502235 | hsa-miR-18b-5p | -0.921105116 | 0.048244568 |
| 20503103 | hsa-miR-485-5p | -1.14942575 | 0.028583457 |
| 20503789 | hsa-miR-491-5p | -1.428675868 | 0.036225686 |
| 20503815 | hsa-miR-498 | -0.781359511 | 0.028689738 |
| 20506712 | hsa-miR-1180-3p | -1.368384999 | 0.021927127 |
| 20506779 | hsa-miR-1231 | -1.143148949 | 0.049455571 |
| 20509071 | hsa-miR-1825 | 1.082171943 | 0.019381526 |
| 20515610 | hsa-miR-3180-3p | -1.649820744 | 0.033453598 |
| 20517680 | hsa-miR-4298 | -2.896326194 | 0.009628718 |
| 20517710 | hsa-miR-4253 | -0.785381059 | 0.029027844 |
| 20517816 | hsa-miR-3609 | -0.835284615 | 0.012150248 |
| 20517824 | hsa-miR-3615 | -1.257207235 | 0.011402375 |
| 20517835 | hsa-miR-3621 | -1.707664864 | 0.046644899 |
| 20518432 | hsa-miR-3911 | -1.422234024 | 0.026315856 |
| 20518879 | hsa-miR-4485 | 1.064492457 | 0.017958399 |
| 20518935 | hsa-miR-4534 | -2.978432305 | 0.0146248 |
| 20519425 | hsa-miR-4646-5p | 1.04396474 | 0.01346794 |
| 20519467 | hsa-miR-4669 | -1.248255578 | 0.04614984 |
| 20519518 | hsa-miR-4701-3p | -1.725149372 | 0.024880843 |
| 20520208 | hsa-miR-5006-5p | -1.060709273 | 0.046867903 |
| 20525659 | hsa-miR-6849-5p | -1.09033502 | 0.039203136 |
| 20525739 | hsa-miR-6889-5p | -0.909322238 | 0.043090573 |
| 20526885 | hsa-miR-7162-3p | -0.693227695 | 0.027672155 |
| 20529139 | hsa-miR-7847-3p | -2.541620086 | 0.016964459 |
| 20529781 | hsa-miR-8071 | -1.031638736 | 0.027199168 |

**Supplementary Table 5: Associations between miRNAs and metabolic traits. Effect sizes describe the change of the trait if the miRNA expression changes by 1 log2 value. Age, sex, BMI and the NAFLD activity score (NAS) were used as additional cofactors whenever they did not serve as response value. ID is the respective identifier for the human microRNA on the GeneChipTM miRNA 4.0 Array.**

**Fasting glucose**

| ID | TranscriptID | Effect size (mg/dl) | pvalue |
|---|---|---|---|
| 20500181 | hsa-miR-99a-5p | -63.66280022 | 0.033300311 |
| 20500188 | hsa-miR-29b-3p | 24.9981997 | 0.021879139 |
| 20500450 | hsa-miR-182-5p | 15.05931669 | 0.023330445 |
| 20500469 | hsa-miR-212-3p | 21.96008843 | 0.029517028 |
| 20500555 | hsa-miR-200b-5p | 23.2295626 | 0.01569918 |
| 20500726 | hsa-miR-122-5p | -308.8007271 | 0.038362935 |
| 20500777 | hsa-miR-138-1-3p | 27.03728029 | 0.003919391 |
| 20501249 | hsa-miR-381-3p | 30.99292755 | 0.043971454 |
| 20501276 | hsa-miR-330-3p | 23.96377313 | 0.018866177 |
| 20503103 | hsa-miR-485-5p | 30.18963325 | 0.033486154 |
| 20503815 | hsa-miR-498 | 21.84842646 | 0.023343802 |
| 20504379 | hsa-miR-629-5p | 25.47803502 | 0.023456714 |
| 20506839 | hsa-miR-1247-3p | 17.61529435 | 0.019707088 |
| 20515603 | hsa-miR-3175 | -32.70444942 | 0.027489277 |
| 20515610 | hsa-miR-3180-3p | 45.60498957 | 0.015644691 |
| 20517680 | hsa-miR-4298 | 51.18981326 | 0.044496208 |
| 20517726 | hsa-miR-4269 | 23.4389842 | 0.033090519 |
| 20517820 | hsa-miR-3613-5p | -7.183487615 | 0.031272022 |
| 20518880 | hsa-miR-4486 | 45.53584312 | 0.033936611 |
| 20519494 | hsa-miR-1343-5p | 44.14511191 | 0.04807315 |
| 20519507 | hsa-miR-4695-5p | 49.5644105 | 0.023416318 |
| 20519607 | hsa-miR-4749-5p | 42.90451435 | 0.032967432 |
| 20521811 | hsa-miR-664b-3p | -37.57536784 | 0.001570536 |
| 20521825 | hsa-miR-5585-3p | 21.66527205 | 0.042655458 |
| 20525023 | hsa-miR-6511a-5p | 25.81401756 | 0.027725535 |
| 20525523 | hsa-miR-6781-5p | 24.61352015 | 0.048094489 |
| 20525539 | hsa-miR-6789-5p | 50.08317553 | 0.03084604 |
| 20525659 | hsa-miR-6849-5p | 32.86642473 | 0.01878475 |
| 20525739 | hsa-miR-6889-5p | 21.82819316 | 0.047945358 |
| 20526885 | hsa-miR-7162-3p | 17.18882928 | 0.042951976 |
| 20529781 | hsa-miR-8071 | 29.37695547 | 0.026504429 |

**HbA1c**

| ID | TranscriptID | Effect size (%) | pvalue |
|---|---|---|---|
| 20500149 | hsa-miR-24-2-5p | 0.660376999 | 0.02792399 |
| 20500150 | hsa-miR-25-5p | 0.760972884 | 0.01354268 |
| 20500181 | hsa-miR-99a-5p | -2.073332418 | 0.04955868 |
| 20500188 | hsa-miR-29b-3p | 0.611183244 | 0.0356238 |
| 20500394 | hsa-miR-197-5p | 1.371718291 | 0.01701529 |
| 20500450 | hsa-miR-182-5p | 0.472059468 | 0.04606791 |
| 20500490 | hsa-miR-224-3p | 0.806526173 | 0.01593033 |
| 20500726 | hsa-miR-122-5p | -10.05375408 | 0.00472102 |
| 20500777 | hsa-miR-138-1-3p | 0.748801853 | 0.0101476 |
| 20501177 | hsa-miR-99b-3p | 0.891550411 | 0.00428029 |
| 20501249 | hsa-miR-381-3p | 1.20045254 | 0.00089872 |
| 20501276 | hsa-miR-330-3p | 0.92445956 | 0.00039006 |
| 20503103 | hsa-miR-485-5p | 1.346853072 | 0.02278658 |
| 20503789 | hsa-miR-491-5p | 1.11452125 | 0.0098573 |
| 20504379 | hsa-miR-629-5p | 0.818528306 | 0.0311345 |
| 20504569 | hsa-miR-1271-5p | 0.975571366 | 0.02046567 |
| 20504572 | hsa-miR-1301-3p | 0.972796561 | 0.00641999 |
| 20505791 | hsa-miR-877-5p | 1.804065946 | 0.04329844 |
| 20506712 | hsa-miR-1180-3p | 0.777300576 | 0.00541548 |
| 20517680 | hsa-miR-4298 | 1.942646889 | 0.01683822 |
| 20517816 | hsa-miR-3609 | 0.557378801 | 0.01313464 |
| 20519467 | hsa-miR-4669 | 1.119340924 | 0.03207667 |
| 20521811 | hsa-miR-664b-3p | -0.754147594 | 0.02301778 |
| 20525659 | hsa-miR-6849-5p | 0.901536062 | 0.01292098 |
| 20529139 | hsa-miR-7847-3p | 1.87641212 | |

**Serum insulin**

| ID | TranscriptID | Effect size (pmol/l) | pvalue |
|---|---|---|---|
| 20500115 | hsa-let-7b-5p | -114.7110822 | 0.0319743 |
| 20500164 | hsa-miR-31-5p | -19.46165022 | 0.047548 |
| 20500173 | hsa-miR-93-5p | -84.73214297 | 0.0177801 |
| 20500489 | hsa-miR-224-5p | 29.93448592 | 0.0180121 |
| 20500745 | hsa-miR-140-5p | 26.4891447 | 0.0485933 |
| 20500746 | hsa-miR-140-3p | -93.35904586 | 0.0070981 |
| 20500787 | hsa-miR-185-5p | -79.22898225 | 0.0251669 |
| 20500848 | hsa-miR-320a | -71.27490064 | 0.0095674 |
| 20501243 | hsa-miR-378a-3p | -66.72678983 | 0.0382696 |
| 20501291 | hsa-miR-148b-3p | 28.72004889 | 0.0227703 |
| 20502235 | hsa-miR-18b-5p | 44.57166127 | 0.0206499 |
| 20502451 | hsa-miR-452-5p | 36.73690552 | 0.0154262 |
| 20503105 | hsa-miR-486-5p | -33.84092605 | 0.011823 |
| 20504295 | hsa-miR-572 | 47.11064396 | 0.0065453 |
| 20504421 | hsa-miR-654-3p | 33.7023895 | 0.0338244 |
| 20504562 | hsa-miR-320b | -70.2959101 | 0.0123537 |
| 20504563 | hsa-miR-320c | -75.17117163 | 0.0124384 |
| 20505790 | hsa-miR-885-3p | 79.82301748 | 0.0258005 |
| 20506872 | hsa-miR-1275 | -74.18177938 | 0.0156031 |
| 20509070 | hsa-miR-320d | -61.91855055 | 0.0406623 |
| 20509228 | hsa-miR-1910-5p | 44.36571618 | 0.0158373 |
| 20509237 | hsa-miR-1915-3p | 79.37997518 | 0.0181387 |
| 20511558 | hsa-miR-2115-5p | -59.33539845 | 0.0235906 |
| 20515603 | hsa-miR-3175 | -52.93820784 | 0.0247474 |
| 20515617 | hsa-miR-3185 | 80.66435819 | 0.0048253 |
| 20517690 | hsa-miR-4310 | 57.86025408 | 0.0107995 |
| 20517730 | hsa-miR-4271 | 28.56066784 | 0.0471921 |
| 20517835 | hsa-miR-3621 | 68.11684831 | 0.0387131 |
| 20517899 | hsa-miR-3648 | 35.40769076 | 0.0439624 |
| 20517919 | hsa-miR-3665 | 109.678474 | 0.0078236 |
| 20518801 | hsa-miR-4429 | -77.34139562 | 0.0319667 |
| 20518834 | hsa-miR-4454 | -38.48901508 | 0.0147287 |
| 20518843 | hsa-miR-3135b | 39.77284466 | 0.0312321 |
| 20518852 | hsa-miR-4467 | 46.77244449 | 0.0169314 |
| 20518910 | hsa-miR-4513 | 31.69991427 | 0.0476149 |
| 20518933 | hsa-miR-4532 | 51.52360291 | 0.0389452 |
| 20519048 | hsa-miR-3960 | 98.00987056 | 0.0158992 |
| 20519463 | hsa-miR-4668-5p | -34.59339979 | 0.0490228 |
| 20519474 | hsa-miR-4674 | 68.45539447 | 0.0351662 |
| 20519507 | hsa-miR-4695-5p | 70.89728482 | 0.0262261 |
| 20519580 | hsa-miR-4734 | 68.02239165 | 0.0488992 |
| 20519589 | hsa-miR-4739 | 61.37649256 | 0.0453221 |
| 20519615 | hsa-miR-371b-5p | 21.30757495 | 0.0288355 |
| 20519679 | hsa-miR-4787-5p | 119.2927514 | 0.0053233 |
| 20519702 | hsa-miR-4800-3p | 28.83403774 | 0.0364043 |
| 20520198 | hsa-miR-5001-5p | 53.35771351 | 0.0106614 |
| 20521825 | hsa-miR-5585-3p | 38.09071841 | 0.0123428 |
| 20523000 | hsa-miR-6068 | 49.74774142 | 0.0074984 |
| 20523007 | hsa-miR-6075 | 50.9348939 | 0.0008842 |
| 20524035 | hsa-miR-6125 | 79.63387901 | 0.0212661 |
| 20525017 | hsa-miR-6508-5p | 41.30704059 | 0.0175202 |
| 20525499 | hsa-miR-6769a-5p | 73.6996385 | 0.0045716 |
| 20525509 | hsa-miR-6774-5p | 31.32261652 | 0.0380932 |
| 20525539 | hsa-miR-6789-5p | 71.91805233 | 0.0293751 |
| 20525659 | hsa-miR-6849-5p | 62.38971507 | 0.0022223 |
| 20525699 | hsa-miR-6869-5p | 82.05955664 | 0.0267813 |
| 20526178 | hsa-miR-7110-5p | -65.30418345 | 0.0318021 |
| 20526885 | hsa-miR-7162-3p | 30.86799199 | 0.0181154 |
| 20528733 | hsa-miR-7704 | 91.68401363 | 0.0376099 |
| 20529566 | hsa-miR-7975 | -24.21904311 | 0.031434 |
| 20529568 | hsa-miR-7977 | -32.81241496 | 0.0471134 |
| 20529774 | hsa-miR-8064 | 29.8649197 | 0.0418896 |
| 20529783 | hsa-miR-8073 | 26.02839473 | 0.0482356 |
| 20529785 | hsa-miR-8075 | -73.69114191 | 0.0117931 |

**Serum triglycerides**

| ID | TranscriptID | Effect size (mg/dl) | pvalue |
|---|---|---|---|
| 20500142 | hsa-miR-21-3p | 50.08425257 | 0.0486782 |
| 20500159 | hsa-miR-28-3p | -102.7234757 | 0.0422017 |
| 20500164 | hsa-miR-31-5p | 35.37525142 | 0.0054054 |
| 20500395 | hsa-miR-197-3p | -71.06256065 | 0.0054828 |
| 20500448 | hsa-miR-181c-5p | 64.42576109 | 0.0391396 |
| 20500461 | hsa-miR-204-3p | 62.02282544 | 0.0066963 |
| 20500724 | hsa-miR-30b-5p | -49.77523881 | 0.0436168 |
| 20500741 | hsa-miR-135a-3p | 62.68077602 | 0.0044663 |
| 20500780 | hsa-miR-149-5p | -53.88220808 | 0.0424714 |
| 20500781 | hsa-miR-149-3p | 100.9673616 | 0.0153507 |
| 20503803 | hsa-miR-494-3p | -112.7202908 | 0.0442979 |
| 20503887 | hsa-miR-505-3p | -46.80394921 | 0.0080958 |
| 20503908 | hsa-miR-532-3p | -63.93146803 | 0.0493159 |
| 20504273 | hsa-miR-92b-5p | 102.4227359 | 0.0046645 |
| 20504298 | hsa-miR-574-3p | -60.24957287 | 0.0438155 |
| 20504324 | hsa-miR-550a-5p | 65.20456983 | 0.0156667 |
| 20504396 | hsa-miR-642a-3p | 39.3450694 | 0.0009644 |
| 20504413 | hsa-miR-663a | 84.35571409 | 0.0378762 |
| 20504559 | hsa-miR-1224-5p | 51.0305232 | 0.0133812 |
| 20505789 | hsa-miR-885-5p | -77.47517523 | 0.0347254 |
| 20506009 | hsa-miR-939-5p | 71.09546605 | 0.0133274 |
| 20506801 | hsa-miR-1207-5p | 95.39245798 | 0.0026669 |
| 20507742 | hsa-miR-1469 | 124.6511768 | 0.0306403 |
| 20515623 | hsa-miR-3187-3p | 46.78956561 | 0.0303583 |
| 20517710 | hsa-miR-4253 | 49.19274843 | 0.0025061 |
| 20517745 | hsa-miR-4286 | -28.26780013 | 0.0241786 |
| 20517907 | hsa-miR-3656 | 96.64845845 | 0.0392785 |
| 20518721 | hsa-miR-642b-3p | 40.01140349 | 0.0029987 |
| 20518782 | hsa-miR-1268b | 100.0182516 | 0.0147945 |
| 20518800 | hsa-miR-4428 | 66.25495032 | 0.0064779 |
| 20518802 | hsa-miR-4430 | 77.03307295 | 0.0352248 |
| 20518807 | hsa-miR-4433-3p | 102.1849835 | 0.0086011 |
| 20518818 | hsa-miR-4443 | 79.85100991 | 0.0089457 |
| 20518826 | hsa-miR-4449 | 69.49968801 | 0.0219871 |
| 20518839 | hsa-miR-4459 | 123.0269313 | 0.0196477 |
| 20518845 | hsa-miR-4463 | 104.9801422 | 0.0238506 |
| 20518878 | hsa-miR-4484 | 35.31107907 | 0.0296754 |
| 20518881 | hsa-miR-4487 | 63.99040424 | 0.0197624 |
| 20518892 | hsa-miR-4497 | 175.4639649 | 0.001058 |
| 20518935 | hsa-miR-4534 | 100.0671355 | 0.0237637 |
| 20519457 | hsa-miR-4665-5p | 87.19298512 | 0.0098648 |
| 20519497 | hsa-miR-4689 | 81.77009816 | 0.0271815 |
| 20519524 | hsa-miR-4706 | 68.68848811 | 0.0253045 |
| 20519554 | hsa-miR-4721 | 42.6603946 | 0.0382923 |
| 20519600 | hsa-miR-4745-5p | 97.18696644 | 0.0231352 |
| 20519626 | hsa-miR-4758-5p | 135.5578843 | 0.0011152 |
| 20520218 | hsa-miR-5010-5p | 65.12393173 | 0.0092994 |
| 20520577 | hsa-miR-5196-5p | 58.213598 | 0.0220347 |
| 20521786 | hsa-miR-5572 | 55.01512566 | 0.0132016 |
| 20521811 | hsa-miR-664b-3p | -49.53116747 | 0.0450987 |
| 20521837 | hsa-miR-5589-3p | -45.07119025 | 0.0423704 |
| 20522537 | hsa-miR-5787 | 111.3057854 | 0.0168262 |
| 20523018 | hsa-miR-6086 | 55.28662251 | 0.0433946 |
| 20524036 | hsa-miR-6126 | 61.93524725 | 0.0058541 |
| 20525383 | hsa-miR-6716-5p | 109.9120439 | 0.0002835 |
| 20525444 | hsa-miR-6741-5p | 66.00849207 | 0.0318247 |
| 20525465 | hsa-miR-6752-5p | 92.81555392 | 0.0141595 |
| 20525469 | hsa-miR-6754-5p | 30.29499244 | 0.0315119 |
| 20525487 | hsa-miR-6763-5p | 98.28387949 | 0.0080386 |
| 20525491 | hsa-miR-6765-5p | 125.8560685 | 0.0086862 |
| 20525497 | hsa-miR-6768-5p | 44.15913796 | 0.0282417 |
| 20525513 | hsa-miR-6776-5p | 67.20954428 | 0.0136579 |
| 20525525 | hsa-miR-6782-5p | 87.34829056 | 0.0013683 |
| 20525533 | hsa-miR-6786-5p | 130.6119552 | 0.0157769 |
| 20525535 | hsa-miR-6787-5p | 75.31037003 | 0.0464252 |
| 20525541 | hsa-miR-6790-5p | 82.82007963 | 0.0183736 |
| 20525549 | hsa-miR-6794-5p | 97.38551698 | 0.0117046 |
| 20525557 | hsa-miR-6798-5p | 88.78104074 | 0.0076302 |
| 20525565 | hsa-miR-6802-5p | 97.46898482 | 0.0178135 |
| 20525567 | hsa-miR-6803-5p | 102.9401739 | 0.0251537 |
| 20525577 | hsa-miR-6808-5p | 112.1224589 | 0.0167276 |
| 20525609 | hsa-miR-6824-5p | 62.92983416 | 0.0322147 |
| 20525611 | hsa-miR-6825-5p | 62.99395585 | 0.0250129 |
| 20525613 | hsa-miR-6826-5p | 73.48036984 | 0.0037254 |
| 20525633 | hsa-miR-6780b-5p | 42.99661863 | 0.0010749 |
| 20525644 | hsa-miR-6840-3p | 39.72376532 | 0.025044 |
| 20525683 | hsa-miR-6860 | 62.18296293 | 0.0365236 |
| 20525701 | hsa-miR-6870-5p | 82.20413946 | 0.0341289 |
| 20525703 | hsa-miR-6871-5p | 57.689363 | 0.024121 |
| 20525719 | hsa-miR-6879-5p | 98.98636811 | 0.015095 |
| 20525743 | hsa-miR-6891-5p | 81.13009844 | 0.0374277 |
| 20526170 | hsa-miR-7106-5p | 63.28872538 | 0.0381292 |
| 20526172 | hsa-miR-7107-5p | 117.1620187 | 0.0131619 |
| 20529131 | hsa-miR-4433b-3p | 97.27291087 | 0.0146591 |
| 20529143 | hsa-miR-7851-3p | 29.16852909 | 0.0467373 |

**NAS**

| ID | TranscriptID | Effect size (score) | pvalue |
|---|---|---|---|
| 20500164 | hsa-miR-31-5p | 0.297417961 | 0.032643 |
| 20500400 | hsa-miR-199a-3p | 0.151546626 | 0.0116789 |
| 20500433 | hsa-miR-139-3p | -0.137996761 | 0.0393452 |
| 20500442 | hsa-miR-34a-5p | 0.127212674 | 0.0270248 |
| 20500450 | hsa-miR-182-5p | 0.281202284 | 0.0395469 |
| 20500458 | hsa-miR-199b-3p | 0.151546626 | 0.0116789 |
| 20500489 | hsa-miR-224-5p | 0.454668445 | 2.75E-06 |
| 20500490 | hsa-miR-224-3p | 0.259411688 | 0.0003291 |
| 20500556 | hsa-miR-200b-3p | 0.307112536 | 0.0083946 |
| 20500715 | hsa-let-7i-5p | 0.089732514 | 0.0470841 |
| 20500746 | hsa-miR-140-3p | -0.076198569 | 0.0377642 |
| 20500796 | hsa-miR-193a-3p | 0.155323585 | 0.0399858 |
| 20501083 | hsa-miR-155-5p | 0.153958847 | 0.0209109 |
| 20501163 | hsa-miR-200a-3p | 0.315031006 | 0.0054494 |
| 20501224 | hsa-miR-376c-3p | 0.199539608 | 0.0153768 |
| 20501242 | hsa-miR-378a-5p | -0.105824082 | 0.0235599 |
| 20501243 | hsa-miR-378a-3p | -0.107782849 | 0.0086162 |
| 20501291 | hsa-miR-148b-3p | 0.213699362 | 0.0464779 |
| 20502451 | hsa-miR-452-5p | 0.353791142 | 2.68E-05 |
| 20503100 | hsa-miR-483-5p | -0.127386792 | 0.0172393 |
| 20503793 | hsa-miR-146b-5p | 0.144252007 | 0.0242275 |
| 20503811 | hsa-miR-181d-5p | 0.129701901 | 0.0391071 |
| 20504273 | hsa-miR-92b-5p | 0.119937756 | 0.0127754 |
| 20504396 | hsa-miR-642a-3p | 0.388062334 | 0.0062842 |
| 20506882 | hsa-miR-1281 | -0.108515394 | 0.0404831 |
| 20515610 | hsa-miR-3180-3p | 0.135947234 | 0.0033892 |
| 20517675 | hsa-miR-378c | -0.084218729 | 0.0479962 |
| 20517710 | hsa-miR-4253 | 0.268915591 | 0.0113427 |
| 20518425 | hsa-miR-3180 | 0.118308935 | 0.0203599 |
| 20518446 | hsa-miR-3921 | -0.15564759 | 0.0341761 |
| 20518802 | hsa-miR-4430 | 0.106582863 | 0.0309841 |
| 20518818 | hsa-miR-4443 | 0.14509551 | 0.0118879 |
| 20518874 | hsa-miR-4481 | 0.1994447 | 0.004912 |
| 20518936 | hsa-miR-378i | -0.084067802 | 0.0324355 |
| 20519439 | hsa-miR-4655-5p | 0.164609493 | 0.0444508 |
| 20519498 | hsa-miR-4690-5p | 0.10350598 | 0.0171747 |
| 20519524 | hsa-miR-4706 | 0.138102181 | 0.0178876 |
| 20519533 | hsa-miR-4710 | 0.145991707 | 0.0029219 |
| 20519554 | hsa-miR-4721 | 0.200792785 | 0.0219112 |
| 20519609 | hsa-miR-4750-5p | 0.122253771 | 0.0075841 |
| 20521825 | hsa-miR-5585-3p | 0.177104255 | 0.0382307 |
| 20523018 | hsa-miR-6086 | 0.133293189 | 0.0461745 |
| 20525386 | hsa-miR-6511b-5p | 0.146223595 | 0.0346745 |
| 20525453 | hsa-miR-6746-5p | 0.177557819 | 0.0406788 |
| 20525513 | hsa-miR-6776-5p | 0.146935162 | 0.0246324 |
| 20525519 | hsa-miR-6779-5p | 0.101175548 | 0.0081308 |
| 20525525 | hsa-miR-6782-5p | 0.136875203 | 0.0312569 |
| 20525587 | hsa-miR-6813-5p | 0.290600765 | 0.010268 |
| 20525653 | hsa-miR-6846-5p | 0.141312052 | 0.0280904 |
| 20525657 | hsa-miR-6848-5p | 0.116074984 | 0.0163035 |
| 20525684 | hsa-miR-6861-5p | 0.167382674 | 0.0273753 |
| 20525735 | hsa-miR-6887-5p | 0.152226076 | 0.030194 |

**Supplementary Table 6: Associations between miRNAs and excluding factors age, BMI and sex. To control for associations with both confounding factors, also the HbA1c level was considered as cofactor for linear regression models. Effect sizes describe the change of the trait if the miRNA expression changes by 1 log2 value. ID is the respective identifier for the human microRNA on the GeneChipTM miRNA 4.0 Array.**

**Age**

| ID | TranscriptID | Effect size (years) | pvalue |
|---|---|---|---|
| 20500112 | hsa-let-7a-5p | -4.419927305 | 0.042853537 |
| 20500119 | hsa-let-7d-5p | -5.839419589 | 0.047445394 |
| 20500121 | hsa-let-7e-5p | -4.395563611 | 0.029236182 |
| 20500123 | hsa-let-7f-5p | -3.224283323 | 0.047797105 |
| 20500154 | hsa-miR-26b-5p | -2.372769966 | 0.037892631 |
| 20500422 | hsa-miR-30c-5p | -4.796226969 | 0.04711037 |
| 20500432 | hsa-miR-139-5p | -4.69845816 | 0.012706722 |
| 20500489 | hsa-miR-224-5p | -3.124774331 | 0.044213328 |
| 20500721 | hsa-miR-23b-3p | -6.115126205 | 0.039985152 |
| 20500724 | hsa-miR-30b-5p | -3.399520917 | 0.037727989 |
| 20500769 | hsa-miR-126-3p | -6.450119915 | 0.020496411 |
| 20500797 | hsa-miR-194-5p | -5.992987295 | 0.030056113 |
| 20501183 | hsa-miR-30e-3p | -2.734264856 | 0.020947379 |
| 20501224 | hsa-miR-376c-3p | -3.524504115 | 0.022924422 |
| 20501244 | hsa-miR-379-5p | -6.347263188 | 0.021755107 |
| 20501281 | hsa-miR-337-5p | -3.493573361 | 0.013571476 |
| 20502235 | hsa-miR-18b-5p | -3.399321847 | 0.035000237 |
| 20503102 | hsa-miR-484 | -4.753021128 | 0.006262291 |
| 20503103 | hsa-miR-485-5p | -4.634440168 | 0.043670022 |
| 20503800 | hsa-miR-432-5p | -5.894436508 | 0.019875154 |
| 20503877 | hsa-miR-501-5p | -3.817246908 | 0.035865966 |
| 20504218 | hsa-miR-487b-3p | -4.130125394 | 0.015000433 |
| 20504379 | hsa-miR-629-5p | -3.674826474 | 0.026071477 |
| 20504421 | hsa-miR-654-3p | -2.974744815 | 0.029642496 |
| 20506776 | hsa-miR-1229-5p | 5.214526408 | 0.028296834 |
| 20515585 | hsa-miR-3162-5p | 6.359800477 | 0.017617996 |
| 20517680 | hsa-miR-4298 | -8.054019527 | 0.031892115 |
| 20517816 | hsa-miR-3609 | -2.731179233 | 0.020406247 |
| 20517948 | hsa-miR-3687 | 2.025472155 | 0.049913995 |
| 20518818 | hsa-miR-4443 | -4.537756239 | 0.047943941 |
| 20518879 | hsa-miR-4485 | 3.075414227 | 0.028702676 |
| 20518892 | hsa-miR-4497 | -8.027125948 | 0.033976 |
| 20519493 | hsa-miR-4687-3p | 5.916680398 | 0.035885308 |
| 20519636 | hsa-miR-4763-3p | 5.878863077 | 0.046012972 |
| 20520349 | hsa-miR-1273f | 2.487760878 | 0.045648661 |
| 20525603 | hsa-miR-6821-5p | 6.267358231 | 0.048445422 |
| 20525699 | hsa-miR-6869-5p | 7.265899867 | 0.038928874 |
| 20529132 | hsa-miR-1273h-5p | 2.656903267 | 0.019455407 |

**BMI**

| ID | TranscriptID | Effect size (kg/m^2) | pvalue |
|---|---|---|---|
| 20500144 | hsa-miR-22-3p | -7.098861175 | 0.02871589 |
| 20500158 | hsa-miR-28-5p | -7.799023622 | 0.04780724 |
| 20500163 | hsa-miR-30a-3p | -4.668150377 | 0.04682778 |
| 20500164 | hsa-miR-31-5p | -1.99056271 | 0.04919936 |
| 20500181 | hsa-miR-99a-5p | -11.36031667 | 0.01257518 |
| 20500183 | hsa-miR-100-5p | -8.399122048 | 0.01359067 |
| 20500386 | hsa-miR-192-3p | -5.660312592 | 0.02918485 |
| 20500424 | hsa-miR-30d-5p | -8.304935608 | 0.03933463 |
| 20500471 | hsa-miR-214-5p | -4.523696876 | 0.02350643 |
| 20500722 | hsa-miR-27b-5p | -4.965718362 | 0.031749 |
| 20500781 | hsa-miR-149-3p | 6.683750947 | 0.02691111 |
| 20501179 | hsa-miR-296-3p | 7.257369069 | 0.00214253 |
| 20501242 | hsa-miR-378a-5p | -6.855182083 | 0.01994397 |
| 20501244 | hsa-miR-379-5p | -7.095477881 | 0.0169212 |
| 20501249 | hsa-miR-381-3p | -4.992215199 | 0.04649563 |
| 20501277 | hsa-miR-328-5p | 9.194410752 | 0.0067222 |
| 20501293 | hsa-miR-331-3p | -3.625222835 | 0.02568889 |
| 20502122 | hsa-miR-422a | -5.904585961 | 0.03517289 |
| 20502126 | hsa-miR-424-3p | -6.01599017 | 0.02343512 |
| 20503803 | hsa-miR-494-3p | -8.269607419 | 0.04810279 |
| 20503815 | hsa-miR-498 | 3.016591203 | 0.04814493 |
| 20504218 | hsa-miR-487b-3p | -3.937362295 | 0.03937509 |
| 20504273 | hsa-miR-92b-5p | 6.457268874 | 0.02833517 |
| 20504391 | hsa-miR-638 | 7.540352493 | 0.04148059 |
| 20504559 | hsa-miR-1224-5p | 3.259334697 | 0.0418891 |
| 20504586 | hsa-miR-762 | 8.182195561 | 0.0439578 |
| 20506763 | hsa-miR-1225-5p | 4.736207761 | 0.03356762 |
| 20506774 | hsa-miR-1228-5p | 7.431769601 | 0.03427703 |
| 20506801 | hsa-miR-1207-5p | 4.770415074 | 0.04544078 |
| 20515550 | hsa-miR-3141 | 7.468204297 | 0.02109152 |
| 20515618 | hsa-miR-3065-5p | -4.518867578 | 0.03227897 |
| 20517690 | hsa-miR-4310 | 5.106110078 | 0.01223967 |
| 20517717 | hsa-miR-4327 | 3.34219455 | 0.04210695 |
| 20517729 | hsa-miR-4270 | 6.860508696 | 0.02896455 |
| 20517736 | hsa-miR-4281 | 9.095434747 | 0.00250261 |
| 20517907 | hsa-miR-3656 | 7.189565818 | 0.0336131 |
| 20518783 | hsa-miR-378e | -3.203904854 | 0.03883934 |
| 20518788 | hsa-miR-378f | -6.409064755 | 0.04089703 |
| 20518794 | hsa-miR-378g | -5.841165377 | 0.02575099 |
| 20518800 | hsa-miR-4428 | 3.832504221 | 0.04639508 |
| 20518839 | hsa-miR-4459 | 8.813108333 | 0.02144266 |
| 20518851 | hsa-miR-4466 | 8.712015587 | 0.04503238 |
| 20518903 | hsa-miR-2392 | 6.555565539 | 0.02498024 |
| 20518931 | hsa-miR-4530 | 5.622075338 | 0.03582399 |
| 20518936 | hsa-miR-378i | -8.464619147 | 0.01434749 |
| 20518940 | hsa-miR-1587 | 5.084992471 | 0.02152021 |
| 20519429 | hsa-miR-4649-5p | 5.588071847 | 0.03490998 |
| 20519493 | hsa-miR-4687-3p | 6.091192189 | 0.04900076 |
| 20519494 | hsa-miR-1343-5p | 7.742546083 | 0.01930758 |
| 20519525 | hsa-miR-4707-5p | 7.73057443 | 0.03194012 |
| 20519607 | hsa-miR-4749-5p | 6.740222531 | 0.03015396 |
| 20519626 | hsa-miR-4758-5p | 6.490790847 | 0.04575511 |
| 20520569 | hsa-miR-5189-3p | -4.508340232 | 0.01053344 |
| 20521811 | hsa-miR-664b-3p | -3.92738696 | 0.04216495 |
| 20521825 | hsa-miR-5585-3p | 3.71194448 | 0.02373439 |
| 20522537 | hsa-miR-5787 | 7.969605813 | 0.01905971 |
| 20523017 | hsa-miR-6085 | 8.334485248 | 0.0192028 |
| 20523019 | hsa-miR-6087 | 15.79070915 | 0.00674514 |
| 20523020 | hsa-miR-6088 | 9.370619428 | 0.01476177 |
| 20524034 | hsa-miR-6124 | 4.976324972 | 0.04381062 |
| 20524054 | hsa-miR-6133 | 5.511313859 | 0.03342694 |
| 20525392 | hsa-miR-6721-5p | 2.758554243 | 0.03415762 |
| 20525394 | hsa-miR-6722-3p | 6.505013899 | 0.03523324 |
| 20525396 | hsa-miR-6724-5p | 5.777659792 | 0.04352059 |
| 20525420 | hsa-miR-6729-5p | 8.497427863 | 0.03894006 |
| 20525448 | hsa-miR-6743-5p | 5.42924222 | 0.0394956 |
| 20525459 | hsa-miR-6749-5p | 8.132866982 | 0.01419978 |
| 20525511 | hsa-miR-6775-5p | 13.57572826 | 0.0016103 |
| 20525519 | hsa-miR-6779-5p | 8.927920075 | 0.01361481 |
| 20525531 | hsa-miR-6785-5p | 5.665136657 | 0.01405824 |
| 20525533 | hsa-miR-6786-5p | 8.426933202 | 0.03473565 |
| 20525565 | hsa-miR-6802-5p | 6.645209918 | 0.02711902 |
| 20525577 | hsa-miR-6808-5p | 8.749422937 | 0.00704848 |
| 20525578 | hsa-miR-6808-3p | -5.091323641 | 0.02176278 |
| 20525585 | hsa-miR-6812-5p | 11.67978928 | 0.01585062 |
| 20525657 | hsa-miR-6848-5p | 6.243085764 | 0.03171274 |
| 20525681 | hsa-miR-6769b-5p | 10.14371175 | 0.00897609 |
| 20525684 | hsa-miR-6861-5p | 4.395229209 | 0.01313822 |
| 20525743 | hsa-miR-6891-5p | 7.126018348 | 0.00700704 |
| 20526172 | hsa-miR-7107-5p | 8.10353794 | 0.01864312 |
| 20526861 | hsa-miR-7150 | 4.858392956 | 0.02783996 |
| 20529131 | hsa-miR-4433b-3p | 7.146553412 | 0.01201086 |
| 20529137 | hsa-miR-7845-5p | 7.744673952 | 0.001808 |
| 20529799 | hsa-miR-8089 | 5.134438594 | 0.02512747 |

**Sex**

| ID | TranscriptID | Coefficient estimate | pvalue |
|---|---|---|---|
| 20500489 | hsa-miR-224-5p | -0.985218993 | 0.02305482 |
| 20500490 | hsa-miR-224-3p | -1.271340976 | 0.02386457 |
| 20500762 | hsa-miR-191-3p | 1.220163099 | 0.04072433 |
| 20500798 | hsa-miR-195-5p | -1.608400428 | 0.04332735 |
| 20502451 | hsa-miR-452-5p | -1.510417859 | 0.01877042 |
| 20503809 | hsa-miR-497-5p | -2.333799238 | 0.03842024 |
| 20504289 | hsa-miR-551b-5p | -1.522616403 | 0.01685506 |
| 20505760 | hsa-miR-708-5p | -0.680802107 | 0.04375668 |
| 20506712 | hsa-miR-1180-3p | -1.134140892 | 0.04673579 |
| 20506830 | hsa-miR-1304-3p | 1.244115413 | 0.01943593 |
| 20518785 | hsa-miR-4417 | 0.847512595 | 0.03474595 |
| 20518933 | hsa-miR-4532 | 1.437533625 | 0.03728473 |
| 20525017 | hsa-miR-6508-5p | 0.986543308 | 0.03696451 |

**Supplementary Table 7: Predicted target genes from miRTarBase used for pathway analysis.**

| pathway_list | hsa-miR-138-1-3p | hsa-miR-149-5p | hsa-miR-182-5p | hsa-miR-224-3p | hsa-miR-24-2-5p | hsa-miR-330-3p | hsa-miR-381-3p | hsa-miR-485-5p |
|---|---|---|---|---|---|---|---|---|
| HIF-1 signaling pathway | PDK1;HIF1A | IL6 | CDKN1A;BCL2;CDKN1B | | BCL2 | VEGFA | | |
| Thyroid hormone signaling pathway | HIF1A | | FOXO1;GSK3B | | | | | |
| AGE-RAGE signaling pathway in diabetic complications | | IL6 | FOXO1;BCL2;SMAD4;CDKN1B | | BCL2 | VEGFA;CDC42 | | |
| Apoptosis | | BBC3;FASLG | BCL2 | | BCL2 | | NFKBIA | |
| cAMP signaling pathway | | PTGER2 | ADCY6;CREB1;BDNF;CREB5;PLD1;TIAM1 | | | | NFKBIA | |
| Cellular senescence | | FOXM1;IL6 | CDKN1A;FOXO3;FOXO1;CCND2;PTEN;CHEK2 | | | E2F1 | | |
| Cytokine-cytokine receptor interaction | | IL6;FASLG | | | | | | |
| Cytosolic DNA-sensing pathway | | IL6 | | | | | NFKBIA | |
| EGFR tyrosine kinase inhibitor resistance | | IL6 | FOXO3;BCL2;PTEN;GSK3B | | BCL2 | VEGFA | | |
| Endocrine resistance | | SP1 | CDKN1A;ADCY6;BCL2;CDKN1B | | BCL2 | E2F1;SP1 | | |
| Estrogen signaling pathway | | SP1 | ADCY6;CREB1;BCL2;CREB5 | | BCL2 | SP1 | | |
| FoxO signaling pathway | | IL6;FASLG | CDKN1A;FOXO3;FOXO1;CCND2;SMAD4;PTEN;CDKN1B | | | | | |
| Hippo signaling pathway | | BBC3 | CCND2;SNAI2;SMAD4;GSK3B | | | | ID1 | FZD7 |
| Insulin resistance | | IL6 | FOXO1;CREB1;PTEN;GSK3B;CREB5 | | | | NFKBIA | PPARGC1A |
| JAK-STAT signaling pathway | | IL6 | CDKN1A;BCL2;CCND2 | | BCL2 | | | |
| MAPK signaling pathway | | FGFR1;MYD88;FGF21;FASLG | FGF9;BDNF | | | VEGFA;CDC42 | | |
| Neurotrophin signaling pathway | | FASLG | FOXO3;BCL2;GSK3B;BDNF | | BCL2 | NTRK3;CDC42 | NFKBIA | |
| NF-kappa B signaling pathway | | MYD88 | CYLD;BCL2 | | BCL2 | | NFKBIA | |
| Non-alcoholic fatty liver disease | | IL6;FASLG | GSK3B;UQCRFS1 | | | CDC42 | | |
| p53 signaling pathway | | BBC3 | CDKN1A;BCL2;CCND2;PTEN;CHEK2;THBS1 | | BCL2 | | | |
| Parathyroid hormone synthesis, secretion and action | | SP1;FGFR1 | CDKN1A;ADCY6;CREB1;BCL2;CREB5;PLD1 | | BCL2 | SP1 | | |
| PI3K-Akt signaling pathway | | FGFR1;IL6;FGF21;FASLG | CDKN1A;FOXO3;CREB1;FGF9;BCL2;CCND2;PTEN;GSK3B;BDNF;CREB5;CDKN1B;THBS1 | | BCL2 | VEGFA | | |
| Ras signaling pathway | | FGFR1;FGF21;FASLG | FGF9;BDNF;PLD1;TIAM1 | | | VEGFA;CDC42 | | |
| TGF-beta signaling pathway | | SP1 | SMAD4;THBS1 | | | SP1 | ID1 | |
| TNF signaling pathway | | IL6 | CREB1;CREB5 | | | | NFKBIA | |
| AMPK signaling pathway | | | FOXO3;FOXO1;CREB1;CREB5 | | | | | PPARGC1A |
| Apelin signaling pathway | | | ADCY6;SMAD4 | | | | HDAC4 | PPARGC1A |
| Cell cycle | | | CDKN1A;CCND2;SMAD4;GSK3B;CHEK2;CDKN1B | | | E2F1 | WEE1 | |
| cGMP-PKG signaling pathway | | | ADCY6;CREB1;CREB5 | | | | | |
| Circadian rhythm | | | CLOCK;CREB1 | | | | | |
| Endocrine and other factor-regulated calcium reabsorption | | | ADCY6 | | | | | |
| ErbB signaling pathway | | | CDKN1A;GSK3B;CDKN1B | | | | | |
| Ether lipid metabolism | | | PLD1 | | | | | |
| Gastric acid secretion | | | ADCY6 | | | | | |
| Glucagon signaling pathway | | | FOXO1;CREB1;CREB5 | | | | | PPARGC1A |
| Glutamatergic synapse | | | ADCY6;PLD1 | | | | | |
| Glycerophospholipid metabolism | | | PLD1 | | | | | |
| Growth hormone synthesis, secretion and action | | | ADCY6;CREB1;GSK3B;CREB5 | | | | | |
| Hedgehog signaling pathway | | | BCL2;CCND2;GSK3B | | BCL2 | | | |
| Inositol phosphate metabolism | | | PTEN | | | | | |
| Insulin secretion | | | ADCY6;CREB1;CREB5 | | | | | |
| Insulin signaling pathway | | | FOXO1;FLOT1;GSK3B | | | | | FLOT1;PPARGC1A |
| Longevity regulating pathway | | | FOXO3;FOXO1;ADCY6;CREB1;CREB5 | ATG5;RB1CC1 | | | | PPARGC1A |
| mTOR signaling pathway | | | PTEN;GSK3B | | | | | FZD7 |
| Phosphatidylinositol signaling system | | | PTEN | | | | | |
| Phospholipase D signaling pathway | | | ADCY6;PLD1 | | | | | |
| Sphingolipid signaling pathway | | | BCL2;PTEN;PLD1 | | BCL2 | | | |
| Thyroid cancer | | | CDKN1A | | | | | |
| Thyroid hormone synthesis | | | ADCY6;CREB1;CREB5 | | | | | |
| Wnt signaling pathway | | | CCND2;SMAD4;GSK3B | | | | | FZD7 |
| Glycosphingolipid biosynthesis - lacto and neolacto series | | | | FUT4 | | | | |
| Mannose type O-glycan biosynthesis | | | | FUT4 | | | | |
| mRNA surveillance pathway | | | | | | MSI1 | | |
| Adipocytokine signaling pathway | | | | | | | NFKBIA | PPARGC1A |

**Supplementary Table 8: Candidate genes for hsa-mR-182-5p from database entries stratified by metabolic pathways from Gene Ontology. Additional column qPCR contains target genes which were considered for analysis by qPCR in the complete cohort.**

| Name | GO Term | Gene list | qPCR |
|---|---|---|---|
| Insulin receptor signaling pathway | GO:0008286 | AP3S1, APC, APPL1, EIF4EBP2, FOXO1, GRB2, GSK3B, IGF1R, IGF1R, INSR, IRS1, IRS4, PDK4, PHIP, PIK3C2A, PIK3R1, PTPN1, SLC2A8, SOGA1 | FOXO1, INSR, IRS1 |
| Gluconeogenesis | GO:0006094 | ATF3, ATF4, CERS2, CRTC2, G6PC1, GPI, PC, SLC37A4 | G6PC1 (G6PC) |
| Lipid biosynthetic process | GO:0008610 | ACLY, ACSL4, FDFT1, PRKAA1, PRKAA2, SC5D, SCD | SCD |
| Sterol biosynthetic process | GO:0016126 | DHCR7, FDFT1, HMGCS1, INSIG1, LBR, PRKAA1, PRKAA2, SC5D | DHCR7, HMGCS1, INSIG1 |
| Lipid metabolism | GO:0006629 | ABCA2, ABHD12, ACAA2, ACACA, ACER3, ACLY, ACSL4, ADIPOR1, AGPAT2, ALDH1A3, ALDH1A3, ALDH3A2, B3GALT1, B4GALT3, B4GALT5, BCAT2, BCKDHB, BRCA1, CBR4, CDIPT, CERS2, CERS5, CERS6, CHST10, CROT, DDHD1, DDHD2, DGKH, DHCR7, ELOVL6, ETNK2, FAR1, FASN, FDFT1, G6PD, HMGCLL1, HMGCS1, INSIG1, JAZF1, LBR, LIPT1, LPCAT3, LRP10, LRP8, MBLAC2, MGST2, NAA40, NEU3, OCRL, OSBPL10, PAFAH1B1, PAFAH1B1, PAFAH1B2, PC, PCCA, PDE3A, PDSS2, PIGA, PIK3C2A, PIK3CB, PIP5K1A, PLAGL2, PLCB4, PPARD, PPARG, PRDX6, PRKAA1, PRKAA2, PTDSS1, PTGR2, PTGS2, RAB7A, RDH10, RDH11, RNF213, RNF213, SC5D, SCD, SERINC5, SGMS1, SGPL1, SPTSSA, SPTSSB, SRD5A1, ST6GALNAC6, ST8SIA1, ST8SIA1, SYNJ1, TTC39B, UGCG, VLDLR, XBP1, ABCA1, ACER2, ACOT1, ACOT2, ALDH1A1, APOB, BCAT1, GDPD2, GLA, HADHB, MLYCD, OXSM, PAM, PCYT1B, PIK3C2G, PIP5K1B, PLA1A, PLA2G12A, PLA2G3, PNPLA2, PTEN, PTPN22, ST3GAL4, UGT2B17 | ACACA, SCD, PNPLA2, FASN, ELOVL6, DHCR7, HMGCS1, INSIG1 |
| Glycogen synthesis | GO:0005978 | -- | -- |
| Glycogen breakdown | GO:0005980 | G6PC1 | G6PC1 (G6PC) |
| Insulin response | GO:0032868 | CAT, EIF6, GALP, IGF1R, IGF1R, IRS1, KLF15, SESN3, SIRT1, SORT1, SRSF4, SRSF6, TSC1, OTC, SESN2 | IRS1 |
| Lipolysis | GO:0016042 | CRTC3, DDHD1, DDHD2, NEU3, PAFAH1B1, PAFAH1B1, PAFAH1B2, PCCA, PLCB4, PRDX6, RAB7A, TBL1XR1, APOB, PLA1A, PLA2G12A, PNPLA2 | PNPLA2 |
| Lipogenesis | GO:0008610 | ACLY, ACSL4, FDFT1, PRKAA1, PRKAA2, SC5D, SCD | SCD |
| Cholesterol metabolism | GO:0008203 | CAT, CEBPA, DHCR7, FDFT1, HMGCS1, INSIG1, LBR, PPARD, PRKAA1, PRKAA2, TSKU, VLDLR, ABCA1, APOB | DHCR7, HMGCS1, INSIG1 |
| Canonical Wnt signaling pathway | GO:0060070 | BCL9L, CSNK1E, CTDNEP1, EDNRB, FGF2, FGF9, FOXO1, FOXO3, FZD3, FZD6, ISL1, PYGO2, RARG, RECK, SIAH2, WNT5A, YAP1, LRP6, PTEN | FOXO1, LRP6 |

Abbrevations (only genes of interest listed in "qPCR"): 3'UTR, 3' untranslated region; bp, base pairs; ACACA, Acetyl-CoA Carboxylase Alpha; DHCR/, 7-Dehydrocholesterol Reductase; ELOVL6, ELOVL Fatty Acid Elongase 6; FASN, Fatty Acid Synthase; FOXO1, Forkhead box protein O1; G6PC, Glucose-6-phosphatase, catalytic; HMGCS1, 3-Hydroxy-3-Methylglutaryl-CoA Synthase 1; INSIG1, insulin-induced gene 1; INSR, insulin receptor; IRS1, Insulin receptor substrate 1; LRP6, Low-density lipoprotein receptor-related protein 6; PCK1, Phosphoenolpyruvate carboxykinase 1; PDK1, Phosphoinositide-dependent kinase-1; PNPLA2, Patatin Like Phospholipase Domain Containing 2; PPARA, Peroxisome Proliferator Activated Receptor Alpha; SCD, Stearoyl-CoA Desaturase;

**Supplementary Table 9: Result of all consulted target gene databases and a comprehensive 3'UTR screening for each potential target gene of hsa-miR-182-5p. Besides the relative position of the seed sequences within the 3'-UTR of the target gene, also the relative AU content of the surrounding 60 nt, whether there is an additional base pairing between the mRNA and the miRNA 3' of the seed and the relative position of the seed within the target 3'UTR is listed.**

| target_gene | database prediction | seed match in 3'UTR | validated (positive experiment) | Conserved_miRNA_TarScan_v7 | | TargetScan_v7 | | TarBase_v7 | | TarBase_v8 | | | miRTarBase_MTI | miRDB_v6 | Affymetrix (databases microcosm; MTI; | |
|---|---|---|---|---|---|---|---|---|---|---|---|---|---|---|---|---|
| | | | | seed | score | seed | score | cell line/tissue | method | cell line/tissue | method | score | method | target score | target genes | validated |
| ***SCD*** | yes | yes | yes | -- | -- | -- | | MCF7/mammary gland; TZMBL/cervix | immunoprecipitation | see TarBase_v7 | | | -- | -- | -- | -- |
| ***FASN*** | yes | yes | yes | 7mer_A1 | 0.00 | -- | -- | HELA/cervix | immunoprecipitation | see TarBase_v7 | | | -- | -- | -- | -- |
| ***ELOVL6*** | yes | yes | yes | 8mer; 7mer_m8 | 0.00; 0.02 | -- | -- | HEK293T/kidney; MCF/mammary gland | correlation with microarray data; immunoprecipitation | Liver (Melanoma metastatic Liver Cells ) | correlation with microarray data | negative | -- | -- | -- | -- |
| ***ACACA*** | yes | no | yes | -- | -- | -- | -- | HEK293T/kidney | correlation with microarray data; immunoprecipitation | see TarBase_v7 | | | -- | -- | -- | -- |
| ***PNPLA2*** | yes | yes | no | 7mer_m8 | 0.00 | -- | -- | -- | -- | -- | -- | -- | -- | -- | yes | -- |
| ***PPARA*** | yes | yes | no | 7mer_m8; 7mer_A1 | 0.05; 0.00 | -- | -- | -- | -- | -- | -- | -- | -- | -- | -- | -- |
| ***LRP6*** | yes | yes | no | 7mer_m8 | 0.00 | -- | -- | -- | -- | Liver (Melanoma metastatic | correlation with microarray | negative | -- | -- | yes | -- |
| ***PDK1*** | yes | yes | no | 7mer_A1; 7mer_m8 | 0.00; 0.02 | -- | -- | -- | -- | -- | -- | -- | -- | -- | -- | -- |
| ***INSR*** | yes | yes | no | 8mer | 0.00 | -- | -- | -- | -- | -- | -- | -- | -- | 57 | -- | -- |
| ***IRS1*** | yes | yes | yes | 7mer_m8 | 0.15 | -- | -- | MCF7/mammary gland; TZMBL/cervix | immunoprecipitation | MCF7/mammary gland; TZMBL/cervix | immunoprecipitation | 0.68 | -- | -- | -- | -- |
| ***FOXO1*** | yes | yes | yes | 7mer_A1 | 0.72 | 7mer_A1 | 0.72 | HEK293T/kidney; MCF7/mammary gland; | correlation with microarray data, Luciferase Reporter Assay, Western Blot, qPCR | see TarBase_v7 | | | Luciferase Reporter Assay, | 75 | yes | yes |
| ***G6PC*** | yes | yes | no | 7mer_m8 | 0.01 | -- | -- | -- | -- | -- | -- | -- | -- | -- | -- | -- |
| ***PCK1*** | no | yes | no | -- | -- | -- | -- | -- | -- | -- | -- | -- | -- | -- | -- | -- |
| ***DHCR7*** | yes | yes | yes | -- | -- | -- | -- | TZMBL/cervix | immunoprecipitation | see TarBase_v7 | | | -- | | -- | -- |
| ***HMGCS1*** | yes | yes | yes | 7mer_A1 | 0.00 | -- | -- | TZMBL/cervix | immunoprecipitation | see TarBase_v7 | | 0.51 | -- | | -- | -- |
| ***INISG1*** | yes | yes | no | 7mer_m8 | 0.73 | 7mer_m8 | 0.73 | -- | -- | -- | -- | -- | -- | 90 | -- | -- |

| target_gene | regulator | Position_8mer | Position_7mer_m8 | Position_7mer_A1 | Position_6mer | Position_offset_6mer | AU_content_within_60nt | additional_3'_mRNA_binding | relative_within_UTR |
|---|---|---|---|---|---|---|---|---|---|
| | mature miRNA | 3' UTR bp | 3' UTR bp | 3' UTR bp | 3' UTR bp | 3' UTR bp | fraction | position - bases | |
| ***SCD*** | hsa-miR-182-5p | 0 | 0 | 0 | 1186 | 0 | 0.514705882 | 0 - 0 | 0.304649371 |
| | hsa-miR-182-5p | 0 | 0 | 0 | 0 | 1293 | 0.529411765 | 0 - 0 | 0.332134601 |
| | hsa-miR-182-5p | 0 | 0 | 0 | 0 | 3707 | 0.617647059 | 0 - 0 | 0.952221937 |
| ***FASN*** | hsa-miR-182-5p | 0 | 0 | 0 | 366 | 0 | 0.323529412 | 0 - 0 | 0.454658385 |
| ***ELOVL6*** | hsa-miR-182-5p | 657 | 0 | 0 | 0 | 0 | 0.529411765 | 0 - 0 | 0.119628551 |
| | hsa-miR-182-5p | 0 | 4419 | 0 | 0 | 0 | 0.529411765 | 0 - 0 | 0.804624909 |
| | hsa-miR-182-5p | 0 | 4535 | 0 | 0 | 0 | 0.5 | 0 - 0 | 0.82574654 |
| ***ACACA*** | hsa-miR-182-5p | -- | -- | -- | -- | -- | -- | -- | -- |
| ***PNPLA2*** | hsa-miR-182-5p | 0 | 76 | 0 | 0 | 0 | 0.352941176 | 0 - 0 | 0.106891702 |
| ***PPARA*** | hsa-miR-182-5p | 0 | 5335 | 0 | 0 | 0 | 0.558823529 | 0 - 0 | 0.637014925 |
| | hsa-miR-182-5p | 0 | 6414 | 0 | 0 | 0 | 0.544117647 | 0 - 0 | 0.765850746 |
| | hsa-miR-182-5p | 0 | 0 | 3202 | 0 | 0 | 0.602941176 | 0 - 0 | 0.382328358 |
| | hsa-miR-182-5p | 0 | 0 | 0 | 2483 | 0 | 0.382352941 | 0 - 0 | 0.296477612 |
| | hsa-miR-182-5p | 0 | 0 | 0 | 0 | 2886 | 0.661764706 | 0 - 0 | 0.344597015 |
| | hsa-miR-182-5p | 0 | 0 | 0 | 0 | 6209 | 0.367647059 | 0 - 0 | 0.741373134 |
| ***LRP6*** | hsa-miR-182-5p | 0 | 1788 | 0 | 0 | 0 | 0.632352941 | 1781 - 5 | 0.350519506 |
| | hsa-miR-182-5p | 0 | 0 | 0 | 2332 | 0 | 0.661764706 | 0 - 0 | 0.457165262 |
| | hsa-miR-182-5p | 0 | 0 | 0 | 0 | 1644 | 0.558823529 | 0 - 0 | 0.322289747 |
| ***PDK1*** | hsa-miR-182-5p | 0 | 4672 | 0 | 0 | 0 | 0.573529412 | 0 - 0 | 0.367874016 |
| | hsa-miR-182-5p | 0 | 0 | 618 | 0 | 0 | 0.602941176 | 0 - 0 | 0.048661417 |
| | hsa-miR-182-5p | 0 | 0 | 0 | 2618 | 0 | 0.632352941 | 0 - 0 | 0.206141732 |
| | hsa-miR-182-5p | 0 | 0 | 0 | 11601 | 0 | 0.544117647 | 0 - 0 | 0.913464567 |
| | hsa-miR-182-5p | 0 | 0 | 0 | 0 | 1839 | 0.691176471 | 0 - 0 | 0.14480315 |
| | hsa-miR-182-5p | 0 | 0 | 0 | 0 | 4099 | 0.735294118 | 0 - 0 | 0.322755906 |
| | hsa-miR-182-5p | 0 | 0 | 0 | 0 | 4207 | 0.661764706 | 0 - 0 | 0.331259843 |
| | hsa-miR-182-5p | 0 | 0 | 0 | 0 | 6622 | 0.529411765 | 0 - 0 | 0.521417323 |
| | hsa-miR-182-5p | 0 | 0 | 0 | 0 | 10591 | 0.573529412 | 0 - 0 | 0.833937008 |
| | hsa-miR-182-5p | 0 | 0 | 0 | 0 | 11494 | 0.544117647 | 0 - 0 | 0.90503937 |
| ***INSR*** | hsa-miR-182-5p | 2093 | 0 | 0 | 0 | 0 | 0.691176471 | 0 - 0 | 0.436860781 |
| ***IRS1*** | hsa-miR-182-5p | 0 | 3873 | 0 | 0 | 0 | 0.558823529 | 0 - 0 | 0.780846774 |
| | hsa-miR-182-5p | 0 | 0 | 0 | 210 | 0 | 0.617647059 | 199 - 6;200 - 5 | 0.04233871 |
| | hsa-miR-182-5p | 0 | 0 | 0 | 4932 | 0 | 0.644067797 | 0 - 0 | 0.994354839 |
| | hsa-miR-182-5p | 0 | 0 | 0 | 0 | 4255 | 0.632352941 | 0 - 0 | 0.857862903 |
| ***FOXO1*** | hsa-miR-182-5p | 0 | 0 | 265 | 0 | 0 | 0.661764706 | 0 - 0 | 0.078356002 |
| | hsa-miR-182-5p | 0 | 0 | 2140 | 0 | 0 | 0.588235294 | 0 - 0 | 0.632761679 |
| | hsa-miR-182-5p | 0 | 0 | 0 | 0 | 557 | 0.529411765 | 0 - 0 | 0.164695446 |
| ***G6PC*** | hsa-miR-182-5p | 0 | 0 | 0 | 697 | 0 | 0.647058824 | 0 - 0 | 0.231407703 |
| ***PCK1*** | hsa-miR-182-5p | 0 | 0 | 0 | 1172 | 0 | 0.426470588 | 0 - 0 | 0.5058265 |
| | hsa-miR-182-5p | 0 | 0 | 0 | 0 | 1591 | 0.455882353 | 0 - 0 | 0.686663789 |
| | hsa-miR-182-5p | 0 | 0 | 0 | 0 | 2095 | 0.397058824 | 0 - 0 | 0.904186448 |
| ***DHCR7*** | hsa-miR-182-5p | 0 | 0 | 0 | 0 | 213 | 0.485294118 | 0 - 0 | 0.22327044 |
| ***HMGCS1*** | hsa-miR-182-5p | 0 | 0 | 214 | 0 | 0 | 0.544117647 | 0 - 0 | 0.058453974 |
| ***INISG1*** | hsa-miR-182-5p | 0 | 1780 | 0 | 0 | 0 | 0.735294118 | 0 - 0 | 0.905852417 |
| | hsa-miR-182-5p | 0 | 0 | 0 | 0 | 311 | 0.691176471 | 0 - 0 | 0.15826972 |
| | hsa-miR-182-5p | 0 | 0 | 0 | 0 | 849 | 0.602941176 | 0 - 0 | 0.432061069 |
| | hsa-miR-182-5p | 0 | 0 | 0 | 0 | 1780 | 0.735294118 | 0 - 0 | 0.905852417 |

Abbrevations: 3'UTR, 3' untranslated region; bp, base pairs; ACACA, Acetyl-CoA Carboxylase Alpha; DHCR/, 7-Dehydrocholesterol Reductase; ELOVL6, ELOVL Fatty Acid Elongase 6; FASN, Fatty Acid Synthase; FOXO1, Forkhead box protein O1; G6PC, Glucose-6-phosphatase, catalytic; HMGCS1, 3-Hydroxy-3-Methylglutaryl-CoA Synthase 1; INSIG1, insulin-induced gene 1; INSR, insulin receptor; IRS1, Insulin receptor substrate 1; LRP6, Low-density lipoprotein receptor-related protein 6; PCK1, Phosphoenolpyruvate carboxykinase 1; PDK1, Phosphoinositide-dependent kinase-1; PNPLA2, Patatin Like Phospholipase Domain Containing 2; PPARA, Peroxisome Proliferator Activated Receptor Alpha; SCD, Stearoyl-CoA Desaturase;
